# Single individual structural variant detection uncovers widespread hemizygosity in molluscs

**DOI:** 10.1101/2020.09.15.298695

**Authors:** Andrew D Calcino, Nathan J Kenny, Marco Gerdol

## Abstract

The advent of complete genomic sequencing has opened a window into genomic phenomena obscured by fragmented assemblies. A good example of these is the existence of hemizygous regions of autosomal chromosomes, which can result in marked differences in gene content between individuals within species. While these hemizygous regions, and presence/absence variation of genes that can result, are well known in plants, firm evidence has only recently emerged for their existence in metazoans.

Here we use recently published, complete genomes from wild-caught molluscs to investigate the prevalence of hemizygosity and pan-genomes across a well-known and ecologically important clade. We show that hemizygous regions are widespread in mollusc genomes, not clustered in individual chromosomes, and often contain genes linked to transposition, DNA repair and stress response. With targeted investigations of *HSP70-12* and *C1qDC*, we also show how individual gene families are distributed within pan-genomes.

This work suggests that pan-genomes are widespread across the conchiferan Mollusca, and represent useful tools for genomic evolution, allowing the maintenance of additional genetic diversity within the population. As genomic sequencing and re-sequencing becomes more routine, the prevalence of hemizygosity, and its impact on selection and adaptation, are key targets for research across the tree of life.

## Background

The rapid development of third generation sequencing technologies and the subsequent increase in the number of species with an assembled genome has led to extraordinary new insights into gene family evolution between related species. At the same time, the flood of new genomic information has given rise to new questions regarding the dynamics of gene family expansion and contraction, and the mechanisms that could be driving such potentially adaptive processes in disparate taxa.

With increasing depth of sequencing data, it has become apparent that not all individuals of a species possess structurally identical genomes. While small-scale variation (single nucleotide polymorphisms, indels, duplications, inversions and translocations) between individuals was entirely expected, large-scale structural variations, incorporating gene content disparity, have come as somewhat of a surprise in metazoan lineages. Variation in genic content between individuals of a species forms the basis of the pan-genome concept [1]. First described in bacteria, a pan-genome consists of a core set of genes shared by all members of a species in addition to a set of ‘dispensable’ genes which are subject to presence/absence variation (PAV) between individuals. In the context of a diploid eukaryote, dispensable genes may be present in either one, two or zero copies in any given individual, meaning that an assembled genome of a single individual of a pan-genomic species may not capture the full genic complement of that species.

In plants, two recent papers on the pan-genomes of rapeseed and tomato have highlighted the extensive PAV in these species and also the functional importance of some of their dispensable genes [2,3]. In total, at least 12 plant pan-genomes have now been described (reviewed in [4]). In plants, most SVs are associated with transposons and are rich in repeat sequences.

In metazoans, information on pan-genomes is only starting to emerge with genomic variation between individuals noted in animals such as the roundworm *Caenorhabditis brenneri [5]*, the family *Caenorhabditis* more generally [6], and the ascidian *Ciona savignyi [7]*.

Analyses of humans, probably the most re-sequenced metazoan species, are also beginning to show evidence of a closed pan-genome (i.e. a pan-genome with a low rate of dispensable to core genes). Based on the resequencing of 2504 human genomes, a total of 240 genes were found to be occasionally subject to homozygous deletions in healthy individuals (and therefore likely dispensable) [8]. Analysis of the genome from 910 individuals of African descent led to the assembly of 296 Mb of genomic sequence not included in the reference human genome [9]. In pigs, the size of the pig pan-genome, estimated based on the resequencing of 12 individuals, revealed the presence of 72.5 Mb additional genomic sequence [10]. High levels of genomic heterozygosity and gene presence/absence have been recorded in the bivalve molluscs *Mytilus galloprovincialis* [11] and *Crassostrea gigas* [12]. In *M. galloprovincialis*, a species characterized by an open pan-genome (i.e. by a 1:3 dispensable to core genes rate) this has been firmly linked to hemizygosity [11], however, it is unknown how widely this trait is shared with other animals, or with other molluscs in particular.

Hemizygosity occurs in a genome where only one of the two chromosomal pairs encodes a region or block of DNA. In mammals the most prominent example of hemizygous DNA comes from the X chromosome when it occurs in males. Under the male condition, no homologous region for the majority of the X chromosome exists on the Y chromosome which leaves the genes encoded by these regions monoallelic. To cope with this, dosage compensation mechanisms have evolved to alleviate the issues associated with reduced transcriptional output in males [13]. Hemizygous regions can also result from insertions or deletions (indels) of blocks of DNA and can occur through potentially pathological pathways (ie. retrovirus or transposon insertion, eg. [14]). The detection of hemizygous DNA that encodes genes in a single individual is evidence that at the population level, these genes are likely to be subject to presence/absence variation as individuals may possess one (hemizygous), two (homozygous) or zero copies (nullizygous) of the dispensable DNA block.

Questions arising from the increasing observation of intraspecific genomic structural variation include, how representative are the single genomes we have for most species of the population or species from which they derive? Are the gene family contractions and expansions observed in these representative genomes shared in their entirety by all other members of the species? If not, what are the implications for species-wide phenotypic variability, gametic compatibility and ecological adaptability? While some of these questions will not be resolved until data from multiple individuals per species, spanning multiple phyla are available, targeted analysis of individual high quality genomes can be utilised to determine how prevalent genomic structural variability and pan-genomes may be within particular clades.

Molluscan species, which are commonly profligate broadcast spawners [15], are often found inhabiting quite variable environments [15]. Many aquatic species, particularly bivalves, are sessile filter feeders that occur in high-density beds. This places them at risk of infection and exposure to locally unfavourable conditions [16]. The maintenance of genetic variation through a pan-genome therefore could enable a population to adapt to changes in environment or extreme local conditions [17].

Here we examine the hemizygous DNA complement of eight high-quality and near complete conchiferan mollusc genomes (Fig. 1a). We detail the impacts of widespread hemizygosity on the genomic architectures of this species-rich and ecologically important clade, and investigate the hemizygous gene complement of each species. We note that retroelement-related genes and those involved in splice repair are over-represented in our datasets. Similarly, HSP70, C1qDC proteins, C-type lectins and immune-related GTPases are common in hemizygous regions, suggesting possible adaptive roles in stress response and immunity. Our method, using freely available public data, could be applied to any well-sampled clade. It will be broadly applicable across the tree of life as more high quality genome sequences become available, providing a clear means of investigating this under-studied, but potentially widespread, means of genomic adaptation.

**Figure 1.**
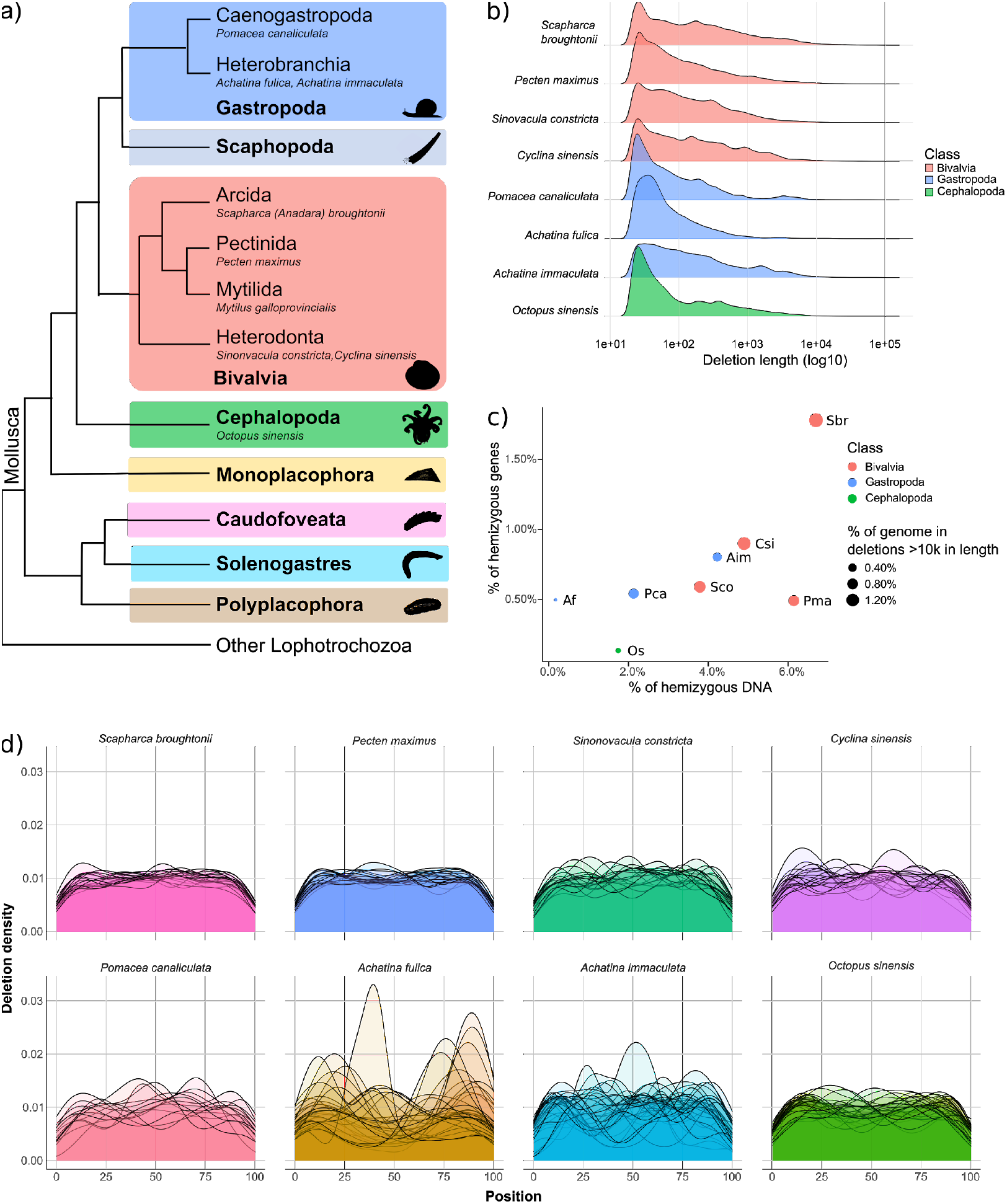
Phylogeny and hemizygous loci analyses of eight molluscan species. a) Representative cladogram of mollusc relationships after [77]. Species referenced in this manuscript shown in italics. Note, in Bivalvia and Gastropoda, numerous subclades are not shown. b) Length distribution of hemizygous regions (deletions). c) Percentage of each genome which is hemizygous versus the percentage of all genes which reside entirely within hemizygous DNA. The size of each point is proportional to the percentage of the genome found in large (>10 kb) hemizygous regions. Species include *Achatina fulica* (Af), *Achatina immaculata* (Aim), *Cyclina sinensis* (Csi), *Octopus sinensis* (Os), *Pecten maximus* (Pma), *Pomacea canaliculata* (Pca), *Scapharca broughtonii* (Sbr) and *Sinonovacula constricta* (Sco). d) Density of hemizygous loci for each species. Each chromosome is represented by an individual data series (line) which spans the beginning (0% distance) to the end (100% distance) of each chromosome.

## Methods

Genome assemblies and datasets used for all subsequent analyses are provided in Table 1. Details on command line options for each step in the analysis pipeline can be found in Supplementary File 1. A brief description of each step involved is provided here.

**Table 1:** Genomes used, and basic statistics regarding assemblies.

| Genome | Abbreviation | Bioproject | Heterozygosity | Genome size | % hemizygosity |
| --- | --- | --- | --- | --- | --- |
| *Pecten maximus* | Pma | PRJEB22206 | 1.69% | 918,306,378 | 6.14% |
| *Sinonovacula constricta* | Sco | PRJNA508451 | 2.42% | 1,220,848,272 | 3.78% |
| *Cyclina sinensis* | Csi | PRJNA612143 | 1.11% | 903,119,975 | 4.89% |
| *Scapharca (Anadara) broughtonii* | Sbr | PRJNA521075 | 1.79% | 884,566,040 | 6.69% |
| *Pomacea canaliculata* | Pca | PRJNA427478 | 1.92% | 440,159,624 | 2.12% |
| *Achatina (=Lissachatina) fulica* | Afu | PRJNA511624 | 0.19% | 1,855,892,613 | 0.17% |
| *Achatina (=Lissachatina) immaculata* | Aim | PRJNA561271 | 0.74% | 1,653,153,977 | 4.22% |
| *Octopus sinensis* | Osi | PRJNA541812 |  | 2,719,136,158 | 1.74% |

### Species selection

To document hemizygosity across conchiferan molluscs, we selected publicly available genomes that met three criteria. 1) The available assembly of each genome should be at or approaching chromosomal level, 2) the individual sequenced should be wild caught and 3) there should be publicly available PacBio and Illumina datasets available that were produced from the same individual from which the assembly was generated. These features were sought so that analyses of chromosomal distribution could be performed, artefacts from captive breeding could be avoided, so that precise boundaries of hemizygous regions could be determined and so that *k*-*mer* coverage of hemizygous regions could be performed. Hi-C Illumina short read datasets were avoided as they do not map uniformly to their respective genomes.

### Structural variant detection

To identify hemizygous regions, PacBio’s structural variant detection pipeline pbsv v2.2.2 [18] was run on each of the eight genomes. The mapped long read bam file and tandem repeats annotation files required for pbsv were generated with pbmm2 v1.0.0 [19] and Tandem Repeats Finder v4.09 [20] respectively. As the aim was to identify regions of the genome assemblies that were both hemizygous and contained previously annotated genes, the type of structural variants detected by pbsv that were used for subsequent analyses were limited to deletions. Detected deletions that passed pbsv’s default criteria were filtered for further analysis.

To visualise the level of hemizygosity in each species, deletions at least 10kb in length that were not associated with tandem repeats were extracted and used as input for chromoMap v0.2 [21].

### *K* mer analysis of hemizygous regions

Illumina short read libraries were preprocessed with bbduk [22] and mapped to their respective genomes with bwa mem v0.7.16 [23]. In cases where more than one dataset was used per species, mapped bam files were merged with samtools merge [24]. Mapped reads were extracted with samtools view and converted to fasta format with bbtools reformat.sh [22] and *k*-mer histograms (*k*=21) of all mapped reads were produced with Jellyfish v2.3.0 [25]. This histogram was uploaded to Genomescope [26] to obtain heterozygosity estimates for each species.

To produce histograms of *k*-mers from reads mapped to hemizygous regions, reads mapping to hemizygous regions were extracted with samtools view after which only those falling entirely within the defined regions were filtered with bedops bedmap v2.4.37 [27]. A fasta file of all filtered reads was extracted with bedtools fastaFromBed v2.29.0 [28] and these were used to produce *k*-mer histograms with Jellyfish as was performed for the whole mapped library.

### Read coverage of hemizygous regions

The same mapped reads used for *k*-mer coverage were also used to determine read coverage of hemizygous regions and to compare to read coverage of the whole genomes. A sam file of all reads mapping entirely within deletions was produced and from this a bam file was produced with samtools view. The median coverage of each deletion was calculated with mosdepth v0.2.9 [29]. For the whole genome, bedtools genomecov v2.29.0 [28] was used to calculate coverage at every position in the genome and then the median coverage of every 1000 nt (1 nt step) window was calculated.

### Hemizygous gene identification

Genes were extracted from hemizygous regions of the genome using bedtools intersect v2.29.2 [28]. Only those genes falling fully within hemizygous regions were extracted for analysis (using the -F 1 option as detailed in Supplementary File 1). From these lists, genes were identified and extracted from the full gene list for use in enrichment analyses.

### Protein domain and Gene Ontology enrichment analysis

The predicted protein translations obtained from the longest annotated isoform for each gene in the target molluscan genomes were functionally annotated with Pfam conserved domains ID and Gene Ontology terms as follows. Amino acid sequences were subject to a BLASTp analysis against UniProtKB, with an *E*-value threshold set to 1E^−5^. Gene Ontology cellular component, biological process and molecular functions terms associated with the top 10 best hits were extracted and used to annotate matching query sequences. Protein sequences were also subject to conserved domain annotation. This analysis used the hmmscan module of HMMER v3.3.1 [30] and the search was conducted against the Pfam-A v. 33.1 database [31], annotating domains based on the default *E*-value threshold.

The subset of sequences associated with hemizygous regions, identified as described in the previous paragraph, were then subject to hypergeometric tests on annotations [32] using the script included in the SciPy 1.5.2 package, which were run separately for GO terms and Pfam domain IDs. We here report significantly enriched GO terms and Pfam domains, filtering out based on their over-representation in the tested subset of sequences, compared with the full genome. Namely, enriched annotations were reported for terms associated with p-values < 0.05 and a difference between the number of observed and expected genes >= 5.

### Phylogenetic analysis

HSP70 and C1qDC genes were identified within the genomes of the target species using genes of known homology for local BlastP searches (*E*-value cutoff, initially *E*^−9^) [33]. These were then reciprocally blasted against the nr database to confirm likely identity. These sequences, alongside known sequences from previous publications, were aligned using the MAFFT 7 online tool and the G-INS-i strategy [34,35]. The resulting alignments were trimmed with TrimAL v1.2 [36] and the “-gappyout” setting.

The resulting alignments were tested for model fit using ModelFinder [37] as integrated in IQ-TREE multicore version 1.6.10 [38]. The best-fit model was used for analysis of each phylogeny as noted in the Fig. 4 legend. IQ-TREE multicore version 1.6.10 was used for maximum likelihood (ML) analysis with 1000 non-parametric bootstrap replicates. The resulting consensus phylogeny was then opened in FigTree v1.4.4 (https://github.com/rambaut/figtree/releases) for annotation and display.

## Results

### Hemizygosity and heterozygosity in molluscs

Hemizygosity (flagged as deletions by pbsv) of the individual representatives of the eight molluscan species investigated here ranged from 0.17% of the total genome length in the giant african snail *Achatina (=Lissachatina, [39,40]*) *fulica* to 6.69% in the ark clam *Scapharca broughtonii* (Table 1). The *A. fulica* hemizygosity content was a clear outlier amongst the eight species with the next lowest belonging to the octopus *Octopus sinensis* at 1.74%, while the congeneric *Achatina (=Lissachatina) immaculata* had 4.22% hemizygous DNA content. The number and size (bp) of these regions is shown further in Table 2.

**Table 2:** Statistics regarding putatively hemizygous regions.

| Genome | # of deletions | Length of deletions | # of non-TANDEM deletions | Length of non-tandem dels | # of >10k deletions | Length of >10k deletions | % hemizygous genes |
| --- | --- | --- | --- | --- | --- | --- | --- |
| *Pecten maximus* | 145,503 | 56,374,042 | 60,503 | 35,699,112 | 428 | 7,359,495 | 0.50% |
| *Sinonovacula constricta* | 108,006 | 46,151,533 | 56,831 | 34,057,642 | 410 | 12,089,568 | 0.60% |
| *Cyclina sinensis* | 75,880 | 44,129,845 | 46,216 | 35,875,686 | 456 | 11,645,828 | 0.91% |
| *Scapharca (Anadara) broughtonii* | 83,518 | 59,203,570 | 47,022 | 47,290,026 | 671 | 13,984,916 | 1.79% |
| *Pomacea canaliculata* | 24,569 | 9,343,212 | 15,080 | 8,489,934 | 111 | 2,830,553 | 0.55% |
| *Achatina (=Lissachatina) fulica* | 16,208 | 3,197,224 | 3,112 | 1,330,508 | 38 | 789,841 | 0.50% |
| *Achatina (=Lissachatina) immaculata* | 128,736 | 69,768,216 | 27,939 | 41,999,937 | 351 | 7,946,998 | 0.81% |
| *Octopus sinensis* | 153,503 | 47,178,568 | 23,187 | 24,977,873 | 131 | 2,440,767 | 0.14% |

Repetitive sequences appear to be a major component of hemizygous DNA in these species with between 39% and 85% of deletions being flagged as tandem duplication-containing by pbsv (Table 2). While the length of most hemizygous regions are short, we observed between 38 and 671 hemizygous regions that exceeded 10kb in length (Fig. 1b,c). Due to the limitations of pbsv, which does not annotate deletions above 100kb in length, the maximum deletion size remains unknown for any of the eight samples. This limitation also means that the total number of deletions and total hemizygous DNA content of each sample are both likely under-estimations.

Previous work on human structural variation showed that the number of SVs was non-randomly distributed along each chromosome with the greatest density occurring within 5 Mb of the telomeric chromosomal ends [41]. The pbsv pipeline’s conservative approach to annotating SVs located towards the ends of chromosomes means that putative terminal SVs are not flagged as a PASS and as such were not included for further analysis here (see methods). This results in the appearance that hemizygosity is diminished at the terminal regions of chromosomes and highlights the fact that the numbers of hemizygous regions reported here are conservative lower estimates (Fig. 1d).

Focusing on deletions over 10 kb in length and which were not flagged as tandem-repeat associated by pbsv, it is evident that large hemizygous regions are not confined to particular chromosomes or chromosomal regions in any of the eight species investigated (Fig. 2). Although there are a large number of hemizygous regions in all species, there is a clear difference in the number of larger deletions present in the bivalves versus the gastropods and cephalopods (Fig. 1b, 2). This seems to also translate into the proportion of the genomes that are hemizygous in each of the three molluscan classes however more species will need to be analysed before these trends can be confirmed (Fig. 1c).

**Figure 2:**
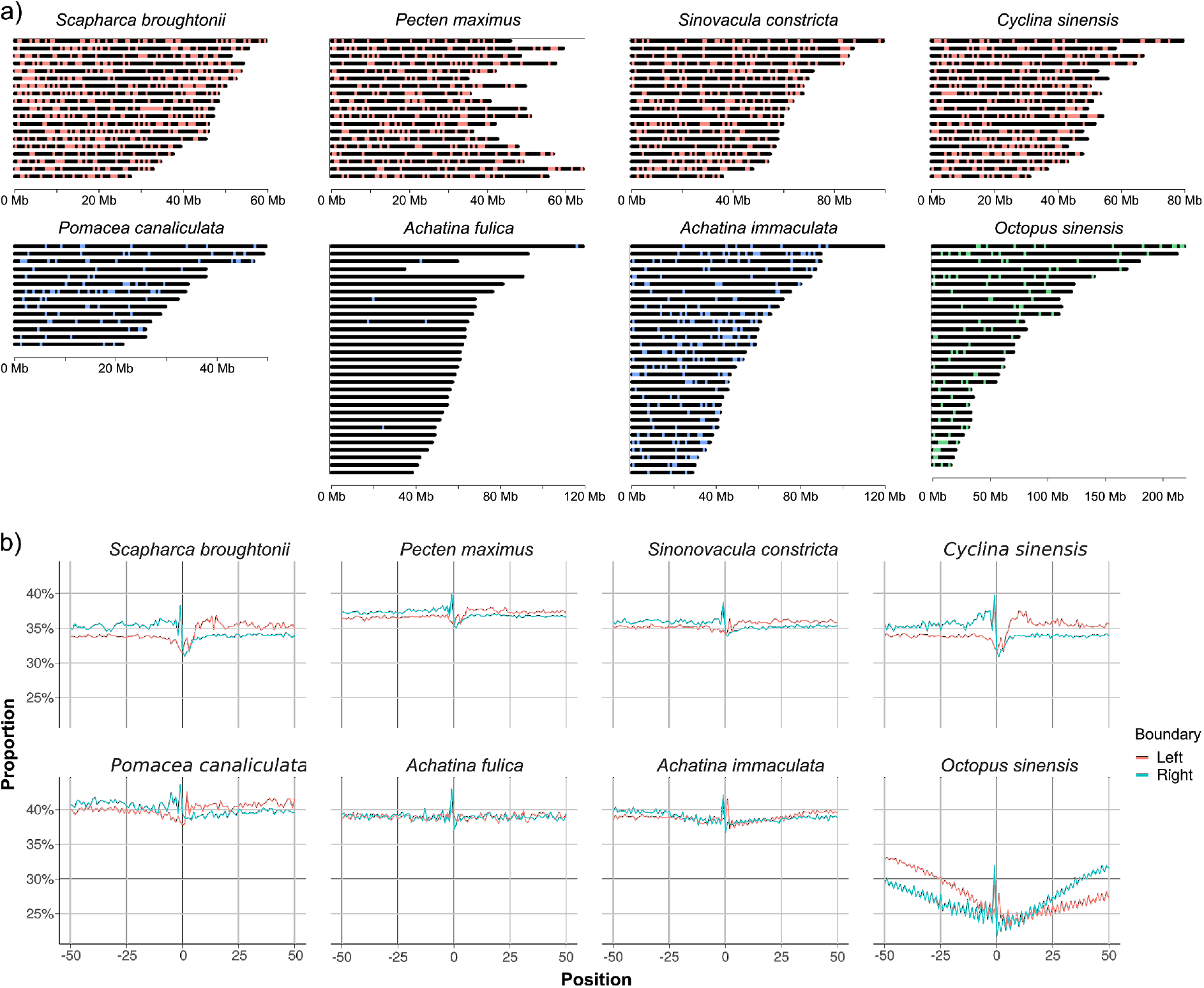
Chromosomal maps of hemizygous loci and G/C content across homozygous/hemizygous boundaries. a) Hemizygous loci greater than 10 kb in length which were not flagged by pbsv as ‘tandem repeats’. Each locus is marked as a single point which is not proportional in length to the actual size of the locus. Genomes with red loci are bivalves, those with blue loci are gastropods and the genome with green loci is a cephalopod. b) Average G/C content spanning 50 bp downstream and 50 bp upstream of the left homozygous/hemizygous boundary or 50 bp downstream and 50 bp upstream of the right homozygous/hemizygous boundary for all annotated hemizygous loci. In each species the transition between homozygous and hemizygous DNA is marked by a G/C spike and apart from the octopus, hemizygous DNA is generally more G/C rich than the flanking homozygous regions. For octopus, hemizygous loci are more A/T rich than the flanking homozygous regions and the entire region surrounding the boundary is relatively depleted of G/C nucleotides.

Heterozygosity of each sample as determined by Genomescope [26] ranged from 0.19% in *A. fulica* to 2.42% in the razor clam *Sinonovacula constricta*. Heterozygosity of the octopus was not able to be determined as the model failed to converge (Supplementary File 2), however visual inspection of the *k*-mer histogram suggests that it is likely to be low and second only to *A. fulica* (Fig. 3, see below), consistent with a previous report from the congeneric species *Octopus bimaculoides* [42].

**Figure 3:**
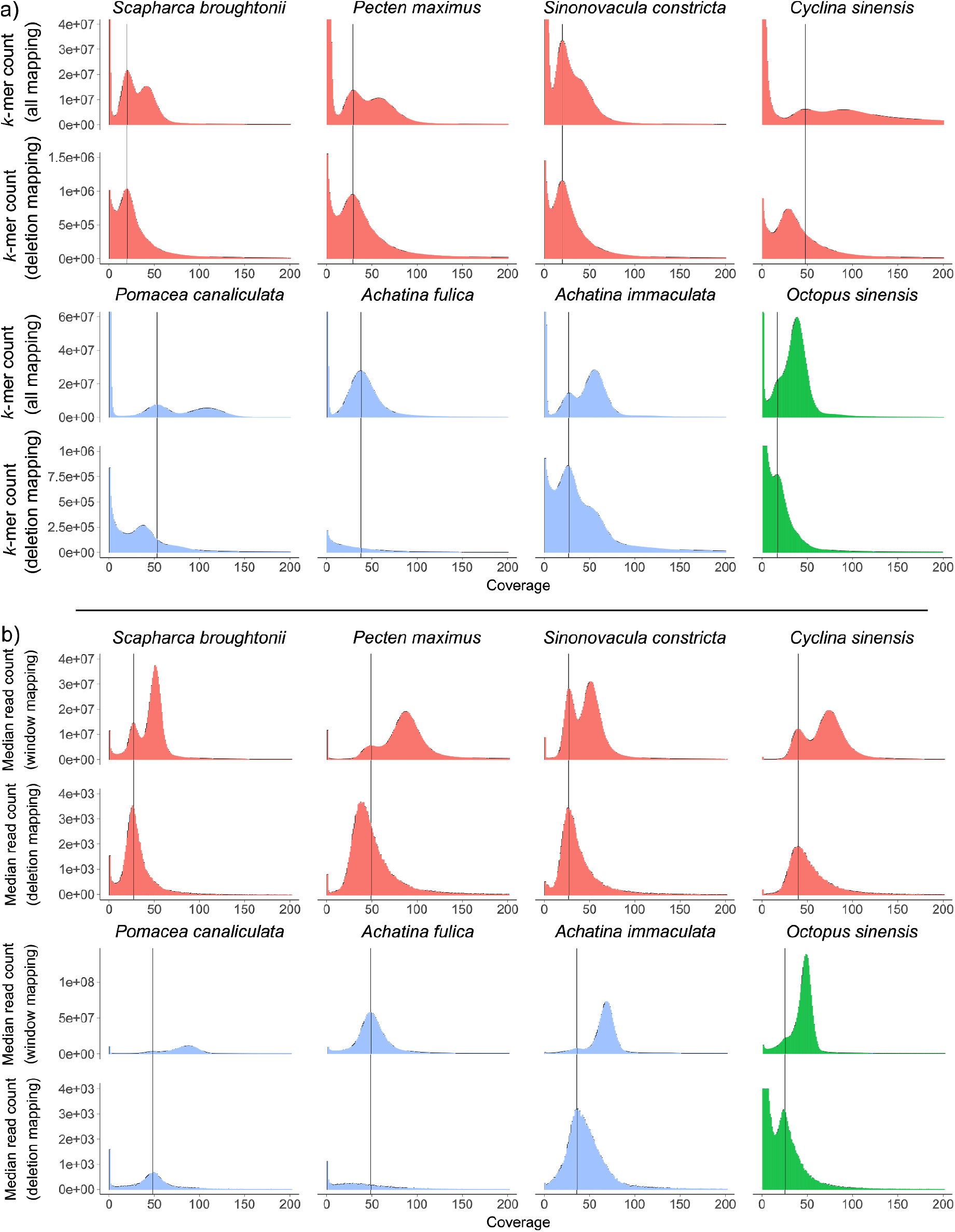
*k*-mer and median read coverage analysis of hemizygous regions. a) *k*-mer counts of all mapped reads for each genome with the corresponding *k*-*mer* counts of reads that map entirely within hemizygous regions located directly below. b) Median read coverage of all 1000 bp sliding windows for each genome with the corresponding median read coverage of all annotated hemizygous regions located directly below. For both a) and b) the black vertical lines mark the ‘heterozygous’ peaks of the total mapped reads *k*-*mer* or median read coverage plots. Species are colour coded by class with red for bivalves, blue for gastropods and green for the cephalopod.

### *K* mer and read coverage of hemizygous regions

Both *k*-mer coverage and and read coverage of the putative hemizygous regions provide strong support that these regions are in fact hemizygous. In six of the eight species, two clear peaks in both *k*-mer coverage (Fig. 3a, top row) and read coverage (Fig. 3b, top row) correspond to what are typically described as the heterozygous and homozygous peaks. The remaining two species, *A. fulica* and *O. sinensis*, have relatively low levels of both heterozygosity and hemizygosity and accordingly only have a single homozygous peak.

In contrast to the whole genome datasets, *k*-mers and reads extracted from hemizygous regions only show a single peak of coverage that for most datasets corresponds to the ‘heterozygous’ peak of the whole genome plots (Fig. 3a,b, bottom rows). In the *k*-mer plots of *C. sinensis* and *P. canaliculata*, the hemizygous peaks have slightly reduced coverage relative to the whole genome heterozygous peaks, possibly due to high error rates in these datasets, and the same is true for the *P. maximus* read coverage datasets.

Unlike the other datasets, the *A. immaculata* hemizygous *k*-mer plot has a second small peak corresponding to the homozygous peak of the whole genome *k*-mer histogram. This may be explained by the fact that a member of the *Achatina* (actually *Lissachatina*, see [39]) genus underwent a whole genome duplication event prior to the divergence of *A. fulica* and *A. immaculata [43]*. Duplicated hemizygous regions would be expected to encode a proportion of identical *k*-mers on each of the two copies and this would result in a peak of coverage corresponding to the whole genome homozygous peak.

### Gene content in hemizygous regions, and gene family overrepresentation

We have examined the gene content of hemizygous regions within our target species, directly by extracting the annotation of these genes, and more indirectly, by looking at the over-representation (enrichment) of both Pfam domains and GO terms within these gene complements. On several occasions, hemizygous genes appear to be associated with clusters of tandemly duplicated genes: for example *ADAM17 in P. canaliculata, GTPase IMAP family member 9* in *A. immaculata, Deoxycytidylate deaminase* in *O. sinensis*, and numerous other examples.

The number of genes associated with hemizygous regions in *O. sinensis* (0.14% of the total in its genome, Table 2) is markedly fewer than that found in the bivalve and gastropod species. As such, the subsequent absence of GO term enrichments and the limited number of Pfam domain enrichments associated with these loci were in line with expectations. We do however see many zinc finger domain genes in our blast results for Octopus (7/35 genes). Similarly, the gastropods *A. fulica* and *A. immaculata* display limited gene and domain enrichments which may be the result of decisions made during gene annotation in these species (see below for more details). Visual inspection of the *A. immaculata* hemizygous gene set revealed many repeat-containing genes with poor annotations, resulting in poor enrichment analyses, both for Pfam domains and GO terms. Interestingly, most of the annotated genes in the hemizygous regions of the *Achatina* species seem to be involved in disparate, unrelated processes, the functions of which can only be speculated upon.

While the genes found in hemizygous regions belonged to a variety of gene families, transposable element linked domains and GO terms were over-represented in our analyses, with a tendency for these genes to be involved in break repair/genomic stability, or immunity. We have investigated these genes in particular below, with full details of blast and enrichment analyses in Supplementary Files 3 and 4.

#### Transposable Elements

Within hemizygous regions, transposable element (TE) associated genes are common in most of our target species. In *P. canaliculata*, for example, we see classical retrotransposon-like (gag-pol polyprotein) elements in our gene lists (Supplementary File 3). In several species the number of hits to TEs are more striking in Pfam domain enrichment analyses, as HMM profile-based searches are much more sensitive than BLAST where clear TEs are often slightly below the *E*-value threshold for annotation. This is also clear in our over-representation analysis of GO terms, particularly for *P. canaliculata, S. broughtonii* and *P. maximus*. GO terms such as GO:0003964 “RNA-directed DNA polymerase activity”, GO:0006313 “transposition, DNA-mediated” and GO:0015074 “DNA integration”, and associated functions, are clearly enriched in these gene sets. Pfam enrichment analyses show the same clear signal of transcriptional element over-representation. Domains such as PF13975.6 “gag-polyprotein putative aspartyl protease; retrotransposon-associated”, PF00078.27 “reverse transcriptase” and PF03221.16 “Tc5 transposase DNA-binding domain” are conspicuous in these lists of enriched domains.

Some species do not possess TE-associated genes in hemizygous regions, eg. the two snails *A. fulica* and *A. immaculata*, and the bivalve *C. sinensis*. This may be a technical artefact arising from the removal of over-represented (repetitive) genes during gene annotation, or it could represent a true biological observation. Possible explanations for the association of hemizygous DNA and transposons include the use of hemizygous regions to aid transposon replication, the difficulty of hemizygous regions to purge TEs or that hemizygous regions are formed through the action of TE replication. This pattern of TE-associated gene enrichment is also seen in plant pan-genomes (e.g. [44]) suggesting a broader pattern to the link between TEs and hemizygosity.

The over-representation of Zinc-finger domains in hemizygous regions in several of our species is also interesting, given that clusters of these genes are known to be found at hotspots for copy number variation, as a means of defence against endogenous retroviruses [45]. It is possible that these genes are playing a similar role here - zinc-finger genes have a conserved role as transcriptional repressors, although their exact functionality and DNA binding affinities have yet to be fully investigated.

#### DNA break repair and remodelling

We note that many of the genes found in hemizygous regions can be linked to DNA stability, repair and remodelling. GO terms such as GO:0006281 “DNA repair”, GO:0045739 “positive regulation of DNA repair” GO:0000733 and DNA strand renaturation (*P. maximus)* and GO:0006310 “DNA recombination” (*S. broughtonii* and *P. canaliculata*) were found to be enriched, indicating that hemizygous regions may code for their own stability. Even within the limited hemizygous-associated gene set of *O. sinensis* is a *SETMAR* orthologue which in primates is known to play a role in DNA double-strand break repair, stalled replication fork restart and DNA integration [46].

In the section above, we note the presence of Zinc-finger domain genes in hemizygous regions in several species, which are known to aid genome stability at otherwise fast-evolving sites. Several of the genes found in *P. maximus* also have significant homology to PIF1, RECQL and Werner syndrome ATP-dependent helicases, which are all involved in genome stability e.g. [47]. Helicase-like domains were associated with dispensable genes in mussel *M. galloprovincialis [11]*, although they share little to no primary sequence homology with matches in UniProt. Their possible implication in the structural aspects of hemizygous regions is an excellent target for future research.

In *P. maximus* we note the presence of several G-quadruplex GO annotations (GO:0044806 G-quadruplex DNA unwinding GO:0051880 and G-quadruplex DNA binding) in addition to G/C peaks located at homozygous/hemizygous DNA transition boundaries of all species investigated here (Fig. 2b). G-quadruplexes are four-stranded DNA or RNA secondary structures formed from guanine tetramers. While a complete understanding of their function is still being elucidated, their presence in telomeres, promoter sequences and retroelements suggests a link to genome stability, gene regulation, transposon and retroviral biology [48–51].

#### Immunity

Overall, several immunity-related domains are shared in the species investigated here, but each species has its own characteristic profile. This is also the case for GO term over-representation, although we do note the importance of ontologies such as GO:0002230 “positive regulation of defense response to virus by host”, GO:0045087 “innate immune response” and GO:0051607 “defense response to virus” in our enrichment analyses.

We commonly observe genes encoding *immunoglobulin-domain containing proteins* and *C-type lectins* in our hemizygous region datasets in a number of species. These have also been observed as over-represented in *M. galloprovincalis [11]*. The function of these is yet to be fully understood, but due to their high plasticity in protein-protein and protein-carbohydrate interactions, they have been noted elsewhere as potentially important tools for immune recognition [11].

*AIG1 immunity related GTPase* genes are also observed. These are also subject to presence/absence variation (PAV) in mussels. *AIG1 immunity related GTPase* gene function is still obscure, but it plays an important role in host-parasite interactions in gastropods [52].

We also note the presence of defense peptides in hemizygous regions of these genomes. The presence of *Stomoxyn*, *toxin 32* and other Antimicrobial peptide (AMP) annotations might indicate components of the innate immune system are present in hemizygous regions. These are characterized by high intraspecific sequence diversity [53], and hemizygosity would result in greater variation between individual phenotypes for these genes.

### Individual gene families, hemizygosity and PAV

As an assay for the impact of hemizygosity on gene duplication rates and gene evolution more generally, we have studied in detail two gene families where multiple genes were found in hemizygous regions in multiple species, the HSP70 superfamily and the C1qDC containing genes. These are involved in resilience to stress and are important pattern recognition receptors in innate immunity of invertebrates, respectively [54–57], and it is possible that PAV in hemizygous regions is linked to differential adaptive capacity across the ranges of these genes.

Both HSP70 and C1qDC gene family expansions have been noted previously in bivalves [58] and we observe that their occurrences within hemizygous regions are much more prevalent in bivalves than in other molluscs, despite gastropods also possessing multiple duplicates of these genes. In Fig. 4a, it can be seen that multiple lineage specific duplications have occurred in many of the genes and gene families within the HSP70 superfamily and these duplications are not limited to bivalves. *A. fulica* and *A. immaculata* in particular share many duplicate, paralogous copies of *HSP70* (*HSPA1)*, however none of these are found in hemizygous regions. In contrast, the disparate *HSP70-12* (*HSPA12*) genes do frequently occur in hemizygous regions (Fig. 4b). Full sequences, alignments, and alternative representations of the phylogeny for these figures (showing all bootstrap support values) can be found in Supplementary File 5.

**Figure 4:**
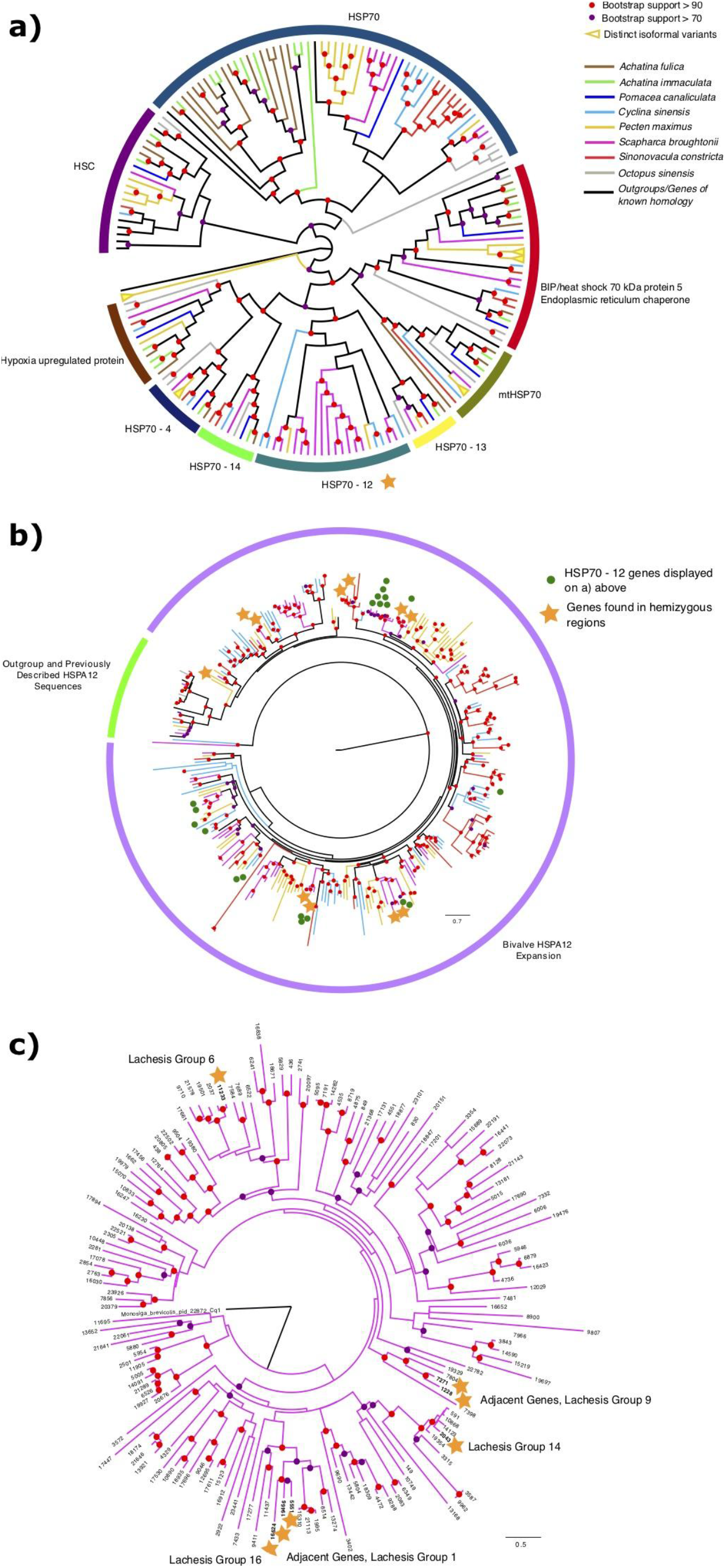
Phylogenies and cladograms of HSP70 superfamily and C1qDC genes, showing the potential of hemizygous regions as a reservoir and driver of gene diversity. a) Diagrammatic cladogram of HSP70 superfamily genes from the 8 species examined here, with branches coloured according to species identity as seen in the key, and rooted with *Arabidopsis thaliana* HSP70 sequence. Arcs surrounding the cladogram indicate gene families. Phylogeny upon which this cladogram is based, inferred using the LG+R6 model along with raw sequences, alignment and tree file, available in Supplementary File 5. Note: this cladogram is not exhaustive and excludes some HSP70-related gene sequences due to alignment and trimming. b) *HSP70-12* phylogeny genes from the 8 species examined here, along with outgroups and genes of known identity. Phylogeny inferred using the LG+F+R8 model. Note genes from hemizygous regions, indicated with a star. Genes also included in Fig. 4a above indicated with a greet dot. Phylogeny rooted with *Arabidopsis thaliana* HSP70 sequence. c) C1qDC superfamily gene interrelationships in *Scapharca broughtonii*, displayed in a phylogeny reconstructed using the WAG+F+R6 model. Note genes from hemizygous regions, indicated with a star. The linkage groups for these genes as assigned by Lachesis, are also noted alongside them.

#### HSP70-12

There is very little diversity of *HSP70-12* sequence in gastropods (1 copy in each of the 3 species examined here) or in *O. sinensis* (3 copies). However, this family of genes has exploded in bivalves. This is especially prominent in *S. broughtonii*, which possesses 8 copies within hemizygous regions, and 76 copies overall*. S. broughtonii* also possesses an *HSP90* gene within a hemizygous region (*EVM0009939*). *S. constricta* (2) and *P. maximus* (1) also possess hemizygous copies of this gene, and more than 60 paralogues in total (Fig. 4b). No *C. sinensis* copies of this gene are within hemizygous regions, although it possesses 66 copies spread across its genome.

*HSP70 - 12* genes have been studied in detail in scallops [57], where they are known to be protective against toxic dinoflagellates. In that study, drastic expansion of the *HSP70 - 12* family was observed, with a total of 47 paralogous copies of *HSP70-12* noted, although the authors do not draw any link to hemizygosity. The large numbers of *HSP70 - 12* genes observed here are therefore not unusual for bivalves. Given their role in protecting against specific pathogens, which may vary across the ranges of these species, PAV for *HSP70-12* may provide an adaptive phenotype, although we have not formally tested this here.

In our phylogeny (Fig. 4b), we note that there is little or no phylogenetic signal for an evolutionarily conserved relationship between *HSP70-12* genes from the hemizygous regions of different species. These are often in genes separated by a number of paralogy events, in strongly supported clades. Rather it seems that association events between HSP70-12 genes and hemizgyous regions, possibly through the action of transposons, have occurred in a lineage-specific manner as opposed to being derived from a common ancestor. (supplementary material, figure S1).

We observe a clear phylogenetic signal for a close relationship between pairs of hemizygous genes within species. Of the 8 copies of *HSP70-12* seen in *S. broughtonii*, all are paired with a gene of similar sequence, indicated by a star on Fig. 4b. All of these paired genes, when located in the genome, were found in close proximity. These pairs are: genes *11804* and *3214*, found on pseudochromosome “Lachesis Group 6” 10kb bp apart, genes *1411* and *9653*, from Lachesis Group 11 (8 kb apart), *3314* and *13896*, from Lachesis Group 8 (5kb apart), and *17648* and *3510*, from Contig00525, separated by only 6kb. These paired genes are all likely tandem duplicates, potentially mediated by the action of TEs.

In the *S. constricta* genome, the two *HSP70-12* genes seen at hemizygous loci are *evm.model.Chr12.1234* and *evm.model.Chr12.1237*, which are nearly 50,000 base pairs apart, at sites 37468857-37468878 and 37518784-37518839 on chromosome 12, in two separate hemizygous sites.

#### C1qDC

We also investigated the diversity of C1qDC containing genes, as these have been noted previously as being protective against environmental and pathogenic impacts [54,55], and subject to PAV in mussels [11]. These genes are widespread in molluscs, and in bivalves in particular, with more than 100 copies commonly observed in complete genomes. We found copies of C1qDC genes in hemizygous regions (see Supplementary File 3) of *C. sinensis* (1 copy, *evm.model.Hic_asm_12.1571*) and *S. broughtonii* (7 copies). We therefore chose *S. broughtonii* for specific investigation, as can be seen in Fig. 4c.

Of the 7 copies of C1qDC genes found in hemizygous regions, 2 pairs of adjacently located genes were observed (in “Lachesis Groups” 1 and 9), and 3 single gene loci were found (in “Lachesis Groups” 6, 14, and 16). The pair of genes found in Lachesis Group 1, *EVM0005551* and *EVM0019466*, are 44kb apart from one another, at sites 21191896-21194964 and 21238944-21240116 respectively, in a single deletion (pbsv.DEL.17342). The pair in Group 9, *EVM0007271* and *EVM0001228*, are only 24kb apart, in a single deletion (pbsv.DEL.114852) at positions 42253538-42255235 and 42278930-42291255. Given their relatively broad spacing and the relatively long branches separating these pairs phylogenetically, these could be ancient, rather than recent, tandem duplicates, or could have come about by other processes.

The sister gene to the two genes found in Lachesis Group 1, *EVM0016624*, is itself found in a hemizygous region, on Lachesis Group 16. Paralogous copies of these genes are therefore also found in *trans* across the genome more broadly. It is possible their movement is mediated by the TEs enriched in hemizygous regions (see previous section), although this has not been formally tested here.

## Discussion

### The prevalence and significance of hemizygosity in molluscs and across the tree of life

The detection of hemizygosity in an individual genome is indirect evidence of PAV of chromosomal regions and possibly transcriptional products within a population. This is due to inheritance of the hemizygous region-containing chromosome pairs from the two parents which themselves may have been hemizygous, homozygous or nullizygous for the particular locus. The detection of hemizygous chromosomal regions within an individual can therefore provide an initial indication of the level of chromosomal and genic variability that might exist within the population or species to which it belongs.

The complete chromosomal, transcriptional and regulatory repertoire that exists within a population or species is termed its pan-genome [1]. While such a complete snapshot of a species genetic complement can only be attained through the sequencing of multiple individuals, the detection of substantial hemizygosity within a single individual can provide strong evidence of the existence of a species-level pan-genome. The investigation of the prevalence and impact of hemizygosity on evolution is still in its infancy. However, given the evidence presented here, we can begin to comment on some aspects of this, and suggest fertile ground for future investigations.

Pan-genomes have been described in plants, fungi and bacteria, and it is only recently that they have been noted in metazoans [10,11,59,60]. To date evidence from animals is sparse, and with patchy phylogenetic distribution [4]. Here we show that, at least among conchiferan molluscs, hemizygosity is a common phenomenon. Whether this is true of other metazoan phyla remains to be investigated. However, the read-mapping approach used here would be straightforward to apply to other taxa with little additional cost, as long as a chromosome-scale genome assembly is available, and would give an initial indication of the ubiquity of this phenomenon.

While the link between hemizygosity and pan-genomes is clear, estimating a species pan-genome size based on the hemizygosity estimates of a single individual is not possible. As such, an accurate pan-genome description can only be provided by re-sequencing of multiple individuals. In general, only traditional model organisms and humans have been the target of deep re-sequencing efforts. The most extensive re-sequencing efforts of any metazoan species have come from humans and recent results suggest that here species-wide genomic variation is associated with relatively small and rare structural variants, which account on average for 5Mb of DNA sequence per individual, i.e. < 0.2% of the genome size [59]. By considering population size and the level of presence/absence variation observed between the three genome assemblies in the study, it was estimated that the human pan-genome would likely include an extra 19-40 Mb of DNA relative to the reference genome, ie. up to a 1.25% increase. However a more recent study that resequenced 910 humans of African descent extended this to 296.5 Mb of additional DNA which equates to approximately a 10% increase on the reference human assembly [9]

In contrast with animals, in plants and fungi, hemizygosity is likely to be quite widespread across most genomes [4]. In fungi, some species have open pan-genomes, with up to 60% of coding gene sequences found to vary between individuals (e.g. *Parastagonospora* spp. [61]) although figures for genomic sequence variability are not yet available. In plants, figures of up to 42% of the complete genome being absent (the “accessory/variable/dispensable genome”) in some individuals have been reported [62]. However, these values are perhaps at the extreme end of the structural variation continuum in these clades. Core genomic retention of > 80% of the complete genome is more common [4].

While it is not possible to estimate pan-genome size based on hemizygosity levels within a single individual, our figures suggest that molluscs are likely to display less genomic PAV than is seen in plants but far more than has been currently observed in vertebrates, with the maximum recorded hemizygosity seen here 6.69%.

For each hemizygous locus observed, there are four genotypes possible for each parent and 16 possible crosses (supplementary material, figure S2). Of these, 14 crosses could potentially give rise to a heterozygous (hemizygous) offspring. As both heterozygous (hemizygous) and homozygous positive individuals possess at least one copy of the locus, in the binary determination of presence or absence, both states would be counted as examples of locus presence while only those homozygous negative (nullizygous) for the locus would be counted as absence.

This means that for each hemizygous locus identified in an individual, six of the potential 14 genotype crosses involve one parent with at least one copy of the locus and the other parent with no copies of the locus. As a result, if the parents of the hemizygous individual were sampled, there would be a 3/7 chance that each locus would be subject to presence/absence variation between the two parents. If this ratio is applied to total nucleotides located within hemizygous regions, for the species with the highest observed rate of hemizygosity in this study, *Scapharca broughtonii* at 6.69%, the total genome percentage subject to presence/absence variation between the two parents would be 2.87% - at least an order of magnitude higher than the <0.2% difference between individuals observed in humans [59]. This number is also likely to be a significant underestimate as it only considers loci that are hemizygous in the offspring of the two individuals and does not take into account loci for which the sampled individual is nullizygous.

Until more phyla are sampled, we cannot comment on whether the situation in molluscs or vertebrates is more representative of the norm in the Metazoa. However, it is likely that hemizygosity and the existence of pan-genomes is much more common than we presently appreciate.

### Sex chromosome evolution and hemizygosity

Sex chromosomes represent a possible reservoir of hemizygosity which must be taken into consideration when presenting widespread evidence of this phenomenon across the Mollusca. In mammals, hemizygosity is largely restricted to sex chromosomes [63]. In molluscs, no sex chromosome has ever been described and consistent with their absence, hemizygous DNA appears to be fairly evenly distributed between and within all chromosomes (Figs. 1d, 2). Although a small amount of variability in the density of hemizygous loci can be observed across some chromosomes, we have found no evidence to suggest a link between hemizygosity and sex determination in the targeted species, which is also consistent with the observations previously collected in *M. galloprovincialis* [11].

In mammals, the chromosome carrying the sex determining gene (*SrY*) has become progressively degenerated over time through a process of recombination suppression and genetic drift [63]. In contrast, the evidence shown here suggests mollusc autosomes have become hemizygous through transposon and/or retroviral activity.

### Hemizygosity and mollusc biology

Molluscs, and particularly bivalve molluscs, are commonly broadcast spawning species with high population numbers, and high levels of genomic heterozygosity. This heterozygosity can at least in part be explained by hemizygosity. In the *k*-*mer* plots of total mapped reads (Fig. 3a, top row), the first peak (heterozygous peak) has a coverage approximately half that of the second peak (homozygous peak) however *k*-*mer*s that come from unique (non-repetitive) hemizygous regions form coverage peaks corresponding to the whole genome heterozygous peaks and so will also contribute to this ‘heterozygous’ peak (Fig. 3a, bottom row). *k*-mers that come from repetitive hemizygous regions (greater than two copies within hemizygous regions or one copy in a hemizygous region and at least one copy in a homozygous region) will not contribute to either the heterozygous or homozygous peaks as their coverage will be at least 1.5x that of homozygous k-*mer*s. Finally, *kmer*s for which there are two copies both located within hemizygous regions would be expected to contribute to the ‘homozygous’ peak.

While the number of two copy hemizygous *k*-*mer*s is expected to be low and thus comprise only a tiny portion of the homozygous peak, organisms with significant non-repetitive hemizygous DNA content should have a significant portion of their putative heterozygous peak being composed of hemizygous *k*-*mer*s. Many molluscs have very high reported heterozygosity levels as determined by *k*-*mer* analysis but our findings suggest that this rate may be at least partially explained by large non-repetitive hemizygous DNA content which is impossible to discern from heterozygous DNA using a *k*-*mer*-based approach. This has also been noted elsewhere [11], but the additional evidence provided by the extra species sampled here makes this obvious.

At first sight, the *M. galloprovincialis* genome [11] is characterized by somewhat “extreme” levels of hemizygosity and gene PAV compared to the species considered in this study. The fraction of hemizygous genomic sequence in the mussel is nearly sixfold that of *S. broughtonii*, the species with the highest hemizygosity among those included in this study (i.e. 36.78% vs 6.69%). Moreover, ~35% mussel protein-coding genes were encoded by hemizygous genomic regions, as opposed to < 2% found in the eight molluscan species studied here (Table 2). These differences may be partially attributable to the fact that the estimates provided here derive from only single individuals.

Further sampling will be needed to find the regions that are homozygous absent (nullizygous) from these specimens, but present more generally in the wider population. The size of the hemizygous regions not incorporated into these particular assemblies could be significant. Furthermore, we have not annotated ‘insertions’, regions present in these specimens but absent from the published genome assemblies. These could again represent a sizable fraction of the genome. In summary, the *M. galloprovincialis* pan-genome appears to have a significantly higher “openness” (i.e. a higher rate of dispensable/core genes) than most other molluscs, but this could be artefactual and bears further investigation.

The very low level of heterozygosity observed in *A. fulica* when compared to *A. immaculata* and the other species examined here may be the outcome of founder effects following its unintentional introduction to China sometime prior to 1931 [64,65]. Both *Achatina* species are invasive in China however due to the sparsity of information available on the size and diversity of the introduced populations, assumptions regarding the impacts of these events on the genetic diversity of the two subsequent populations would be speculative. What is clear is that genetic diversity of the *A. fulica* specimen is far lower than that of the conspecific *A. immaculata* specimen, however the reason for this difference is unresolved.

As noted in the results, in *A. immaculata*, a small peak of deletion-mapping *k*-*mer*s corresponds to the homozygous peak in the whole genome *k*-*mer* histogram. If these *k*-*mer*s are the result of duplicated hemizygous regions that arose and have been maintained since the whole genome duplication event that occured in an *Achatina* ancestor approximately 70 million years ago (MYA) [43], this would have significant consequences for our understanding of hemizygous DNA evolution, birth/death dynamics and long term persistence within genomes.

### Gene content in hemizygous regions, and gene family overrepresentation

The idea that dispensable genes may provide improved adaptation potential, suggested ever since the first definition of the pan-genome concept in scientific literature [66], is now broadly accepted thanks to multiple large-scale genome resequencing studies carried out in several prokaryotes and in a few eukaryotes. The number of pangenomic investigations carried out in metazoans remains extremely limited, and the only molluscan species which has been so far targeted by this type of investigation is the Mediterranean mussel *M. galloprovincialis*. In line with the adaptive nature of prokaryotic and eukaryotic pan-genomes, mussel dispensable genes were found to be highly enriched in functions related with innate immune response and survival, which may explain the high biotic resilience and invasiveness of this species [11].

Even though the enrichment of gene families associated with adaptation were not as strong in our datasets, some notable overlaps were identified. For example, AIG1 immune-related GTPases, expanded in several stress-adapted invertebrates [56], were enriched in *P. canaliculata*, and recurrent annotations linked with stress-related HSP70-like proteins, as well as with C1qDC proteins and C-type lectins, which are thought to act as soluble receptors for microbe-associated molecular patterns (MAMPs) in the molluscan immune system [67] were also found.

We have investigated the contribution of *HSP70-12* and *C1qDC* genes to hemizygous regions in particular here. In both of these families, multiple gene copies are found in hemizygous regions in bivalves, while in cephalopods and gastropods they appear to be restricted to homozygous regions. Where they are found, they commonly occur in duplicate which is likely the result of arising through TE-mediated tandem duplication.

Dispensable genes provide a significant contribution to phenotypic plasticity in bacteria and viruses, enabling their rapid adaptation to new ecological niches and modulating the interaction of pathogenic species with their hosts [68,69]. Similarly, the presence of open pan-genomes may explain the cosmopolitan distribution of some marine microalgae, able to thrive in largely different environmental conditions thanks to the accessory metabolic functions provided by dispensable genes [17]. Moreover, dispensable genes play a key role in the interplay between plants and associated fungi, providing improved biotic resistance to the host and enhanced pathogenicity to their parasites [70], [71]. The dispensable genes in mollusc pan-genomes may likewise provide the potential for situationally, regionally and ecologically adaptive variation.

### How do these hemizygous structural variants come about, and why does gene PAV persist?

The most likely sources of hemizygous DNA are transposon and/or retroviral insertions. Evidence for this lies in the enrichment for retroelement associated genes in several of the species investigated here, in addition to the conserved GC bias profile of the homozygous/hemizygous boundaries (Fig. 2b). Retroelements are known to encode G-rich sequences at their 5’ and/or 3’ boundaries which form four-stranded secondary structures known as G-quadruplexes [72]. G-quadruplexes have also been implicated in transposon-regulating Piwi interacting RNA (piRNA) biogenesis [73,74] and the GC enrichment observed here at the homozygous/hemizygous boundaries is consistent with transposon terminal G-quadruplexes (Fig. 2b). The outcome of retroelement-mediated gene duplication coupled with rapid gene turnover is likely to present as lineage-specific gene expansion as has been previously observed for both HSP70 and C1qDC (supplementary material, figure S1, [58]).

Hemizygous regions likely impose a cost on the genomes that carry them. The additional load of transposable elements, coupled to the over-representation of DNA repair related genes they encode, suggests that hemizygosity goes hand-in-hand with a decrease in DNA stability. Hemizygosity, if prevalent, may also interfere with homologous recombination-based DNA damage repair mechanisms through recombination suppression (supplementary material, figure S3), and could potentially impact breeding between populations with high levels of haplotypic variation through post-zygotic selection, as previously suggested in *M. galloprovincialis* [11,75,76]. Furthermore, the insertion or deletion of large blocks of DNA, regardless of their coding capacity, is likely to impact flanking gene expression due to modification of the cis-regulatory lansdscape as was recently demonstrated in the tomato [3]. How, then, are these regions not rapidly purged from populations by natural selection?

It is possible that hemizygosity, while adding to the standing pool of genetic variation and thus adaptive at a population-level, results in a larger number of errors while “crossing-over” during meiosis. This could result in a need for a higher number of double strand break repair genes (e.g. here, *tankyrase, RAG51*), and similar repair mechanisms. These could migrate into hemizygous regions over time (through TE/retroviral action) and be conserved by natural selection. Alternatively, new hemizygous DNA that is introduced but does not include stability genes on arrival, could be purged quickly leaving only those hemizygous regions that encode stability related genes left for us to observe. It is also possible that the large population size of many mollusc species makes these alleles (which could be rare) difficult to purge from populations as a whole. These hypotheses have not been tested rigorously here, but as additional data becomes available across the tree of life, these questions will be able to be addressed coherently.

### Future Perspectives and Open Questions

There are a number of “known unknowns” still to resolve regarding genes in hemizygous sites. None of the genomes investigated here have annotated long non-coding RNAs (lncRNAs) or small non-coding RNAs (snRNAs), and therefore we are unable to speculate as to whether these are also found in these regions, even though a large number of dispensable LncRNA genes have been reported in *M. galloprovincialis* [11].

In order to make reliable quantitative comparative assessments of hemizygosity between genomes, future comparative studies should utilise genomes built with consistent assembly and annotation pipelines. Independently built genomes like those assessed here likely suffer from assumptions made by the underlying software regarding how to treat hemizygous regions. Under some scenarios, longer (but lower coverage) alleles might be preferred while other pipelines may prioritise more consistent higher coverage, with low coverage alleles excluded from the final assembly. Crucially for gene enrichment analyses, custom repeat libraries might flag repetitive transposon-associated genes, marking them for exclusion from final gene sets. This would result in their exclusion from subsequent enrichment analyses and may explain why enrichment for TE-associated genes was not universally detected here. Standardised assembly and annotation pipelines would aid in dealing with these issues.

The widespread presence of hemizygosity in mollusc genomes also suggests that some modern assembly algorithms may need adjustment to take into account the prevalence of hemizygous regions. Upon encountering hemizygous regions with coverage significantly below that of the remainder of the assembly, it is plausible that some algorithms may break contigs at the point of low coverage in the assumption that the low coverage region corresponds to contamination or other artefact. Haplotype-aware assembly algorithms will likely cope with this in many cases. However, along with haplotype-blind assembly methods, even haplotype-aware assemblers may ignore the “deletion” genotype, particularly when generating a haploid approximation of the full diploid genome sequence.

Resequencing of multiple individuals will be important to obtain a truer picture of the complete pangenomic complement, and to determine how common particular dispensable genes and genomic regions are within the species. When performing re-sequencing experiments, it would be interesting to contrast wild caught individuals and those that have gone through bottlenecks (domesticated/island effect) to test the impact of these on genomic evolution.

## Conclusions

In this work we have put forward the first systematic investigation of the prevalence of hemizygosity across a metazoan clade. We have found that a number of recently sequenced conchiferan molluscs show widespread hemizygosity at multiple loci across their genomes. Bivalves in particular have a striking pattern of hemizygosity, which may reflect the broadcast spawning lifecycle of the species sequenced. Genes found in these regions in the mollusc species examined are enriched for functions related to transposition, DNA repair and stress response, suggesting that these loci could be both linked to repetitive elements, and could provide adaptive potential under specific environmental circumstances.

This approach, which is both cost-effective and broadly applicable, will be useful for assaying for the presence and utility of hemizygosity more generally across the tree of life. This phenomenon remains under-investigated, but may have profound implications for our understanding of genomic evolution at both the population and species level.

## Acknowledgements

Our thanks to Dr Angus Davison and Dr Maurine Neiman for organising the Pearls of Wisdom symposium and curating this special issue. We thank the members of our labs for their support in our work. In particular, we thank Samuele Greco of the Gerdol lab for his assistance. We acknowledge and thank Noah Schlottman, Casey Dunn, Nobu Tamura, T. Michael Keesey, Scott Hartmann, Katie S Collins, and Brockhaus and Efron for their Phylopic images (http://creativecommons.org/licenses/by-sa/3.0/).

## Supplementary material

### Supplementary Files

Supplementary File 1: All commands used to run analyses in the command line (.txt)

Supplementary File 2: *Octopus sinensis* heterozygosity calculation, Genomescope (.tar.gz)

Supplementary File 3: All blast hits to genes found in hemizygous regions (zipped .xlsx)

Supplementary File 4: Enrichment analyses, full data (zipped .xlsx)

Supplementary File 5: Sequences, alignments, initial tree files and alternative displays of final trees (inc. all bootstrap support values) for trees shown in Fig. 4. (.tar.gz)

### Supplementary Figure Legends

**Figure S1:** Birth and death dynamics of a hypothetical transposon-mediated gene family expansion. In the last common ancestor of two species, a single copy of a gene (1) becomes associated with a retroelement which duplicates it in tandem through two separate transposition events (2). A subsequent speciation event gives rise to two daughter species which independently undergo further retroelement-mediated duplication events (3). Over time, further independent duplication and gene loss events in each lineage give rise to distinct repertoires of the gene family (4) which, when subjected to phylogenetic analysis, display patterns of inheritance consistent with independent lineage-specific expansions.

**Figure S2:** Parental crosses that may give rise to hemizygous offspring. Of the 16 genotype combinations possible for a single locus that is subject to population-level presence absence variation, 14 have the potential to give rise to a hemizygous offspring. Those combinations that can not result in a hemizygous offspring are coloured in red. Of those 14, both parents would possess at least one copy of the hemizygous locus under eight situations (coloured in yellow) while for six matings, one parent would be nullizygous and the other either homozygous or heterozygous for the locus (green).

**Figure S3:** Hemizygous DNA leads to localised recombination suppression. Following the formation of a double strand break (DSB, pair of vertical black lines) at a homozygous locus, DNA repair through homologous recombination proceeds through second end capture by the homologous region of the chromosomal pair followed by double-Holliday junction formation. When a DSB occurs close to a region of hemizygosity, the absence of homology between the corresponding regions on the chromosomal pairs likely inhibits the formation of a double-Holliday junction as the region homologous to the ‘insertion’ (rectangular boxes) is not present on the chromosome with the ‘deletion’ (grey region).

## Ethics

No ethical permissions are required for the work carried out in this manuscript. This work does not include human tissue, vertebrate animals, fieldwork or museum specimens in its analyses.

## Data, code and materials

The datasets and code supporting this article have been uploaded as part of the supplementary material. All genomic sequences are available from the original sources with accession numbers as cited in the manuscript.

## Competing interests

The authors declare they have no competing interests.

## Authors’ contributions

ADC conceived of the study and designed experiments. ADC, NJK and MG undertook data analysis and drafted the manuscript. All authors gave final approval for publication and agree to be held accountable for the work performed therein.

## Notes

### Competing Interest Statement

The authors have declared no competing interest.

