## Supplementary File 1 for "Single individual structural variant detection uncovers widespread hemizygosity in molluscs"

**Hemizygosity - Command Line Usage**

### 1. **Structural variant detection**

#Find tandem repeats

trf genome.fa 2 7 7 80 10 50 500 -d -h

#Convert trf output to bed

TRFdat_to_bed.py --dat genome.fa.2.7.7.80.10.50.500_mod.dat --bed genome.fa.2.7.7.80.10.50.500.bed

#Convert bed to zero based six field bed with arbitrary sixth field

awk '{print $1"\t"$2-1"\t"$3"\t"$4}' genome.fa.2.7.7.80.10.50.500.bed | sed 's/Sequence://' | awk '{print $0"\t"$3-$2"\t1"}' >genome.fa.2.7.7.80.10.50.500_0based_6field.bed

#Run pbmm2

pbmm2 align -j 16 genome.fa pacbio_reads.fofn genome.aligned.bam --sort --median-filter --sample sample1

#Run pbsv

pbsv discover --tandem-repeats genome.fa.2.7.7.80.10.50.500_0based.bed genome.aligned.bam genome.svsig.gz

pbsv call -j 16 genome.fa genome.svsig.gz genome.var.vcf

#Extract DELs that are not homozygous for the alternative allele (possible genome assembly errors) as a zero based six field bed file

grep DEL genome.var.vcf | grep PASS | grep -v '1/1' | awk '{print $1"\t"$2-1"\t"$2+length($4)-1"\t"$3}' | awk '{print $0"\t"$3-$2"\t1"}' >genome.var.DEL.6field.bed

#Generate chromosome file for chromoMap. The head value equals the number of chromosomal scaffolds in the genome.fa file

bioawk -c fastx '{ print $name, length($seq) }' < genome.fa | head -19 | awk '{print $1"\t1\t"$2}' >genome_chrom.txt

#Make feature file for chromoMap of the >10k deletions that are not annotated as tandem repeats

grep DEL genome.var.vcf | grep PASS | grep -v '1/1' | grep -v TANDEM | awk '{print $1"\t"$2-1"\t"$2+length($4)-1"\t"$3}' | awk '{print $0"\t"$3-$2"\t1"}' | awk '$5>9999' | awk '{print $4"\t"$1"\t"$2"\t"$3}' >genome_dels.txt

#Run chromoMap in R

chromoMap("genome_chrom.txt","genome_dels.txt",canvas_width = 1200,canvas_height = 2000,left_margin = 100)

### 2. **Kmer analysis of hemizygous regions**

#Preprocess illumina reads

bbduk.sh in1=reads_1.fastq in2=reads_2.fastq out1=reads_clean_trimmed_1.fq out2=reads_clean_trimmed_2.fq ref=adapters.fa ktrim=r k=25 mink=11 hdist=1 qtrim=r trimq=30 tpe tbo threads=8

#map cleaned data to genome

bwa mem -t 16 genome.fasta reads_clean_trimmed_1.fastq reads_clean_trimmed_2.fastq | samtools sort -@16 -o reads_bwa_aligned.bam -

#merge bam files if necessary

samtools merge mergedBamFile.bam *.bam

#Get mapped reads in fasta format

samtools view -@ 8 -F 4 -h mergedBamFile.bam >all_mapping.sam

reformat.sh in=all_mapping.sam out=all_mapping.fa

#run jellyfish on all mapped reads

jellyfish count -t 8 -C -m 21 -s 16G all_mapping.fa -o all_reads.jf

jellyfish histo -o all_reads.histo all_reads.jf

#Extract reads that map to deletions. The bedmap step makes sure the reads are fully contained within the deleted regions and not just partially overlapping.

samtools view -@ 8 -F 4 -h -b -L genome.var.DEL.6field.bed mergedBamFile.bam >del_mapping.bam

bedmap --echo --fraction-map 1 <(bam2bed <del_mapping.bam) genome.var.DEL.6field.bed >del_reads.bed

cut -f1-6 del_reads.bed >del_reads.6field.bed

fastaFromBed -fi genome.fa -bed del_reads.6field.bed -fo del_reads.fa

#Run jellyfish on deletion mapping reads

jellyfish count -t 8 -C -m 21 -s 16G del_reads.fa -o del_reads.jf

jellyfish histo -o del_reads.histo del_reads.jf

### 3. **Read coverage of hemizygous regions**

#Get names of reads that map to deletions

cut -f4 del_reads.bed | sort -u >del_reads.names

#Get sam file of del mapped reads

samtools view mergedBamFile.bam | fgrep -w -f del_reads.names >del_reads.sam

#Now manually add headers back to sam

samtools view -H mergedBamFile.bam >del_reads_header.sam

cat del_reads.sam >>del_reads_header.sam

#Convert sam to bam and index

samtools view -@8 -S -b del_reads_header.sam >del_reads_header.bam

samtools index -@8 del_reads_header.bam

#Run mosdepth

mosdepth -t 8 -m -b del_reads.6field.bed genome del_reads_filtered.bam

#Count how many deletions have each level of coverage

zcat genome.regions.bed.gz | awk 'BEGIN{OFS=FS="\t"}{$6=sprintf("%.0f",$5) }1' | cut -f6 | sort -n | uniq -c | sed -e 's/^ *//' -e 's/\ /\t/' | awk '{print $2"\t"$1}' | head -200 >deletion_coverage.txt

### 4. **Read coverage of genome with sliding window**

#Run genomeCoverageBed to calculate coverage at every position in the genome

genomeCoverageBed -d -ibam mergedBamFile.bam >genome.cov

#split coverage file by scaffold/chromosome

awk '{print>$1".cov"}' genome.cov

#Calculate the median coverage of every 1000 bp window with median_sliding_window.gawk script - script available in section 5

for i in *cov ; do cat $i | sh median_sliding_window.gawk | sed 's/\

/\t/g' >`echo $i | sed 's/cov/median/'` ; done

#count how many windows have each coverage value

cat *median | awk 'BEGIN{OFS=FS="\t"}{$4=sprintf("%.0f",$3) }1' | cut -f4 | sort -n | uniq -c | sed -e 's/^ *//' -e 's

/\ /\t/' | awk '{print $2"\t"$1}' >coverage_count.txt

### 5. **Median_sliding_window.gawk script**

#!/usr/bin/sh

gawk -v wsize=1000 '

BEGIN {

if (wsize % 2 == 0) { m1=wsize/2; m2=m1+1; } else { m1 = m2 = (wsize+1)/2; }

}

function roundedmedian() {

asort(window, a);

return (m1==m2) ? a[m1] : int(0.5 + ((a[m1] + a[m2]) / 2));

}

function push(value) {

window[NR % wsize] = value;

}

NR < wsize { window[NR]=$3; next; }

{ push($3);

$3 = roundedmedian();

print $0;

}'

### 6. **Extraction of genes from hemizygous regions**

#Extract only those genes that fall entirely within the boundaries of known hemizygous regions. -F sets the fraction of the gtf that needs to fall within the boundaries of the hemizygous region to be included - here it is 1 (so total), but you could set this to a really low number to get any amount of overlap (e.g. the default is 1E-9, and would detect 1 bp of overlap in almost any chrimosome).

bedtools intersect -a YOURGENOME_HEMIZYGOUS_REGIONS.bed -b YOURGENOME.gtf -wb -F 1 > StrictOverlap.gtf

#As the above file reports all tracks, and you might want just the gene names, extract only the gene tracks (not exons, UTR etc). NB you will have to add the tab either side of the gene manually (UNIX: control + v followed by the tab key)

cat StrictOverlap.gtf | grep "	gene	" > Overlap_genes_deletion

#Make a gtf formatted file out of the output

cut -f7-15 Overlap_genes_deletion > StrictlyFullySpanned_gene_set.gtf

#Extract just the gene names out of the output

cut -f7-15 Overlap_genes_deletion | cut -f9 | sort | uniq > Genes_Completely_Within_deletion.txt
