## Supplementary File 2 for "Single individual structural variant detection uncovers widespread hemizygosity in molluscs": GenomeScope.html


GenomeScope


Toggle navigation

 GenomeScope

- Home
- Info
- Examples 
  - Seabass
  - Oyster
  - Pear
  - Bird
  - Drosophila
  - Arabidopsis F1
- My results 
  - 'my sample' (0 seconds ago)


### my sample

#### Results

Model did not converge

#### Model

Model did not converge

### View analysis later

Return to view your results at any time:


### Progress

starting

round 0 trimming to 6 trying 4peak model... unconverged

round 1 trimming to 11 trying 4peak model... unconverged

round 2 trimming to 16 trying 4peak model... unconverged

round 3 trimming to 21 trying 4peak model... unconverged

fail
