## Supplementary File 2 for "Single individual structural variant detection uncovers widespread hemizygosity in molluscs": analysis_page_script.js

```
var analysis_path="analysis.php?code=";
var description = "";

var num_fails_getting_progress_file = 0;

//////////////////////////////////////////////////////////////
/////// For analysis page:
//////////////////////////////////////////////////////////////

function showProgress() {
    var run_id_code=getUrlVars()["code"];
    var prog=0;

    if (description == "") {
        jQuery.ajax({ 
            type:"POST",
            url: "user_data/" + run_id_code + "/description.txt",
            success: function (obj) {
                description = obj;
                document.getElementById("description_header").innerHTML = description;
            }
        });
    }

//  remember ajax is asynchronous, so only the stuff inside the success: part will be called after retrieving information. If I put something after the statement, it can't use the info from check_progress.php because it is executed before this php script is called
    //alert('before ajax');
    jQuery.ajax({ 
        type:"POST",
        url: "check_progress.php",
        dataType: 'json',
        data: {code: run_id_code},
        success: function (obj) {
            // alert("inside success");
            // alert(obj);
            prog=obj;
            last_line = prog[prog.length-1];
            if (last_line==undefined) {
                console.log("No progress file found, may be an error on the server.");
                num_fails_getting_progress_file++;
                if (num_fails_getting_progress_file > 10) {
                    document.getElementById("progress_panel").className = "panel panel-danger center";
                    document.getElementById("plot_info").innerHTML = "No progress file found, may be an error on the server";
                }
                setTimeout(function(){showProgress();},500);
            } else {
                output_array = prog; //.slice(1,prog.length);
                output_info = ""
                for (var i=0;i < output_array.length; i++) {
                    line = output_array[i];
                    output_info += "<p>" + line + "</p>";
                }

                document.getElementById("plot_info").innerHTML = output_info;

                if (last_line.indexOf('done') > -1) {
                    document.getElementById("progress_panel").className = "panel panel-success center";
                    check_plot_exists(0, true);
                }
                else if (last_line.indexOf("fail") > -1) { // SOMETHING FAILED
                    document.getElementById("progress_panel").className = "panel panel-danger center";
                    check_plot_exists(0, false);
                }
                else {
                    setTimeout(function(){showProgress();},500);
                }
            }
        }
    });

}


function getUrlVars() {
    var vars = {};
    var parts = window.location.href.replace(/[?&]+([^=&]+)=([^&]*)/gi, function(m,key,value) {
        vars[key] = value;
    });
    return vars;
}
//
//function test() {
//    var run_id_code=getUrlVars()["code"];
//    alert(run_id_code);
//}
//
var content_width = $( window ).width();


function check_plot_exists(counter,bool_succeeded) {
    
    var run_id_code=getUrlVars()["code"];
    var file_url_prefix="user_data/"+run_id_code + "/";
    var file_to_check_for=file_url_prefix + "plot.png";
    
    if (counter>100) {
        alert("Taking too long to find "+ file_to_check_for + " counter: " + counter);
    }
    else {
        jQuery.ajax({ 
            url: file_to_check_for,
            error: function() {
                console.log(counter+1);
                setTimeout(function(){check_plot_exists(counter+1,bool_succeeded);},500);
            },
            success: function () {
                document.getElementById("results").style.visibility= 'visible';
                // alert("inside success");
                // document.getElementById("landing_for_plot1").innerHTML='<img class="fluidimage" src="' + "./user_data/"+ run_id_code + "/plot.png"  + ' "/>'; // width="750" height="500" // style="max-height:100%; max-width:100%;" 
                // document.getElementById("landing_for_plot1_details").innerHTML='<iframe  width="600" height="300" src="' + "./user_data/"+ run_id_code + "/summary.html" + '" frameborder="0"></iframe>';

                document.getElementById("landing_for_plot1").innerHTML='<img class="fluidimage" onerror="imgError(this);" src="' + file_url_prefix  + "plot.png" + ' "/>'; 
                document.getElementById("landing_for_plot2").innerHTML='<img class="fluidimage" onerror="imgError(this);" src="' + file_url_prefix  + "plot.log.png" + ' "/>'; 

                if (bool_succeeded) {
                    document.getElementById("landing_for_text1").innerHTML='<iframe width="' + 600 + ' " height="200" src="' + file_url_prefix + "summary.txt" + '" frameborder="0"></iframe>';
                    document.getElementById("landing_for_text2").innerHTML='<iframe width="' + 600 + ' " height="400" src="' + file_url_prefix + "model.txt" + '" frameborder="0"></iframe>';
                } else {
                    document.getElementById("landing_for_text1").innerHTML='Model did not converge';
                    document.getElementById("landing_for_text2").innerHTML='Model did not converge';
                }
                
                imageresize();
            }
        });
    }
}


function imageresize() {
    console.log("resizing")

    var size_fraction = 1; // 1 means fit one plot on the page, 3 means fit 3 plots on the page

    var top_padding = 100;
    var side_padding = 0.05;
    var aspect_ratio = 1;
    var max_size = 1000;

    var height = Math.min(content_width/aspect_ratio*(1-side_padding), $( window ).height()-top_padding)/size_fraction;
    height = Math.min(height,max_size);
    $(".fluidimage").height(height + "px");
    $(".fluidimage").width(height*aspect_ratio + "px");


    //  Fancybox plot zooming
    // http://www.dwuser.com/education/content/click-to-zoom-for-photos-adding-lightbox-effect-to-your-images/
    var addToAll = true;
    var gallery = true;
    var titlePosition = 'inside';
    $(addToAll ? 'img' : 'img.fancybox').each(function(){
        var $this = $(this);
        var title = $this.attr('title');
        var src = $this.attr('data-big') || $this.attr('src');
        var a = $('<a href="#" class="fancybox"></a>').attr('href', src).attr('title', title);
        $this.wrap(a);
    });
    if (gallery)
        $('a.fancybox').attr('rel', 'fancyboxgallery');
    $('a.fancybox').fancybox({
        titlePosition: titlePosition
    });

    $.noConflict();

}


function imgError(image) {
    image.onerror = "";

    var parent = image.parentNode;
    parent.parentNode.removeChild(parent);
    imageresize();
    
    return true;
}


$(document).ready(function() {
    showProgress();
    $(window).bind("resize", function(){ //Adjusts image when browser resized
       imageresize();
    });
});


// How to execute code after getting info from multiple files:
    //
    //$.when(
    //    $.get(filename_input, function(csvString) {
    //        array_input = $.csv.toArrays(csvString, {onParseValue: $.csv.hooks.castToScalar}); 
    //    }),
    //    $.get(filename_output, function(csvString) {
    //        array_ouput = $.csv.toArrays(csvString, {onParseValue: $.csv.hooks.castToScalar} ); 
    //    })
    //).then(function() {
    //    console.log(array_input)
    //    console.log(typeof array_input)
    //    console.log(array_input.length)
    //    console.log(array_input[0].length)
    //    var diff=[]
    //    for (i=0; i<v_tick_max+2; i++){
    //        v_ticks.push(i);
    //    }
    //});
```
