## Supplementary File 2 for "Single individual structural variant detection uncovers widespread hemizygosity in molluscs": analytics.js

```
(function(){/*

 Copyright The Closure Library Authors.
 SPDX-License-Identifier: Apache-2.0
*/
var l=this||self,m=function(a,b){a=a.split(".");var c=l;a[0]in c||"undefined"==typeof c.execScript||c.execScript("var "+a[0]);for(var d;a.length&&(d=a.shift());)a.length||void 0===b?c=c[d]&&c[d]!==Object.prototype[d]?c[d]:c[d]={}:c[d]=b};var q=function(a,b){for(var c in b)b.hasOwnProperty(c)&&(a[c]=b[c])},r=function(a){for(var b in a)if(a.hasOwnProperty(b))return!0;return!1};var t=/^(?:(?:https?|mailto|ftp):|[^:/?#]*(?:[/?#]|$))/i;var u=window,v=document,w=function(a,b){v.addEventListener?v.addEventListener(a,b,!1):v.attachEvent&&v.attachEvent("on"+a,b)};var x={},y=function(){x.TAGGING=x.TAGGING||[];x.TAGGING[1]=!0};var z=/:[0-9]+$/,B=function(a,b){b&&(b=String(b).toLowerCase());if("protocol"===b||"port"===b)a.protocol=A(a.protocol)||A(u.location.protocol);"port"===b?a.port=String(Number(a.hostname?a.port:u.location.port)||("http"==a.protocol?80:"https"==a.protocol?443:"")):"host"===b&&(a.hostname=(a.hostname||u.location.hostname).replace(z,"").toLowerCase());var c=A(a.protocol);b&&(b=String(b).toLowerCase());switch(b){case "url_no_fragment":b="";a&&a.href&&(b=a.href.indexOf("#"),b=0>b?a.href:a.href.substr(0,
b));a=b;break;case "protocol":a=c;break;case "host":a=a.hostname.replace(z,"").toLowerCase();break;case "port":a=String(Number(a.port)||("http"==c?80:"https"==c?443:""));break;case "path":a.pathname||a.hostname||y();a="/"==a.pathname.substr(0,1)?a.pathname:"/"+a.pathname;a=a.split("/");a:if(b=[],c=a[a.length-1],Array.prototype.indexOf)b=b.indexOf(c),b="number"==typeof b?b:-1;else{for(var d=0;d<b.length;d++)if(b[d]===c){b=d;break a}b=-1}0<=b&&(a[a.length-1]="");a=a.join("/");break;case "query":a=a.search.replace("?",
"");break;case "extension":a=a.pathname.split(".");a=1<a.length?a[a.length-1]:"";a=a.split("/")[0];break;case "fragment":a=a.hash.replace("#","");break;default:a=a&&a.href}return a},A=function(a){return a?a.replace(":","").toLowerCase():""},C=function(a){var b=v.createElement("a");a&&(b.href=a);var c=b.pathname;"/"!==c[0]&&(a||y(),c="/"+c);a=b.hostname.replace(z,"");return{href:b.href,protocol:b.protocol,host:b.host,hostname:a,pathname:c,search:b.search,hash:b.hash,port:b.port}};function D(){for(var a=E,b={},c=0;c<a.length;++c)b[a[c]]=c;return b}function F(){var a="ABCDEFGHIJKLMNOPQRSTUVWXYZ";a+=a.toLowerCase()+"0123456789-_";return a+"."}var E,G;function H(a){E=E||F();G=G||D();for(var b=[],c=0;c<a.length;c+=3){var d=c+1<a.length,f=c+2<a.length,e=a.charCodeAt(c),g=d?a.charCodeAt(c+1):0,h=f?a.charCodeAt(c+2):0,k=e>>2;e=(e&3)<<4|g>>4;g=(g&15)<<2|h>>6;h&=63;f||(h=64,d||(g=64));b.push(E[k],E[e],E[g],E[h])}return b.join("")}
function I(a){function b(k){for(;d<a.length;){var n=a.charAt(d++),p=G[n];if(null!=p)return p;if(!/^[\s\xa0]*$/.test(n))throw Error("Unknown base64 encoding at char: "+n);}return k}E=E||F();G=G||D();for(var c="",d=0;;){var f=b(-1),e=b(0),g=b(64),h=b(64);if(64===h&&-1===f)return c;c+=String.fromCharCode(f<<2|e>>4);64!=g&&(c+=String.fromCharCode(e<<4&240|g>>2),64!=h&&(c+=String.fromCharCode(g<<6&192|h)))}};var J;var N=function(){var a=K,b=L,c=M(),d=function(g){a(g.target||g.srcElement||{})},f=function(g){b(g.target||g.srcElement||{})};if(!c.init){w("mousedown",d);w("keyup",d);w("submit",f);var e=HTMLFormElement.prototype.submit;HTMLFormElement.prototype.submit=function(){b(this);e.call(this)};c.init=!0}},O=function(a,b,c){for(var d=M().decorators,f={},e=0;e<d.length;++e){var g=d[e],h;if(h=!c||g.forms)a:{h=g.domains;var k=a,n=!!g.sameHost;if(h&&(n||k!==v.location.hostname))for(var p=0;p<h.length;p++)if(h[p]instanceof
RegExp){if(h[p].test(k)){h=!0;break a}}else if(0<=k.indexOf(h[p])||n&&0<=h[p].indexOf(k)){h=!0;break a}h=!1}h&&(h=g.placement,void 0==h&&(h=g.fragment?2:1),h===b&&q(f,g.callback()))}return f},M=function(){var a={};var b=u.google_tag_data;u.google_tag_data=void 0===b?a:b;a=u.google_tag_data;b=a.gl;b&&b.decorators||(b={decorators:[]},a.gl=b);return b};var P=/(.*?)\*(.*?)\*(.*)/,aa=/([^?#]+)(\?[^#]*)?(#.*)?/;function Q(a){return new RegExp("(.*?)(^|&)"+a+"=([^&]*)&?(.*)")}
var S=function(a){var b=[],c;for(c in a)if(a.hasOwnProperty(c)){var d=a[c];void 0!==d&&d===d&&null!==d&&"[object Object]"!==d.toString()&&(b.push(c),b.push(H(String(d))))}a=b.join("*");return["1",R(a),a].join("*")},R=function(a,b){a=[window.navigator.userAgent,(new Date).getTimezoneOffset(),window.navigator.userLanguage||window.navigator.language,Math.floor((new Date).getTime()/60/1E3)-(void 0===b?0:b),a].join("*");if(!(b=J)){b=Array(256);for(var c=0;256>c;c++){for(var d=c,f=0;8>f;f++)d=d&1?d>>>1^
3988292384:d>>>1;b[c]=d}}J=b;b=4294967295;for(c=0;c<a.length;c++)b=b>>>8^J[(b^a.charCodeAt(c))&255];return((b^-1)>>>0).toString(36)},ca=function(a){return function(b){var c=C(u.location.href),d=c.search.replace("?","");a:{var f=d.split("&");for(var e=0;e<f.length;e++){var g=f[e].split("=");if("_gl"===decodeURIComponent(g[0]).replace(/\+/g," ")){f=g.slice(1).join("=");break a}}f=void 0}b.query=T(f||"")||{};f=B(c,"fragment");e=f.match(Q("_gl"));b.fragment=T(e&&e[3]||"")||{};a&&ba(c,d,f)}};
function U(a,b){if(a=Q(a).exec(b)){var c=a[2],d=a[4];b=a[1];d&&(b=b+c+d)}return b}
var ba=function(a,b,c){function d(e,g){e=U("_gl",e);e.length&&(e=g+e);return e}if(u.history&&u.history.replaceState){var f=Q("_gl");if(f.test(b)||f.test(c))a=B(a,"path"),b=d(b,"?"),c=d(c,"#"),u.history.replaceState({},void 0,""+a+b+c)}},T=function(a){var b=void 0===b?3:b;try{if(a){a:{for(var c=0;3>c;++c){var d=P.exec(a);if(d){var f=d;break a}a=decodeURIComponent(a)}f=void 0}if(f&&"1"===f[1]){var e=f[2],g=f[3];a:{for(f=0;f<b;++f)if(e===R(g,f)){var h=!0;break a}h=!1}if(h){b={};var k=g?g.split("*"):
[];for(g=0;g<k.length;g+=2)b[k[g]]=I(k[g+1]);return b}}}}catch(n){}};function V(a,b,c,d){function f(k){k=U(a,k);var n=k.charAt(k.length-1);k&&"&"!==n&&(k+="&");return k+h}d=void 0===d?!1:d;var e=aa.exec(c);if(!e)return"";c=e[1];var g=e[2]||"";e=e[3]||"";var h=a+"="+b;d?e="#"+f(e.substring(1)):g="?"+f(g.substring(1));return""+c+g+e}
function W(a,b){var c="FORM"===(a.tagName||"").toUpperCase(),d=O(b,1,c),f=O(b,2,c);b=O(b,3,c);r(d)&&(d=S(d),c?X("_gl",d,a):Y("_gl",d,a,!1));!c&&r(f)&&(c=S(f),Y("_gl",c,a,!0));for(var e in b)b.hasOwnProperty(e)&&Z(e,b[e],a)}function Z(a,b,c,d){if(c.tagName){if("a"===c.tagName.toLowerCase())return Y(a,b,c,d);if("form"===c.tagName.toLowerCase())return X(a,b,c)}if("string"==typeof c)return V(a,b,c,d)}function Y(a,b,c,d){c.href&&(a=V(a,b,c.href,void 0===d?!1:d),t.test(a)&&(c.href=a))}
function X(a,b,c){if(c&&c.action){var d=(c.method||"").toLowerCase();if("get"===d){d=c.childNodes||[];for(var f=!1,e=0;e<d.length;e++){var g=d[e];if(g.name===a){g.setAttribute("value",b);f=!0;break}}f||(d=v.createElement("input"),d.setAttribute("type","hidden"),d.setAttribute("name",a),d.setAttribute("value",b),c.appendChild(d))}else"post"===d&&(a=V(a,b,c.action),t.test(a)&&(c.action=a))}}
var K=function(a){try{a:{for(var b=100;a&&0<b;){if(a.href&&a.nodeName.match(/^a(?:rea)?$/i)){var c=a;break a}a=a.parentNode;b--}c=null}if(c){var d=c.protocol;"http:"!==d&&"https:"!==d||W(c,c.hostname)}}catch(f){}},L=function(a){try{if(a.action){var b=B(C(a.action),"host");W(a,b)}}catch(c){}};m("google_tag_data.glBridge.auto",function(a,b,c,d){N();c="fragment"===c?2:1;a={callback:a,domains:b,fragment:2===c,placement:c,forms:!!d,sameHost:!1};M().decorators.push(a)});m("google_tag_data.glBridge.decorate",function(a,b,c){a=S(a);return Z("_gl",a,b,!!c)});m("google_tag_data.glBridge.generate",S);m("google_tag_data.glBridge.get",function(a,b){var c=ca(!!b);b=M();b.data||(b.data={query:{},fragment:{}},c(b.data));c={};if(b=b.data)q(c,b.query),a&&q(c,b.fragment);return c});})(window);
(function(){function La(a){var b=1,c;if(a)for(b=0,c=a.length-1;0<=c;c--){var d=a.charCodeAt(c);b=(b<<6&268435455)+d+(d<<14);d=b&266338304;b=0!=d?b^d>>21:b}return b};/*

 Copyright The Closure Library Authors.
 SPDX-License-Identifier: Apache-2.0
*/
var $c=function(a){this.C=a||[]};$c.prototype.set=function(a){this.C[a]=!0};$c.prototype.encode=function(){for(var a=[],b=0;b<this.C.length;b++)this.C[b]&&(a[Math.floor(b/6)]^=1<<b%6);for(b=0;b<a.length;b++)a[b]="ABCDEFGHIJKLMNOPQRSTUVWXYZabcdefghijklmnopqrstuvwxyz0123456789-_".charAt(a[b]||0);return a.join("")+"~"};var F=window.GoogleAnalyticsObject,H;if(H=void 0!=F)H=-1<(F.constructor+"").indexOf("String");var wa;if(wa=H){var ha=window.GoogleAnalyticsObject;wa=ha?ha.replace(/^[\s\xa0]+|[\s\xa0]+$/g,""):""}var gb=wa||"ga",jd=/^(?:utma\.)?\d+\.\d+$/,kd=/^amp-[\w.-]{22,64}$/,Ba=!1;var vd=new $c;function J(a){vd.set(a)}var Td=function(a){a=Dd(a);a=new $c(a);for(var b=vd.C.slice(),c=0;c<a.C.length;c++)b[c]=b[c]||a.C[c];return(new $c(b)).encode()},Dd=function(a){a=a.get(Gd);ka(a)||(a=[]);return a};var ea=function(a){return"function"==typeof a},ka=function(a){return"[object Array]"==Object.prototype.toString.call(Object(a))},qa=function(a){return void 0!=a&&-1<(a.constructor+"").indexOf("String")},D=function(a,b){return 0==a.indexOf(b)},sa=function(a){return a?a.replace(/^[\s\xa0]+|[\s\xa0]+$/g,""):""},ra=function(){for(var a=O.navigator.userAgent+(M.cookie?M.cookie:"")+(M.referrer?M.referrer:""),b=a.length,c=O.history.length;0<c;)a+=c--^b++;return[hd()^La(a)&2147483647,Math.round((new Date).getTime()/
1E3)].join(".")},ta=function(a){var b=M.createElement("img");b.width=1;b.height=1;b.src=a;return b},ua=function(){},K=function(a){if(encodeURIComponent instanceof Function)return encodeURIComponent(a);J(28);return a},L=function(a,b,c,d){try{a.addEventListener?a.addEventListener(b,c,!!d):a.attachEvent&&a.attachEvent("on"+b,c)}catch(e){J(27)}},f=/^[\w\-:/.?=&%!\[\]]+$/,Nd=/^[\w+/_-]+[=]{0,2}$/,be=function(a,b){return E(M.location[b?"href":"search"],a)},E=function(a,b){return(a=a.match("(?:&|#|\\?)"+
K(b).replace(/([.*+?^=!:${}()|\[\]\/\\])/g,"\\$1")+"=([^&#]*)"))&&2==a.length?a[1]:""},xa=function(){var a=""+M.location.hostname;return 0==a.indexOf("www.")?a.substring(4):a},de=function(a,b){var c=a.indexOf(b);if(5==c||6==c)if(a=a.charAt(c+b.length),"/"==a||"?"==a||""==a||":"==a)return!0;return!1},ya=function(a,b){var c=M.referrer;if(/^(https?|android-app):\/\//i.test(c)){if(a)return c;a="//"+M.location.hostname;if(!de(c,a))return b&&(b=a.replace(/\./g,"-")+".cdn.ampproject.org",de(c,b))?void 0:
c}},za=function(a,b){if(1==b.length&&null!=b[0]&&"object"===typeof b[0])return b[0];for(var c={},d=Math.min(a.length+1,b.length),e=0;e<d;e++)if("object"===typeof b[e]){for(var g in b[e])b[e].hasOwnProperty(g)&&(c[g]=b[e][g]);break}else e<a.length&&(c[a[e]]=b[e]);return c};var ee=function(){this.b=[];this.ea={};this.m={}};ee.prototype.set=function(a,b,c){this.b.push(a);c?this.m[":"+a]=b:this.ea[":"+a]=b};ee.prototype.get=function(a){return this.m.hasOwnProperty(":"+a)?this.m[":"+a]:this.ea[":"+a]};ee.prototype.map=function(a){for(var b=0;b<this.b.length;b++){var c=this.b[b],d=this.get(c);d&&a(c,d)}};var O=window,M=document,va=function(a,b){return setTimeout(a,b)};var Qa=window,Za=document,G=function(a){var b=Qa._gaUserPrefs;if(b&&b.ioo&&b.ioo()||a&&!0===Qa["ga-disable-"+a])return!0;try{var c=Qa.external;if(c&&c._gaUserPrefs&&"oo"==c._gaUserPrefs)return!0}catch(g){}a=[];b=String(Za.cookie).split(";");for(c=0;c<b.length;c++){var d=b[c].split("="),e=d[0].replace(/^\s*|\s*$/g,"");e&&"AMP_TOKEN"==e&&((d=d.slice(1).join("=").replace(/^\s*|\s*$/g,""))&&(d=decodeURIComponent(d)),a.push(d))}for(b=0;b<a.length;b++)if("$OPT_OUT"==a[b])return!0;return Za.getElementById("__gaOptOutExtension")?
!0:!1};var Ca=function(a){var b=[],c=M.cookie.split(";");a=new RegExp("^\\s*"+a+"=\\s*(.*?)\\s*$");for(var d=0;d<c.length;d++){var e=c[d].match(a);e&&b.push(e[1])}return b},zc=function(a,b,c,d,e,g,ca){e=G(e)?!1:eb.test(M.location.hostname)||"/"==c&&vc.test(d)?!1:!0;if(!e)return!1;b&&1200<b.length&&(b=b.substring(0,1200));c=a+"="+b+"; path="+c+"; ";g&&(c+="expires="+(new Date((new Date).getTime()+g)).toGMTString()+"; ");d&&"none"!==d&&(c+="domain="+d+";");ca&&(c+=ca+";");d=M.cookie;M.cookie=c;if(!(d=d!=M.cookie))a:{a=
Ca(a);for(d=0;d<a.length;d++)if(b==a[d]){d=!0;break a}d=!1}return d},Cc=function(a){return encodeURIComponent?encodeURIComponent(a).replace(/\(/g,"%28").replace(/\)/g,"%29"):a},vc=/^(www\.)?google(\.com?)?(\.[a-z]{2})?$/,eb=/(^|\.)doubleclick\.net$/i;var Fa,Ga,fb,Ab,ja=/^https?:\/\/[^/]*cdn\.ampproject\.org\//,Ue=/^(?:www\.|m\.|amp\.)+/,Ub=[],da=function(a){if(ye(a[Kd])){if(void 0===Ab){var b;if(b=(b=De.get())&&b._ga||void 0)Ab=b,J(81)}if(void 0!==Ab)return a[Q]||(a[Q]=Ab),!1}if(a[Kd]){J(67);if(a[ac]&&"cookie"!=a[ac])return!1;if(void 0!==Ab)a[Q]||(a[Q]=Ab);else{a:{b=String(a[W]||xa());var c=String(a[Yb]||"/"),d=Ca(String(a[U]||"_ga"));b=na(d,b,c);if(!b||jd.test(b))b=!0;else if(b=Ca("AMP_TOKEN"),0==b.length)b=!0;else{if(1==b.length&&(b=decodeURIComponent(b[0]),
"$RETRIEVING"==b||"$OPT_OUT"==b||"$ERROR"==b||"$NOT_FOUND"==b)){b=!0;break a}b=!1}}if(b&&tc(ic,String(a[Na])))return!0}}return!1},ic=function(){Z.D([ua])},tc=function(a,b){var c=Ca("AMP_TOKEN");if(1<c.length)return J(55),!1;c=decodeURIComponent(c[0]||"");if("$OPT_OUT"==c||"$ERROR"==c||G(b))return J(62),!1;if(!ja.test(M.referrer)&&"$NOT_FOUND"==c)return J(68),!1;if(void 0!==Ab)return J(56),va(function(){a(Ab)},0),!0;if(Fa)return Ub.push(a),!0;if("$RETRIEVING"==c)return J(57),va(function(){tc(a,b)},
1E4),!0;Fa=!0;c&&"$"!=c[0]||(xc("$RETRIEVING",3E4),setTimeout(Mc,3E4),c="");return Pc(c,b)?(Ub.push(a),!0):!1},Pc=function(a,b,c){if(!window.JSON)return J(58),!1;var d=O.XMLHttpRequest;if(!d)return J(59),!1;var e=new d;if(!("withCredentials"in e))return J(60),!1;e.open("POST",(c||"https://ampcid.google.com/v1/publisher:getClientId")+"?key=AIzaSyA65lEHUEizIsNtlbNo-l2K18dT680nsaM",!0);e.withCredentials=!0;e.setRequestHeader("Content-Type","text/plain");e.onload=function(){Fa=!1;if(4==e.readyState){try{200!=
e.status&&(J(61),Qc("","$ERROR",3E4));var g=JSON.parse(e.responseText);g.optOut?(J(63),Qc("","$OPT_OUT",31536E6)):g.clientId?Qc(g.clientId,g.securityToken,31536E6):!c&&g.alternateUrl?(Ga&&clearTimeout(Ga),Fa=!0,Pc(a,b,g.alternateUrl)):(J(64),Qc("","$NOT_FOUND",36E5))}catch(ca){J(65),Qc("","$ERROR",3E4)}e=null}};d={originScope:"AMP_ECID_GOOGLE"};a&&(d.securityToken=a);e.send(JSON.stringify(d));Ga=va(function(){J(66);Qc("","$ERROR",3E4)},1E4);return!0},Mc=function(){Fa=!1},xc=function(a,b){if(void 0===
fb){fb="";for(var c=id(),d=0;d<c.length;d++){var e=c[d];if(zc("AMP_TOKEN",encodeURIComponent(a),"/",e,"",b)){fb=e;return}}}zc("AMP_TOKEN",encodeURIComponent(a),"/",fb,"",b)},Qc=function(a,b,c){Ga&&clearTimeout(Ga);b&&xc(b,c);Ab=a;b=Ub;Ub=[];for(c=0;c<b.length;c++)b[c](a)},ye=function(a){a:{if(ja.test(M.referrer)){var b=M.location.hostname.replace(Ue,"");b:{var c=M.referrer;c=c.replace(/^https?:\/\//,"");var d=c.replace(/^[^/]+/,"").split("/"),e=d[2];d=(d="s"==e?d[3]:e)?decodeURIComponent(d):d;if(!d){if(0==
c.indexOf("xn--")){c="";break b}(c=c.match(/(.*)\.cdn\.ampproject\.org\/?$/))&&2==c.length&&(d=c[1].replace(/-/g,".").replace(/\.\./g,"-"))}c=d?d.replace(Ue,""):""}(d=b===c)||(c="."+c,d=b.substring(b.length-c.length,b.length)===c);if(d){b=!0;break a}else J(78)}b=!1}return b&&!1!==a};var bd=function(a){return(a?"https:":Ba||"https:"==M.location.protocol?"https:":"http:")+"//www.google-analytics.com"},Ge=function(a){switch(a){default:case 1:return"https://www.google-analytics.com/gtm/js?id=";case 2:return"https://www.googletagmanager.com/gtag/js?id="}},Da=function(a){this.name="len";this.message=a+"-8192"},ba=function(a,b,c){c=c||ua;if(2036>=b.length)wc(a,b,c);else if(8192>=b.length)x(a,b,c)||wd(a,b,c)||wc(a,b,c);else throw ge("len",b.length),new Da(b.length);},pe=function(a,b,
c,d){d=d||ua;wd(a+"?"+b,"",d,c)},wc=function(a,b,c){var d=ta(a+"?"+b);d.onload=d.onerror=function(){d.onload=null;d.onerror=null;c()}},wd=function(a,b,c,d){var e=O.XMLHttpRequest;if(!e)return!1;var g=new e;if(!("withCredentials"in g))return!1;a=a.replace(/^http:/,"https:");g.open("POST",a,!0);g.withCredentials=!0;g.setRequestHeader("Content-Type","text/plain");g.onreadystatechange=function(){if(4==g.readyState){if(d&&"text/plain"===g.getResponseHeader("Content-Type"))try{Ea(d,g.responseText,c)}catch(ca){ge("xhr",
"rsp"),c()}else c();g=null}};g.send(b);return!0},Ea=function(a,b,c){if(1>b.length)ge("xhr","ver","0"),c();else if(3<a.count++)ge("xhr","tmr",""+a.count),c();else{var d=b.charAt(0);if("1"===d)oc(a,b.substring(1),c);else if(a.V&&"2"===d){var e=b.substring(1).split(","),g=0;b=function(){++g===e.length&&c()};for(d=0;d<e.length;d++)oc(a,e[d],b)}else ge("xhr","ver",String(b.length)),c()}},oc=function(a,b,c){if(0===b.length)c();else{var d=b.charAt(0);switch(d){case "d":pe("https://stats.g.doubleclick.net/j/collect",
a.U,a,c);break;case "g":wc("https://www.google.%/ads/ga-audiences".replace("%","com"),a.google,c);(b=b.substring(1))&&(/^[a-z.]{1,6}$/.test(b)?wc("https://www.google.%/ads/ga-audiences".replace("%",b),a.google,ua):ge("tld","bcc",b));break;case "G":if(a.V){a.V("G-"+b.substring(1));c();break}case "x":if(a.V){a.V();c();break}default:ge("xhr","brc",d),c()}}},x=function(a,b,c){return O.navigator.sendBeacon?O.navigator.sendBeacon(a,b)?(c(),!0):!1:!1},ge=function(a,b,c){1<=100*Math.random()||G("?")||(a=
["t=error","_e="+a,"_v=j85","sr=1"],b&&a.push("_f="+b),c&&a.push("_m="+K(c.substring(0,100))),a.push("aip=1"),a.push("z="+hd()),wc(bd(!0)+"/u/d",a.join("&"),ua))};var qc=function(){return O.gaData=O.gaData||{}},h=function(a){var b=qc();return b[a]=b[a]||{}};var Ha=function(){this.M=[]};Ha.prototype.add=function(a){this.M.push(a)};Ha.prototype.D=function(a){try{for(var b=0;b<this.M.length;b++){var c=a.get(this.M[b]);c&&ea(c)&&c.call(O,a)}}catch(d){}b=a.get(Ia);b!=ua&&ea(b)&&(a.set(Ia,ua,!0),setTimeout(b,10))};function Ja(a){if(100!=a.get(Ka)&&La(P(a,Q))%1E4>=100*R(a,Ka))throw"abort";}function Ma(a){if(G(P(a,Na)))throw"abort";}function Oa(){var a=M.location.protocol;if("http:"!=a&&"https:"!=a)throw"abort";}
function Pa(a){try{O.navigator.sendBeacon?J(42):O.XMLHttpRequest&&"withCredentials"in new O.XMLHttpRequest&&J(40)}catch(c){}a.set(ld,Td(a),!0);a.set(Ac,R(a,Ac)+1);var b=[];ue.map(function(c,d){d.F&&(c=a.get(c),void 0!=c&&c!=d.defaultValue&&("boolean"==typeof c&&(c*=1),b.push(d.F+"="+K(""+c))))});!1===a.get(xe)&&b.push("npa=1");b.push("z="+Bd());a.set(Ra,b.join("&"),!0)}
function Sa(a){var b=P(a,fa);!b&&a.get(Vd)&&(b="beacon");var c=P(a,gd),d=P(a,oe),e=c||(d||bd(!1)+"")+"/collect";switch(P(a,ad)){case "d":e=c||(d||bd(!1)+"")+"/j/collect";b=a.get(qe)||void 0;pe(e,P(a,Ra),b,a.Z(Ia));break;default:b?(c=P(a,Ra),d=(d=a.Z(Ia))||ua,"image"==b?wc(e,c,d):"xhr"==b&&wd(e,c,d)||"beacon"==b&&x(e,c,d)||ba(e,c,d)):ba(e,P(a,Ra),a.Z(Ia))}e=P(a,Na);e=h(e);b=e.hitcount;e.hitcount=b?b+1:1;e.first_hit||(e.first_hit=(new Date).getTime());e=P(a,Na);delete h(e).pending_experiments;a.set(Ia,
ua,!0)}function Hc(a){qc().expId&&a.set(Nc,qc().expId);qc().expVar&&a.set(Oc,qc().expVar);var b=P(a,Na);if(b=h(b).pending_experiments){var c=[];for(d in b)b.hasOwnProperty(d)&&b[d]&&c.push(encodeURIComponent(d)+"."+encodeURIComponent(b[d]));var d=c.join("!")}else d=void 0;d&&((b=a.get(m))&&(d=b+"!"+d),a.set(m,d,!0))}function cd(){if(O.navigator&&"preview"==O.navigator.loadPurpose)throw"abort";}
function yd(a){var b=O.gaDevIds||[];if(ka(b)){var c=a.get("&did");qa(c)&&0<c.length&&(b=b.concat(c.split(",")));c=[];for(var d=0;d<b.length;d++){var e;a:{for(e=0;e<c.length;e++)if(b[d]==c[e]){e=!0;break a}e=!1}e||c.push(b[d])}0!=c.length&&a.set("&did",c.join(","),!0)}}function vb(a){if(!a.get(Na))throw"abort";};var hd=function(){return Math.round(2147483647*Math.random())},Bd=function(){try{var a=new Uint32Array(1);O.crypto.getRandomValues(a);return a[0]&2147483647}catch(b){return hd()}};function Ta(a){var b=R(a,Ua);500<=b&&J(15);var c=P(a,Va);if("transaction"!=c&&"item"!=c){c=R(a,Wa);var d=(new Date).getTime(),e=R(a,Xa);0==e&&a.set(Xa,d);e=Math.round(2*(d-e)/1E3);0<e&&(c=Math.min(c+e,20),a.set(Xa,d));if(0>=c)throw"abort";a.set(Wa,--c)}a.set(Ua,++b)};var Ya=function(){this.data=new ee};Ya.prototype.get=function(a){var b=$a(a),c=this.data.get(a);b&&void 0==c&&(c=ea(b.defaultValue)?b.defaultValue():b.defaultValue);return b&&b.Z?b.Z(this,a,c):c};var P=function(a,b){a=a.get(b);return void 0==a?"":""+a},R=function(a,b){a=a.get(b);return void 0==a||""===a?0:Number(a)};Ya.prototype.Z=function(a){return(a=this.get(a))&&ea(a)?a:ua};
Ya.prototype.set=function(a,b,c){if(a)if("object"==typeof a)for(var d in a)a.hasOwnProperty(d)&&ab(this,d,a[d],c);else ab(this,a,b,c)};var ab=function(a,b,c,d){if(void 0!=c)switch(b){case Na:wb.test(c)}var e=$a(b);e&&e.o?e.o(a,b,c,d):a.data.set(b,c,d)};var ue=new ee,ve=[],bb=function(a,b,c,d,e){this.name=a;this.F=b;this.Z=d;this.o=e;this.defaultValue=c},$a=function(a){var b=ue.get(a);if(!b)for(var c=0;c<ve.length;c++){var d=ve[c],e=d[0].exec(a);if(e){b=d[1](e);ue.set(b.name,b);break}}return b},yc=function(a){var b;ue.map(function(c,d){d.F==a&&(b=d)});return b&&b.name},S=function(a,b,c,d,e){a=new bb(a,b,c,d,e);ue.set(a.name,a);return a.name},cb=function(a,b){ve.push([new RegExp("^"+a+"$"),b])},T=function(a,b,c){return S(a,b,c,void 0,db)},db=function(){};var hb=T("apiVersion","v"),ib=T("clientVersion","_v");S("anonymizeIp","aip");var jb=S("adSenseId","a"),Va=S("hitType","t"),Ia=S("hitCallback"),Ra=S("hitPayload");S("nonInteraction","ni");S("currencyCode","cu");S("dataSource","ds");var Vd=S("useBeacon",void 0,!1),fa=S("transport");S("sessionControl","sc","");S("sessionGroup","sg");S("queueTime","qt");var Ac=S("_s","_s");S("screenName","cd");var kb=S("location","dl",""),lb=S("referrer","dr"),mb=S("page","dp","");S("hostname","dh");
var nb=S("language","ul"),ob=S("encoding","de");S("title","dt",function(){return M.title||void 0});cb("contentGroup([0-9]+)",function(a){return new bb(a[0],"cg"+a[1])});var pb=S("screenColors","sd"),qb=S("screenResolution","sr"),rb=S("viewportSize","vp"),sb=S("javaEnabled","je"),tb=S("flashVersion","fl");S("campaignId","ci");S("campaignName","cn");S("campaignSource","cs");S("campaignMedium","cm");S("campaignKeyword","ck");S("campaignContent","cc");
var ub=S("eventCategory","ec"),xb=S("eventAction","ea"),yb=S("eventLabel","el"),zb=S("eventValue","ev"),Bb=S("socialNetwork","sn"),Cb=S("socialAction","sa"),Db=S("socialTarget","st"),Eb=S("l1","plt"),Fb=S("l2","pdt"),Gb=S("l3","dns"),Hb=S("l4","rrt"),Ib=S("l5","srt"),Jb=S("l6","tcp"),Kb=S("l7","dit"),Lb=S("l8","clt"),Ve=S("l9","_gst"),We=S("l10","_gbt"),Xe=S("l11","_cst"),Ye=S("l12","_cbt"),Mb=S("timingCategory","utc"),Nb=S("timingVar","utv"),Ob=S("timingLabel","utl"),Pb=S("timingValue","utt");
S("appName","an");S("appVersion","av","");S("appId","aid","");S("appInstallerId","aiid","");S("exDescription","exd");S("exFatal","exf");var Nc=S("expId","xid"),Oc=S("expVar","xvar"),m=S("exp","exp");S("_slexp",void 0);var Rc=S("_utma","_utma"),Sc=S("_utmz","_utmz"),Tc=S("_utmht","_utmht"),Ua=S("_hc",void 0,0),Xa=S("_ti",void 0,0),Wa=S("_to",void 0,20);cb("dimension([0-9]+)",function(a){return new bb(a[0],"cd"+a[1])});cb("metric([0-9]+)",function(a){return new bb(a[0],"cm"+a[1])});
S("linkerParam",void 0,void 0,Bc,db);var Ze=T("_cd2l",void 0,!1),ld=S("usage","_u"),Gd=S("_um");S("forceSSL",void 0,void 0,function(){return Ba},function(a,b,c){J(34);Ba=!!c});var ed=S("_j1","jid"),ia=S("_j2","gjid");cb("\\&(.*)",function(a){var b=new bb(a[0],a[1]),c=yc(a[0].substring(1));c&&(b.Z=function(d){return d.get(c)},b.o=function(d,e,g,ca){d.set(c,g,ca)},b.F=void 0);return b});
var Qb=T("_oot"),dd=S("previewTask"),Rb=S("checkProtocolTask"),md=S("validationTask"),Sb=S("checkStorageTask"),Uc=S("historyImportTask"),Tb=S("samplerTask"),Vb=S("_rlt"),Wb=S("buildHitTask"),Xb=S("sendHitTask"),Vc=S("ceTask"),zd=S("devIdTask"),Cd=S("timingTask"),Ld=S("displayFeaturesTask"),oa=S("customTask"),ze=S("fpsCrossDomainTask"),V=T("name"),Q=T("clientId","cid"),n=T("clientIdTime"),xd=T("storedClientId"),Ad=S("userId","uid"),Na=T("trackingId","tid"),U=T("cookieName",void 0,"_ga"),W=T("cookieDomain"),
Yb=T("cookiePath",void 0,"/"),Zb=T("cookieExpires",void 0,63072E3),Hd=T("cookieUpdate",void 0,!0),Be=T("cookieFlags",void 0,""),$b=T("legacyCookieDomain"),Wc=T("legacyHistoryImport",void 0,!0),ac=T("storage",void 0,"cookie"),bc=T("allowLinker",void 0,!1),cc=T("allowAnchor",void 0,!0),Ka=T("sampleRate","sf",100),dc=T("siteSpeedSampleRate",void 0,1),ec=T("alwaysSendReferrer",void 0,!1),I=T("_gid","_gid"),la=T("_gcn"),Kd=T("useAmpClientId"),ce=T("_gclid"),fe=T("_gt"),he=T("_ge",void 0,7776E6),ie=T("_gclsrc"),
je=T("storeGac",void 0,!0),oe=S("_x_19"),Ae=S("_fplc","_fplc"),gd=S("transportUrl"),Md=S("_r","_r");S("_slc","_slc");var qe=S("_dp"),ad=S("_jt",void 0,"n"),Ud=S("allowAdFeatures",void 0,!0),xe=S("allowAdPersonalizationSignals",void 0,!0);function X(a,b,c,d){b[a]=function(){try{return d&&J(d),c.apply(this,arguments)}catch(e){throw ge("exc",a,e&&e.name),e;}}};function fc(){var a,b;if((b=(b=O.navigator)?b.plugins:null)&&b.length)for(var c=0;c<b.length&&!a;c++){var d=b[c];-1<d.name.indexOf("Shockwave Flash")&&(a=d.description)}if(!a)try{var e=new ActiveXObject("ShockwaveFlash.ShockwaveFlash.7");a=e.GetVariable("$version")}catch(g){}if(!a)try{e=new ActiveXObject("ShockwaveFlash.ShockwaveFlash.6"),a="WIN 6,0,21,0",e.AllowScriptAccess="always",a=e.GetVariable("$version")}catch(g){}if(!a)try{e=new ActiveXObject("ShockwaveFlash.ShockwaveFlash"),a=e.GetVariable("$version")}catch(g){}a&&
(e=a.match(/[\d]+/g))&&3<=e.length&&(a=e[0]+"."+e[1]+" r"+e[2]);return a||void 0};var Ed=function(a){if("cookie"==a.get(ac))return a=Ca("FPLC"),0<a.length?a[0]:void 0},Fe=function(a){var b;if(b=P(a,oe)&&a.get(Ze))b=De.get(a.get(cc)),b=!(b&&b._fplc);b&&a.set(Ae,Ed(a)||"0")};var aa=function(a){var b=Math.min(R(a,dc),100);return La(P(a,Q))%100>=b?!1:!0},gc=function(a){var b={};if(Ec(b)||Fc(b)){var c=b[Eb];void 0==c||Infinity==c||isNaN(c)||(0<c?(Y(b,Gb),Y(b,Jb),Y(b,Ib),Y(b,Fb),Y(b,Hb),Y(b,Kb),Y(b,Lb),Y(b,Ve),Y(b,We),Y(b,Xe),Y(b,Ye),va(function(){a(b)},10)):L(O,"load",function(){gc(a)},!1))}},Ec=function(a){var b=O.performance||O.webkitPerformance;b=b&&b.timing;if(!b)return!1;var c=b.navigationStart;if(0==c)return!1;a[Eb]=b.loadEventStart-c;a[Gb]=b.domainLookupEnd-b.domainLookupStart;
a[Jb]=b.connectEnd-b.connectStart;a[Ib]=b.responseStart-b.requestStart;a[Fb]=b.responseEnd-b.responseStart;a[Hb]=b.fetchStart-c;a[Kb]=b.domInteractive-c;a[Lb]=b.domContentLoadedEventStart-c;a[Ve]=N.L-c;a[We]=N.ya-c;O.google_tag_manager&&O.google_tag_manager._li&&(b=O.google_tag_manager._li,a[Xe]=b.cst,a[Ye]=b.cbt);return!0},Fc=function(a){if(O.top!=O)return!1;var b=O.external,c=b&&b.onloadT;b&&!b.isValidLoadTime&&(c=void 0);2147483648<c&&(c=void 0);0<c&&b.setPageReadyTime();if(void 0==c)return!1;
a[Eb]=c;return!0},Y=function(a,b){var c=a[b];if(isNaN(c)||Infinity==c||0>c)a[b]=void 0},Fd=function(a){return function(b){if("pageview"==b.get(Va)&&!a.I){a.I=!0;var c=aa(b),d=0<E(P(b,kb),"gclid").length;(c||d)&&gc(function(e){c&&a.send("timing",e);d&&a.send("adtiming",e)})}}};var hc=!1,mc=function(a){if("cookie"==P(a,ac)){if(a.get(Hd)||P(a,xd)!=P(a,Q)){var b=1E3*R(a,Zb);ma(a,Q,U,b);a.data.set(xd,P(a,Q))}(a.get(Hd)||uc(a)!=P(a,I))&&ma(a,I,la,864E5);if(a.get(je)){var c=P(a,ce);if(c){var d=Math.min(R(a,he),1E3*R(a,Zb));d=Math.min(d,1E3*R(a,fe)+d-(new Date).getTime());a.data.set(he,d);b={};var e=P(a,fe),g=P(a,ie),ca=kc(P(a,Yb)),l=lc(P(a,W)),k=P(a,Na);a=P(a,Be);g&&"aw.ds"!=g?b&&(b.ua=!0):(c=["1",e,Cc(c)].join("."),0<d&&(b&&(b.ta=!0),zc("_gac_"+Cc(k),c,ca,l,k,d,a)));le(b)}}else J(75)}},
ma=function(a,b,c,d){var e=nd(a,b);if(e){c=P(a,c);var g=kc(P(a,Yb)),ca=lc(P(a,W)),l=P(a,Be),k=P(a,Na);if("auto"!=ca)zc(c,e,g,ca,k,d,l)&&(hc=!0);else{J(32);for(var w=id(),Ce=0;Ce<w.length;Ce++)if(ca=w[Ce],a.data.set(W,ca),e=nd(a,b),zc(c,e,g,ca,k,d,l)){hc=!0;return}a.data.set(W,"auto")}}},uc=function(a){var b=Ca(P(a,la));return Xd(a,b)},nc=function(a){if("cookie"==P(a,ac)&&!hc&&(mc(a),!hc))throw"abort";},Yc=function(a){if(a.get(Wc)){var b=P(a,W),c=P(a,$b)||xa(),d=Xc("__utma",c,b);d&&(J(19),a.set(Tc,
(new Date).getTime(),!0),a.set(Rc,d.R),(b=Xc("__utmz",c,b))&&d.hash==b.hash&&a.set(Sc,b.R))}},nd=function(a,b){b=Cc(P(a,b));var c=lc(P(a,W)).split(".").length;a=jc(P(a,Yb));1<a&&(c+="-"+a);return b?["GA1",c,b].join("."):""},Xd=function(a,b){return na(b,P(a,W),P(a,Yb))},na=function(a,b,c){if(!a||1>a.length)J(12);else{for(var d=[],e=0;e<a.length;e++){var g=a[e];var ca=g.split(".");var l=ca.shift();("GA1"==l||"1"==l)&&1<ca.length?(g=ca.shift().split("-"),1==g.length&&(g[1]="1"),g[0]*=1,g[1]*=1,ca={H:g,
s:ca.join(".")}):ca=kd.test(g)?{H:[0,0],s:g}:void 0;ca&&d.push(ca)}if(1==d.length)return J(13),d[0].s;if(0==d.length)J(12);else{J(14);d=Gc(d,lc(b).split(".").length,0);if(1==d.length)return d[0].s;d=Gc(d,jc(c),1);1<d.length&&J(41);return d[0]&&d[0].s}}},Gc=function(a,b,c){for(var d=[],e=[],g,ca=0;ca<a.length;ca++){var l=a[ca];l.H[c]==b?d.push(l):void 0==g||l.H[c]<g?(e=[l],g=l.H[c]):l.H[c]==g&&e.push(l)}return 0<d.length?d:e},lc=function(a){return 0==a.indexOf(".")?a.substr(1):a},id=function(){var a=
[],b=xa().split(".");if(4==b.length){var c=b[b.length-1];if(parseInt(c,10)==c)return["none"]}for(c=b.length-2;0<=c;c--)a.push(b.slice(c).join("."));b=M.location.hostname;eb.test(b)||vc.test(b)||a.push("none");return a},kc=function(a){if(!a)return"/";1<a.length&&a.lastIndexOf("/")==a.length-1&&(a=a.substr(0,a.length-1));0!=a.indexOf("/")&&(a="/"+a);return a},jc=function(a){a=kc(a);return"/"==a?1:a.split("/").length},le=function(a){a.ta&&J(77);a.na&&J(74);a.pa&&J(73);a.ua&&J(69)};function Xc(a,b,c){"none"==b&&(b="");var d=[],e=Ca(a);a="__utma"==a?6:2;for(var g=0;g<e.length;g++){var ca=(""+e[g]).split(".");ca.length>=a&&d.push({hash:ca[0],R:e[g],O:ca})}if(0!=d.length)return 1==d.length?d[0]:Zc(b,d)||Zc(c,d)||Zc(null,d)||d[0]}function Zc(a,b){if(null==a)var c=a=1;else c=La(a),a=La(D(a,".")?a.substring(1):"."+a);for(var d=0;d<b.length;d++)if(b[d].hash==c||b[d].hash==a)return b[d]};var Jc=new RegExp(/^https?:\/\/([^\/:]+)/),De=O.google_tag_data.glBridge,Kc=/(.*)([?&#])(?:_ga=[^&#]*)(?:&?)(.*)/,od=/(.*)([?&#])(?:_gac=[^&#]*)(?:&?)(.*)/;function Bc(a){if(a.get(Ze))return J(35),De.generate($e(a));var b=P(a,Q),c=P(a,I)||"";b="_ga=2."+K(pa(c+b,0)+"."+c+"-"+b);(a=af(a))?(J(44),a="&_gac=1."+K([pa(a.qa,0),a.timestamp,a.qa].join("."))):a="";return b+a}
function Ic(a,b){var c=new Date,d=O.navigator,e=d.plugins||[];a=[a,d.userAgent,c.getTimezoneOffset(),c.getYear(),c.getDate(),c.getHours(),c.getMinutes()+b];for(b=0;b<e.length;++b)a.push(e[b].description);return La(a.join("."))}function pa(a,b){var c=new Date,d=O.navigator,e=c.getHours()+Math.floor((c.getMinutes()+b)/60);return La([a,d.userAgent,d.language||"",c.getTimezoneOffset(),c.getYear(),c.getDate()+Math.floor(e/24),(24+e)%24,(60+c.getMinutes()+b)%60].join("."))}
var Dc=function(a){J(48);this.target=a;this.T=!1};Dc.prototype.ca=function(a,b){if(a){if(this.target.get(Ze))return De.decorate($e(this.target),a,b);if(a.tagName){if("a"==a.tagName.toLowerCase()){a.href&&(a.href=qd(this,a.href,b));return}if("form"==a.tagName.toLowerCase())return rd(this,a)}if("string"==typeof a)return qd(this,a,b)}};
var qd=function(a,b,c){var d=Kc.exec(b);d&&3<=d.length&&(b=d[1]+(d[3]?d[2]+d[3]:""));(d=od.exec(b))&&3<=d.length&&(b=d[1]+(d[3]?d[2]+d[3]:""));a=a.target.get("linkerParam");var e=b.indexOf("?");d=b.indexOf("#");c?b+=(-1==d?"#":"&")+a:(c=-1==e?"?":"&",b=-1==d?b+(c+a):b.substring(0,d)+c+a+b.substring(d));b=b.replace(/&+_ga=/,"&_ga=");return b=b.replace(/&+_gac=/,"&_gac=")},rd=function(a,b){if(b&&b.action)if("get"==b.method.toLowerCase()){a=a.target.get("linkerParam").split("&");for(var c=0;c<a.length;c++){var d=
a[c].split("="),e=d[1];d=d[0];for(var g=b.childNodes||[],ca=!1,l=0;l<g.length;l++)if(g[l].name==d){g[l].setAttribute("value",e);ca=!0;break}ca||(g=M.createElement("input"),g.setAttribute("type","hidden"),g.setAttribute("name",d),g.setAttribute("value",e),b.appendChild(g))}}else"post"==b.method.toLowerCase()&&(b.action=qd(a,b.action))};
Dc.prototype.S=function(a,b,c){function d(g){try{g=g||O.event;a:{var ca=g.target||g.srcElement;for(g=100;ca&&0<g;){if(ca.href&&ca.nodeName.match(/^a(?:rea)?$/i)){var l=ca;break a}ca=ca.parentNode;g--}l={}}("http:"==l.protocol||"https:"==l.protocol)&&sd(a,l.hostname||"")&&l.href&&(l.href=qd(e,l.href,b))}catch(k){J(26)}}var e=this;this.target.get(Ze)?De.auto(function(){return $e(e.target)},a,b?"fragment":"",c):(this.T||(this.T=!0,L(M,"mousedown",d,!1),L(M,"keyup",d,!1)),c&&L(M,"submit",function(g){g=
g||O.event;if((g=g.target||g.srcElement)&&g.action){var ca=g.action.match(Jc);ca&&sd(a,ca[1])&&rd(e,g)}}))};function sd(a,b){if(b==M.location.hostname)return!1;for(var c=0;c<a.length;c++)if(a[c]instanceof RegExp){if(a[c].test(b))return!0}else if(0<=b.indexOf(a[c]))return!0;return!1}function ke(a,b){return b!=Ic(a,0)&&b!=Ic(a,-1)&&b!=Ic(a,-2)&&b!=pa(a,0)&&b!=pa(a,-1)&&b!=pa(a,-2)}
function $e(a){var b=af(a),c={};c._ga=a.get(Q);c._gid=a.get(I)||void 0;c._gac=b?[b.qa,b.timestamp].join("."):void 0;b=a.get(Ae);a=Ed(a);return c._fplc=b&&"0"!==b?b:a,c}function af(a){function b(e){return void 0==e||""===e?0:Number(e)}var c=a.get(ce);if(c&&a.get(je)){var d=b(a.get(fe));if(1E3*d+b(a.get(he))<=(new Date).getTime())J(76);else return{timestamp:d,qa:c}}};var p=/^(GTM|OPT)-[A-Z0-9]+$/,q=/;_gaexp=[^;]*/g,r=/;((__utma=)|([^;=]+=GAX?\d+\.))[^;]*/g,Aa=/^https?:\/\/[\w\-.]+\.google.com(:\d+)?\/optimize\/opt-launch\.html\?.*$/,t=function(a){function b(d,e){e&&(c+="&"+d+"="+K(e))}var c=Ge(a.type)+K(a.id);"dataLayer"!=a.B&&b("l",a.B);b("cx",a.context);b("t",a.target);b("cid",a.clientId);b("cidt",a.ka);b("gac",a.la);b("aip",a.ia);a.sync&&b("m","sync");b("cycle",a.G);a.qa&&b("gclid",a.qa);Aa.test(M.referrer)&&b("cb",String(hd()));return c},He=function(a){var b=
void 0,c=(new Date).getTime();O[a.B]=O[a.B]||[];c={"gtm.start":c};a.sync||(c.event="gtm.js");O[a.B].push(c);2===a.type&&function(d,e,g){O[a.B].push(arguments)}("config",a.id,b)};var Jd=function(a,b){b||(b=(b=P(a,V))&&"t0"!=b?Wd.test(b)?"_gat_"+Cc(P(a,Na)):"_gat_"+Cc(b):"_gat");this.Y=b},Rd=function(a,b){var c=b.get(Wb);b.set(Wb,function(e){Pd(a,e,ed);Pd(a,e,ia);var g=c(e);Qd(a,e);return g});var d=b.get(Xb);b.set(Xb,function(e){var g=d(e);if(se(e)){J(80);var ca={U:re(e,1),google:re(e,2),count:0};pe("https://stats.g.doubleclick.net/j/collect",ca.U,ca);e.set(ed,"",!0)}return g})},Pd=function(a,b,c){!1===b.get(Ud)||b.get(c)||("1"==Ca(a.Y)[0]?b.set(c,"",!0):b.set(c,""+hd(),!0))},
Qd=function(a,b){se(b)&&zc(a.Y,"1",P(b,Yb),P(b,W),P(b,Na),6E4,P(b,Be))},se=function(a){return!!a.get(ed)&&!1!==a.get(Ud)},re=function(a,b){var c=new ee,d=function(g){$a(g).F&&c.set($a(g).F,a.get(g))};d(hb);d(ib);d(Na);d(Q);d(ed);1==b&&(d(Ad),d(ia),d(I));c.set($a(ld).F,Td(a));var e="";c.map(function(g,ca){e+=K(g)+"=";e+=K(""+ca)+"&"});e+="z="+hd();1==b?e="t=dc&aip=1&_r=3&"+e:2==b&&(e="t=sr&aip=1&_r=4&slf_rd=1&"+e);return e},Wd=/^gtm\d+$/;var fd=function(a,b){a=a.model;if(!a.get("dcLoaded")){var c=new $c(Dd(a));c.set(29);a.set(Gd,c.C);b=b||{};var d;b[U]&&(d=Cc(b[U]));b=new Jd(a,d);Rd(b,a);a.set("dcLoaded",!0)}};var Sd=function(a){if(!a.get("dcLoaded")&&"cookie"==a.get(ac)){var b=new Jd(a);Pd(b,a,ed);Pd(b,a,ia);Qd(b,a);se(a)&&(a.set(Md,1,!0),a.set(ad,"d",!0),J(79),a.set(qe,{U:re(a,1),google:re(a,2),V:void 0,count:0},!0))}};var Lc=function(){var a=O.gaGlobal=O.gaGlobal||{};return a.hid=a.hid||hd()};var wb=/^(UA|YT|MO|GP)-(\d+)-(\d+)$/,pc=function(a){function b(e,g){d.model.data.set(e,g)}function c(e,g){b(e,g);d.filters.add(e)}var d=this;this.model=new Ya;this.filters=new Ha;b(V,a[V]);b(Na,sa(a[Na]));b(U,a[U]);b(W,a[W]||xa());b(Yb,a[Yb]);b(Zb,a[Zb]);b(Hd,a[Hd]);b(Be,a[Be]);b($b,a[$b]);b(Wc,a[Wc]);b(bc,a[bc]);b(cc,a[cc]);b(Ka,a[Ka]);b(dc,a[dc]);b(ec,a[ec]);b(ac,a[ac]);b(Ad,a[Ad]);b(n,a[n]);b(Kd,a[Kd]);b(je,a[je]);b(Ze,a[Ze]);b(oe,a[oe]);b(hb,1);b(ib,"j85");c(Qb,Ma);c(oa,ua);c(dd,cd);c(Rb,Oa);
c(md,vb);c(Sb,nc);c(Uc,Yc);c(Tb,Ja);c(Vb,Ta);c(Vc,Hc);c(zd,yd);c(Ld,Sd);c(ze,Fe);c(Wb,Pa);c(Xb,Sa);c(Cd,Fd(this));pd(this.model);td(this.model,a[Q]);this.model.set(jb,Lc())};pc.prototype.get=function(a){return this.model.get(a)};pc.prototype.set=function(a,b){this.model.set(a,b)};
pc.prototype.send=function(a){if(!(1>arguments.length)){if("string"===typeof arguments[0]){var b=arguments[0];var c=[].slice.call(arguments,1)}else b=arguments[0]&&arguments[0][Va],c=arguments;b&&(c=za(me[b]||[],c),c[Va]=b,this.model.set(c,void 0,!0),this.filters.D(this.model),this.model.data.m={})}};pc.prototype.ma=function(a,b){var c=this;u(a,c,b)||(v(a,function(){u(a,c,b)}),y(String(c.get(V)),a,void 0,b,!0))};
var td=function(a,b){var c=P(a,U);a.data.set(la,"_ga"==c?"_gid":c+"_gid");if("cookie"==P(a,ac)){hc=!1;c=Ca(P(a,U));c=Xd(a,c);if(!c){c=P(a,W);var d=P(a,$b)||xa();c=Xc("__utma",d,c);void 0!=c?(J(10),c=c.O[1]+"."+c.O[2]):c=void 0}c&&(hc=!0);if(d=c&&!a.get(Hd))if(d=c.split("."),2!=d.length)d=!1;else if(d=Number(d[1])){var e=R(a,Zb);d=d+e<(new Date).getTime()/1E3}else d=!1;d&&(c=void 0);c&&(a.data.set(xd,c),a.data.set(Q,c),(c=uc(a))&&a.data.set(I,c));if(a.get(je)&&(c=a.get(ce),d=a.get(ie),!c||d&&"aw.ds"!=
d)){c={};if(M){d=[];e=M.cookie.split(";");for(var g=/^\s*_gac_(UA-\d+-\d+)=\s*(.+?)\s*$/,ca=0;ca<e.length;ca++){var l=e[ca].match(g);l&&d.push({ja:l[1],value:l[2]})}e={};if(d&&d.length)for(g=0;g<d.length;g++)(ca=d[g].value.split("."),"1"!=ca[0]||3!=ca.length)?c&&(c.na=!0):ca[1]&&(e[d[g].ja]?c&&(c.pa=!0):e[d[g].ja]=[],e[d[g].ja].push({timestamp:ca[1],qa:ca[2]}));d=e}else d={};d=d[P(a,Na)];le(c);d&&0!=d.length&&(c=d[0],a.data.set(fe,c.timestamp),a.data.set(ce,c.qa))}}if(a.get(Hd)&&(c=be("_ga",!!a.get(cc)),
g=be("_gl",!!a.get(cc)),d=De.get(a.get(cc)),e=d._ga,g&&0<g.indexOf("_ga*")&&!e&&J(30),g=d.gclid,ca=d._gac,c||e||g||ca))if(c&&e&&J(36),a.get(bc)||ye(a.get(Kd))){e&&(J(38),a.data.set(Q,e),d._gid&&(J(51),a.data.set(I,d._gid)));g?(J(82),a.data.set(ce,g),d.gclsrc&&a.data.set(ie,d.gclsrc)):ca&&(e=ca.split("."))&&2===e.length&&(J(37),a.data.set(ce,e[0]),a.data.set(fe,e[1]));if(d=d._fplc)J(83),a.data.set(Ae,d);if(c)b:if(d=c.indexOf("."),-1==d)J(22);else{e=c.substring(0,d);g=c.substring(d+1);d=g.indexOf(".");
c=g.substring(0,d);g=g.substring(d+1);if("1"==e){if(d=g,ke(d,c)){J(23);break b}}else if("2"==e){d=g.indexOf("-");e="";0<d?(e=g.substring(0,d),d=g.substring(d+1)):d=g.substring(1);if(ke(e+d,c)){J(53);break b}e&&(J(2),a.data.set(I,e))}else{J(22);break b}J(11);a.data.set(Q,d);if(c=be("_gac",!!a.get(cc)))c=c.split("."),"1"!=c[0]||4!=c.length?J(72):ke(c[3],c[1])?J(71):(a.data.set(ce,c[3]),a.data.set(fe,c[2]),J(70))}}else J(21);b&&(J(9),a.data.set(Q,K(b)));a.get(Q)||(b=(b=O.gaGlobal)&&b.from_cookie&&"cookie"!==
P(a,ac)?void 0:(b=b&&b.vid)&&-1!==b.search(jd)?b:void 0,b?(J(17),a.data.set(Q,b)):(J(8),a.data.set(Q,ra())));a.get(I)||(J(3),a.data.set(I,ra()));mc(a);b=O.gaGlobal=O.gaGlobal||{};c=P(a,Q);a=c===P(a,xd);if(void 0==b.vid||a&&!b.from_cookie)b.vid=c,b.from_cookie=a},pd=function(a){var b=O.navigator,c=O.screen,d=M.location;a.set(lb,ya(!!a.get(ec),!!a.get(Kd)));if(d){var e=d.pathname||"";"/"!=e.charAt(0)&&(J(31),e="/"+e);a.set(kb,d.protocol+"//"+d.hostname+e+d.search)}c&&a.set(qb,c.width+"x"+c.height);
c&&a.set(pb,c.colorDepth+"-bit");c=M.documentElement;var g=(e=M.body)&&e.clientWidth&&e.clientHeight,ca=[];c&&c.clientWidth&&c.clientHeight&&("CSS1Compat"===M.compatMode||!g)?ca=[c.clientWidth,c.clientHeight]:g&&(ca=[e.clientWidth,e.clientHeight]);c=0>=ca[0]||0>=ca[1]?"":ca.join("x");a.set(rb,c);a.set(tb,fc());a.set(ob,M.characterSet||M.charset);a.set(sb,b&&"function"===typeof b.javaEnabled&&b.javaEnabled()||!1);a.set(nb,(b&&(b.language||b.browserLanguage)||"").toLowerCase());a.data.set(ce,be("gclid",
!0));a.data.set(ie,be("gclsrc",!0));a.data.set(fe,Math.round((new Date).getTime()/1E3));if(d&&a.get(cc)&&(b=M.location.hash)){b=b.split(/[?&#]+/);d=[];for(c=0;c<b.length;++c)(D(b[c],"utm_id")||D(b[c],"utm_campaign")||D(b[c],"utm_source")||D(b[c],"utm_medium")||D(b[c],"utm_term")||D(b[c],"utm_content")||D(b[c],"gclid")||D(b[c],"dclid")||D(b[c],"gclsrc"))&&d.push(b[c]);0<d.length&&(b="#"+d.join("&"),a.set(kb,a.get(kb)+b))}},me={pageview:[mb],event:[ub,xb,yb,zb],social:[Bb,Cb,Db],timing:[Mb,Nb,Pb,Ob]};var rc=function(a){if("prerender"==M.visibilityState)return!1;a();return!0},z=function(a){if(!rc(a)){J(16);var b=!1,c=function(){if(!b&&rc(a)){b=!0;var d=c,e=M;e.removeEventListener?e.removeEventListener("visibilitychange",d,!1):e.detachEvent&&e.detachEvent("onvisibilitychange",d)}};L(M,"visibilitychange",c)}};var te=/^(?:(\w+)\.)?(?:(\w+):)?(\w+)$/,sc=function(a){if(ea(a[0]))this.u=a[0];else{var b=te.exec(a[0]);null!=b&&4==b.length&&(this.c=b[1]||"t0",this.K=b[2]||"",this.methodName=b[3],this.a=[].slice.call(a,1),this.K||(this.A="create"==this.methodName,this.i="require"==this.methodName,this.g="provide"==this.methodName,this.ba="remove"==this.methodName),this.i&&(3<=this.a.length?(this.X=this.a[1],this.W=this.a[2]):this.a[1]&&(qa(this.a[1])?this.X=this.a[1]:this.W=this.a[1])));b=a[1];a=a[2];if(!this.methodName)throw"abort";
if(this.i&&(!qa(b)||""==b))throw"abort";if(this.g&&(!qa(b)||""==b||!ea(a)))throw"abort";if(ud(this.c)||ud(this.K))throw"abort";if(this.g&&"t0"!=this.c)throw"abort";}};function ud(a){return 0<=a.indexOf(".")||0<=a.indexOf(":")};var Yd,Zd,$d,A;Yd=new ee;$d=new ee;A=new ee;Zd={ec:45,ecommerce:46,linkid:47};
var u=function(a,b,c){b==N||b.get(V);var d=Yd.get(a);if(!ea(d))return!1;b.plugins_=b.plugins_||new ee;if(b.plugins_.get(a))return!0;b.plugins_.set(a,new d(b,c||{}));return!0},y=function(a,b,c,d,e){if(!ea(Yd.get(b))&&!$d.get(b)){Zd.hasOwnProperty(b)&&J(Zd[b]);a=N.j(a);if(p.test(b)){J(52);if(!a)return!0;c=a.model;var g=d||{};d={id:b,type:1,B:g.dataLayer||"dataLayer",G:!1};c.get("&gtm")==b&&(d.G=!0);d.ia=!!c.get("anonymizeIp");d.sync=e;var ca=String(c.get("name"));"t0"!=ca&&(d.target=ca);G(String(c.get("trackingId")))||
(d.clientId=String(c.get(Q)),d.ka=Number(c.get(n)),g=g.palindrome?r:q,g=(g=M.cookie.replace(/^|(; +)/g,";").match(g))?g.sort().join("").substring(1):void 0,d.la=g,d.qa=E(P(c,kb),"gclid"));He(d);c=t(d)}!c&&Zd.hasOwnProperty(b)?(J(39),c=b+".js"):J(43);if(c){if(a){var l=a.get(oe);qa(l)||(l=void 0)}c&&0<=c.indexOf("/")||(c=(l||bd(!1))+"/plugins/ua/"+c);l=ae(c);a=l.protocol;c=M.location.protocol;if(("https:"==a||a==c||("http:"!=a?0:"http:"==c))&&B(l)){if(l=l.url)a=(a=M.querySelector&&M.querySelector("script[nonce]")||
null)?a.nonce||a.getAttribute&&a.getAttribute("nonce")||"":"",e?(e="",a&&Nd.test(a)&&(e=' nonce="'+a+'"'),f.test(l)&&M.write("<script"+e+' src="'+l+'">\x3c/script>')):(e=M.createElement("script"),e.type="text/javascript",e.async=!0,e.src=l,a&&e.setAttribute("nonce",a),l=M.getElementsByTagName("script")[0],l.parentNode.insertBefore(e,l));$d.set(b,!0)}}}},v=function(a,b){var c=A.get(a)||[];c.push(b);A.set(a,c)},C=function(a,b){Yd.set(a,b);b=A.get(a)||[];for(var c=0;c<b.length;c++)b[c]();A.set(a,[])},
B=function(a){var b=ae(M.location.href);if(D(a.url,Ge(1))||D(a.url,Ge(2)))return!0;if(a.query||0<=a.url.indexOf("?")||0<=a.path.indexOf("://"))return!1;if(a.host==b.host&&a.port==b.port)return!0;b="http:"==a.protocol?80:443;return"www.google-analytics.com"==a.host&&(a.port||b)==b&&D(a.path,"/plugins/")?!0:!1},ae=function(a){function b(l){var k=l.hostname||"",w=0<=k.indexOf("]");k=k.split(w?"]":":")[0].toLowerCase();w&&(k+="]");w=(l.protocol||"").toLowerCase();w=1*l.port||("http:"==w?80:"https:"==
w?443:"");l=l.pathname||"";D(l,"/")||(l="/"+l);return[k,""+w,l]}var c=M.createElement("a");c.href=M.location.href;var d=(c.protocol||"").toLowerCase(),e=b(c),g=c.search||"",ca=d+"//"+e[0]+(e[1]?":"+e[1]:"");D(a,"//")?a=d+a:D(a,"/")?a=ca+a:!a||D(a,"?")?a=ca+e[2]+(a||g):0>a.split("/")[0].indexOf(":")&&(a=ca+e[2].substring(0,e[2].lastIndexOf("/"))+"/"+a);c.href=a;d=b(c);return{protocol:(c.protocol||"").toLowerCase(),host:d[0],port:d[1],path:d[2],query:c.search||"",url:a||""}};var Z={ga:function(){Z.f=[]}};Z.ga();Z.D=function(a){var b=Z.J.apply(Z,arguments);b=Z.f.concat(b);for(Z.f=[];0<b.length&&!Z.v(b[0])&&!(b.shift(),0<Z.f.length););Z.f=Z.f.concat(b)};Z.J=function(a){for(var b=[],c=0;c<arguments.length;c++)try{var d=new sc(arguments[c]);d.g?C(d.a[0],d.a[1]):(d.i&&(d.ha=y(d.c,d.a[0],d.X,d.W)),b.push(d))}catch(e){}return b};
Z.v=function(a){try{if(a.u)a.u.call(O,N.j("t0"));else{var b=a.c==gb?N:N.j(a.c);if(a.A){if("t0"==a.c&&(b=N.create.apply(N,a.a),null===b))return!0}else if(a.ba)N.remove(a.c);else if(b)if(a.i){if(a.ha&&(a.ha=y(a.c,a.a[0],a.X,a.W)),!u(a.a[0],b,a.W))return!0}else if(a.K){var c=a.methodName,d=a.a,e=b.plugins_.get(a.K);e[c].apply(e,d)}else b[a.methodName].apply(b,a.a)}}catch(g){}};var N=function(a){J(1);Z.D.apply(Z,[arguments])};N.h={};N.P=[];N.L=0;N.ya=0;N.answer=42;var we=[Na,W,V];N.create=function(a){var b=za(we,[].slice.call(arguments));b[V]||(b[V]="t0");var c=""+b[V];if(N.h[c])return N.h[c];if(da(b))return null;b=new pc(b);N.h[c]=b;N.P.push(b);c=qc().tracker_created;if(ea(c))try{c(b)}catch(d){}return b};N.remove=function(a){for(var b=0;b<N.P.length;b++)if(N.P[b].get(V)==a){N.P.splice(b,1);N.h[a]=null;break}};N.j=function(a){return N.h[a]};N.getAll=function(){return N.P.slice(0)};
N.N=function(){"ga"!=gb&&J(49);var a=O[gb];if(!a||42!=a.answer){N.L=a&&a.l;N.ya=1*new Date;N.loaded=!0;var b=O[gb]=N;X("create",b,b.create);X("remove",b,b.remove);X("getByName",b,b.j,5);X("getAll",b,b.getAll,6);b=pc.prototype;X("get",b,b.get,7);X("set",b,b.set,4);X("send",b,b.send);X("requireSync",b,b.ma);b=Ya.prototype;X("get",b,b.get);X("set",b,b.set);if("https:"!=M.location.protocol&&!Ba){a:{b=M.getElementsByTagName("script");for(var c=0;c<b.length&&100>c;c++){var d=b[c].src;if(d&&0==d.indexOf(bd(!0)+
"/analytics")){b=!0;break a}}b=!1}b&&(Ba=!0)}(O.gaplugins=O.gaplugins||{}).Linker=Dc;b=Dc.prototype;C("linker",Dc);X("decorate",b,b.ca,20);X("autoLink",b,b.S,25);C("displayfeatures",fd);C("adfeatures",fd);a=a&&a.q;ka(a)?Z.D.apply(N,a):J(50)}};N.da=function(){for(var a=N.getAll(),b=0;b<a.length;b++)a[b].get(V)};var Id=N.N,Od=O[gb];Od&&Od.r?Id():z(Id);z(function(){Z.D(["provide","render",ua])});})(window);
```
