## Supplementary File 2 for "Single individual structural variant detection uncovers widespread hemizygosity in molluscs": bootstrap.js

```
/*!
 * Bootstrap v3.2.0 (http://getbootstrap.com)
 * Copyright 2011-2014 Twitter, Inc.
 * Licensed under MIT (https://github.com/twbs/bootstrap/blob/master/LICENSE)
 */
if("undefined"==typeof jQuery)throw new Error("Bootstrap's JavaScript requires jQuery");+function(a){"use strict";function b(){var a=document.createElement("bootstrap"),b={WebkitTransition:"webkitTransitionEnd",MozTransition:"transitionend",OTransition:"oTransitionEnd otransitionend",transition:"transitionend"};for(var c in b)if(void 0!==a.style[c])return{end:b[c]};return!1}a.fn.emulateTransitionEnd=function(b){var c=!1,d=this;a(this).one("bsTransitionEnd",function(){c=!0});var e=function(){c||a(d).trigger(a.support.transition.end)};return setTimeout(e,b),this},a(function(){a.support.transition=b(),a.support.transition&&(a.event.special.bsTransitionEnd={bindType:a.support.transition.end,delegateType:a.support.transition.end,handle:function(b){return a(b.target).is(this)?b.handleObj.handler.apply(this,arguments):void 0}})})}(jQuery),+function(a){"use strict";function b(b){return this.each(function(){var c=a(this),e=c.data("bs.alert");e||c.data("bs.alert",e=new d(this)),"string"==typeof b&&e[b].call(c)})}var c='[data-dismiss="alert"]',d=function(b){a(b).on("click",c,this.close)};d.VERSION="3.2.0",d.prototype.close=function(b){function c(){f.detach().trigger("closed.bs.alert").remove()}var d=a(this),e=d.attr("data-target");e||(e=d.attr("href"),e=e&&e.replace(/.*(?=#[^\s]*$)/,""));var f=a(e);b&&b.preventDefault(),f.length||(f=d.hasClass("alert")?d:d.parent()),f.trigger(b=a.Event("close.bs.alert")),b.isDefaultPrevented()||(f.removeClass("in"),a.support.transition&&f.hasClass("fade")?f.one("bsTransitionEnd",c).emulateTransitionEnd(150):c())};var e=a.fn.alert;a.fn.alert=b,a.fn.alert.Constructor=d,a.fn.alert.noConflict=function(){return a.fn.alert=e,this},a(document).on("click.bs.alert.data-api",c,d.prototype.close)}(jQuery),+function(a){"use strict";function b(b){return this.each(function(){var d=a(this),e=d.data("bs.button"),f="object"==typeof b&&b;e||d.data("bs.button",e=new c(this,f)),"toggle"==b?e.toggle():b&&e.setState(b)})}var c=function(b,d){this.$element=a(b),this.options=a.extend({},c.DEFAULTS,d),this.isLoading=!1};c.VERSION="3.2.0",c.DEFAULTS={loadingText:"loading..."},c.prototype.setState=function(b){var c="disabled",d=this.$element,e=d.is("input")?"val":"html",f=d.data();b+="Text",null==f.resetText&&d.data("resetText",d[e]()),d[e](null==f[b]?this.options[b]:f[b]),setTimeout(a.proxy(function(){"loadingText"==b?(this.isLoading=!0,d.addClass(c).attr(c,c)):this.isLoading&&(this.isLoading=!1,d.removeClass(c).removeAttr(c))},this),0)},c.prototype.toggle=function(){var a=!0,b=this.$element.closest('[data-toggle="buttons"]');if(b.length){var c=this.$element.find("input");"radio"==c.prop("type")&&(c.prop("checked")&&this.$element.hasClass("active")?a=!1:b.find(".active").removeClass("active")),a&&c.prop("checked",!this.$element.hasClass("active")).trigger("change")}a&&this.$element.toggleClass("active")};var d=a.fn.button;a.fn.button=b,a.fn.button.Constructor=c,a.fn.button.noConflict=function(){return a.fn.button=d,this},a(document).on("click.bs.button.data-api",'[data-toggle^="button"]',function(c){var d=a(c.target);d.hasClass("btn")||(d=d.closest(".btn")),b.call(d,"toggle"),c.preventDefault()})}(jQuery),+function(a){"use strict";function b(b){return this.each(function(){var d=a(this),e=d.data("bs.carousel"),f=a.extend({},c.DEFAULTS,d.data(),"object"==typeof b&&b),g="string"==typeof b?b:f.slide;e||d.data("bs.carousel",e=new c(this,f)),"number"==typeof b?e.to(b):g?e[g]():f.interval&&e.pause().cycle()})}var c=function(b,c){this.$element=a(b).on("keydown.bs.carousel",a.proxy(this.keydown,this)),this.$indicators=this.$element.find(".carousel-indicators"),this.options=c,this.paused=this.sliding=this.interval=this.$active=this.$items=null,"hover"==this.options.pause&&this.$element.on("mouseenter.bs.carousel",a.proxy(this.pause,this)).on("mouseleave.bs.carousel",a.proxy(this.cycle,this))};c.VERSION="3.2.0",c.DEFAULTS={interval:5e3,pause:"hover",wrap:!0},c.prototype.keydown=function(a){switch(a.which){case 37:this.prev();break;case 39:this.next();break;default:return}a.preventDefault()},c.prototype.cycle=function(b){return b||(this.paused=!1),this.interval&&clearInterval(this.interval),this.options.interval&&!this.paused&&(this.interval=setInterval(a.proxy(this.next,this),this.options.interval)),this},c.prototype.getItemIndex=function(a){return this.$items=a.parent().children(".item"),this.$items.index(a||this.$active)},c.prototype.to=function(b){var c=this,d=this.getItemIndex(this.$active=this.$element.find(".item.active"));return b>this.$items.length-1||0>b?void 0:this.sliding?this.$element.one("slid.bs.carousel",function(){c.to(b)}):d==b?this.pause().cycle():this.slide(b>d?"next":"prev",a(this.$items[b]))},c.prototype.pause=function(b){return b||(this.paused=!0),this.$element.find(".next, .prev").length&&a.support.transition&&(this.$element.trigger(a.support.transition.end),this.cycle(!0)),this.interval=clearInterval(this.interval),this},c.prototype.next=function(){return this.sliding?void 0:this.slide("next")},c.prototype.prev=function(){return this.sliding?void 0:this.slide("prev")},c.prototype.slide=function(b,c){var d=this.$element.find(".item.active"),e=c||d[b](),f=this.interval,g="next"==b?"left":"right",h="next"==b?"first":"last",i=this;if(!e.length){if(!this.options.wrap)return;e=this.$element.find(".item")[h]()}if(e.hasClass("active"))return this.sliding=!1;var j=e[0],k=a.Event("slide.bs.carousel",{relatedTarget:j,direction:g});if(this.$element.trigger(k),!k.isDefaultPrevented()){if(this.sliding=!0,f&&this.pause(),this.$indicators.length){this.$indicators.find(".active").removeClass("active");var l=a(this.$indicators.children()[this.getItemIndex(e)]);l&&l.addClass("active")}var m=a.Event("slid.bs.carousel",{relatedTarget:j,direction:g});return a.support.transition&&this.$element.hasClass("slide")?(e.addClass(b),e[0].offsetWidth,d.addClass(g),e.addClass(g),d.one("bsTransitionEnd",function(){e.removeClass([b,g].join(" ")).addClass("active"),d.removeClass(["active",g].join(" ")),i.sliding=!1,setTimeout(function(){i.$element.trigger(m)},0)}).emulateTransitionEnd(1e3*d.css("transition-duration").slice(0,-1))):(d.removeClass("active"),e.addClass("active"),this.sliding=!1,this.$element.trigger(m)),f&&this.cycle(),this}};var d=a.fn.carousel;a.fn.carousel=b,a.fn.carousel.Constructor=c,a.fn.carousel.noConflict=function(){return a.fn.carousel=d,this},a(document).on("click.bs.carousel.data-api","[data-slide], [data-slide-to]",function(c){var d,e=a(this),f=a(e.attr("data-target")||(d=e.attr("href"))&&d.replace(/.*(?=#[^\s]+$)/,""));if(f.hasClass("carousel")){var g=a.extend({},f.data(),e.data()),h=e.attr("data-slide-to");h&&(g.interval=!1),b.call(f,g),h&&f.data("bs.carousel").to(h),c.preventDefault()}}),a(window).on("load",function(){a('[data-ride="carousel"]').each(function(){var c=a(this);b.call(c,c.data())})})}(jQuery),+function(a){"use strict";function b(b){return this.each(function(){var d=a(this),e=d.data("bs.collapse"),f=a.extend({},c.DEFAULTS,d.data(),"object"==typeof b&&b);!e&&f.toggle&&"show"==b&&(b=!b),e||d.data("bs.collapse",e=new c(this,f)),"string"==typeof b&&e[b]()})}var c=function(b,d){this.$element=a(b),this.options=a.extend({},c.DEFAULTS,d),this.transitioning=null,this.options.parent&&(this.$parent=a(this.options.parent)),this.options.toggle&&this.toggle()};c.VERSION="3.2.0",c.DEFAULTS={toggle:!0},c.prototype.dimension=function(){var a=this.$element.hasClass("width");return a?"width":"height"},c.prototype.show=function(){if(!this.transitioning&&!this.$element.hasClass("in")){var c=a.Event("show.bs.collapse");if(this.$element.trigger(c),!c.isDefaultPrevented()){var d=this.$parent&&this.$parent.find("> .panel > .in");if(d&&d.length){var e=d.data("bs.collapse");if(e&&e.transitioning)return;b.call(d,"hide"),e||d.data("bs.collapse",null)}var f=this.dimension();this.$element.removeClass("collapse").addClass("collapsing")[f](0),this.transitioning=1;var g=function(){this.$element.removeClass("collapsing").addClass("collapse in")[f](""),this.transitioning=0,this.$element.trigger("shown.bs.collapse")};if(!a.support.transition)return g.call(this);var h=a.camelCase(["scroll",f].join("-"));this.$element.one("bsTransitionEnd",a.proxy(g,this)).emulateTransitionEnd(350)[f](this.$element[0][h])}}},c.prototype.hide=function(){if(!this.transitioning&&this.$element.hasClass("in")){var b=a.Event("hide.bs.collapse");if(this.$element.trigger(b),!b.isDefaultPrevented()){var c=this.dimension();this.$element[c](this.$element[c]())[0].offsetHeight,this.$element.addClass("collapsing").removeClass("collapse").removeClass("in"),this.transitioning=1;var d=function(){this.transitioning=0,this.$element.trigger("hidden.bs.collapse").removeClass("collapsing").addClass("collapse")};return a.support.transition?void this.$element[c](0).one("bsTransitionEnd",a.proxy(d,this)).emulateTransitionEnd(350):d.call(this)}}},c.prototype.toggle=function(){this[this.$element.hasClass("in")?"hide":"show"]()};var d=a.fn.collapse;a.fn.collapse=b,a.fn.collapse.Constructor=c,a.fn.collapse.noConflict=function(){return a.fn.collapse=d,this},a(document).on("click.bs.collapse.data-api",'[data-toggle="collapse"]',function(c){var d,e=a(this),f=e.attr("data-target")||c.preventDefault()||(d=e.attr("href"))&&d.replace(/.*(?=#[^\s]+$)/,""),g=a(f),h=g.data("bs.collapse"),i=h?"toggle":e.data(),j=e.attr("data-parent"),k=j&&a(j);h&&h.transitioning||(k&&k.find('[data-toggle="collapse"][data-parent="'+j+'"]').not(e).addClass("collapsed"),e[g.hasClass("in")?"addClass":"removeClass"]("collapsed")),b.call(g,i)})}(jQuery),+function(a){"use strict";function b(b){b&&3===b.which||(a(e).remove(),a(f).each(function(){var d=c(a(this)),e={relatedTarget:this};d.hasClass("open")&&(d.trigger(b=a.Event("hide.bs.dropdown",e)),b.isDefaultPrevented()||d.removeClass("open").trigger("hidden.bs.dropdown",e))}))}function c(b){var c=b.attr("data-target");c||(c=b.attr("href"),c=c&&/#[A-Za-z]/.test(c)&&c.replace(/.*(?=#[^\s]*$)/,""));var d=c&&a(c);return d&&d.length?d:b.parent()}function d(b){return this.each(function(){var c=a(this),d=c.data("bs.dropdown");d||c.data("bs.dropdown",d=new g(this)),"string"==typeof b&&d[b].call(c)})}var e=".dropdown-backdrop",f='[data-toggle="dropdown"]',g=function(b){a(b).on("click.bs.dropdown",this.toggle)};g.VERSION="3.2.0",g.prototype.toggle=function(d){var e=a(this);if(!e.is(".disabled, :disabled")){var f=c(e),g=f.hasClass("open");if(b(),!g){"ontouchstart"in document.documentElement&&!f.closest(".navbar-nav").length&&a('<div class="dropdown-backdrop"/>').insertAfter(a(this)).on("click",b);var h={relatedTarget:this};if(f.trigger(d=a.Event("show.bs.dropdown",h)),d.isDefaultPrevented())return;e.trigger("focus"),f.toggleClass("open").trigger("shown.bs.dropdown",h)}return!1}},g.prototype.keydown=function(b){if(/(38|40|27)/.test(b.keyCode)){var d=a(this);if(b.preventDefault(),b.stopPropagation(),!d.is(".disabled, :disabled")){var e=c(d),g=e.hasClass("open");if(!g||g&&27==b.keyCode)return 27==b.which&&e.find(f).trigger("focus"),d.trigger("click");var h=" li:not(.divider):visible a",i=e.find('[role="menu"]'+h+', [role="listbox"]'+h);if(i.length){var j=i.index(i.filter(":focus"));38==b.keyCode&&j>0&&j--,40==b.keyCode&&j<i.length-1&&j++,~j||(j=0),i.eq(j).trigger("focus")}}}};var h=a.fn.dropdown;a.fn.dropdown=d,a.fn.dropdown.Constructor=g,a.fn.dropdown.noConflict=function(){return a.fn.dropdown=h,this},a(document).on("click.bs.dropdown.data-api",b).on("click.bs.dropdown.data-api",".dropdown form",function(a){a.stopPropagation()}).on("click.bs.dropdown.data-api",f,g.prototype.toggle).on("keydown.bs.dropdown.data-api",f+', [role="menu"], [role="listbox"]',g.prototype.keydown)}(jQuery),+function(a){"use strict";function b(b,d){return this.each(function(){var e=a(this),f=e.data("bs.modal"),g=a.extend({},c.DEFAULTS,e.data(),"object"==typeof b&&b);f||e.data("bs.modal",f=new c(this,g)),"string"==typeof b?f[b](d):g.show&&f.show(d)})}var c=function(b,c){this.options=c,this.$body=a(document.body),this.$element=a(b),this.$backdrop=this.isShown=null,this.scrollbarWidth=0,this.options.remote&&this.$element.find(".modal-content").load(this.options.remote,a.proxy(function(){this.$element.trigger("loaded.bs.modal")},this))};c.VERSION="3.2.0",c.DEFAULTS={backdrop:!0,keyboard:!0,show:!0},c.prototype.toggle=function(a){return this.isShown?this.hide():this.show(a)},c.prototype.show=function(b){var c=this,d=a.Event("show.bs.modal",{relatedTarget:b});this.$element.trigger(d),this.isShown||d.isDefaultPrevented()||(this.isShown=!0,this.checkScrollbar(),this.$body.addClass("modal-open"),this.setScrollbar(),this.escape(),this.$element.on("click.dismiss.bs.modal",'[data-dismiss="modal"]',a.proxy(this.hide,this)),this.backdrop(function(){var d=a.support.transition&&c.$element.hasClass("fade");c.$element.parent().length||c.$element.appendTo(c.$body),c.$element.show().scrollTop(0),d&&c.$element[0].offsetWidth,c.$element.addClass("in").attr("aria-hidden",!1),c.enforceFocus();var e=a.Event("shown.bs.modal",{relatedTarget:b});d?c.$element.find(".modal-dialog").one("bsTransitionEnd",function(){c.$element.trigger("focus").trigger(e)}).emulateTransitionEnd(300):c.$element.trigger("focus").trigger(e)}))},c.prototype.hide=function(b){b&&b.preventDefault(),b=a.Event("hide.bs.modal"),this.$element.trigger(b),this.isShown&&!b.isDefaultPrevented()&&(this.isShown=!1,this.$body.removeClass("modal-open"),this.resetScrollbar(),this.escape(),a(document).off("focusin.bs.modal"),this.$element.removeClass("in").attr("aria-hidden",!0).off("click.dismiss.bs.modal"),a.support.transition&&this.$element.hasClass("fade")?this.$element.one("bsTransitionEnd",a.proxy(this.hideModal,this)).emulateTransitionEnd(300):this.hideModal())},c.prototype.enforceFocus=function(){a(document).off("focusin.bs.modal").on("focusin.bs.modal",a.proxy(function(a){this.$element[0]===a.target||this.$element.has(a.target).length||this.$element.trigger("focus")},this))},c.prototype.escape=function(){this.isShown&&this.options.keyboard?this.$element.on("keyup.dismiss.bs.modal",a.proxy(function(a){27==a.which&&this.hide()},this)):this.isShown||this.$element.off("keyup.dismiss.bs.modal")},c.prototype.hideModal=function(){var a=this;this.$element.hide(),this.backdrop(function(){a.$element.trigger("hidden.bs.modal")})},c.prototype.removeBackdrop=function(){this.$backdrop&&this.$backdrop.remove(),this.$backdrop=null},c.prototype.backdrop=function(b){var c=this,d=this.$element.hasClass("fade")?"fade":"";if(this.isShown&&this.options.backdrop){var e=a.support.transition&&d;if(this.$backdrop=a('<div class="modal-backdrop '+d+'" />').appendTo(this.$body),this.$element.on("click.dismiss.bs.modal",a.proxy(function(a){a.target===a.currentTarget&&("static"==this.options.backdrop?this.$element[0].focus.call(this.$element[0]):this.hide.call(this))},this)),e&&this.$backdrop[0].offsetWidth,this.$backdrop.addClass("in"),!b)return;e?this.$backdrop.one("bsTransitionEnd",b).emulateTransitionEnd(150):b()}else if(!this.isShown&&this.$backdrop){this.$backdrop.removeClass("in");var f=function(){c.removeBackdrop(),b&&b()};a.support.transition&&this.$element.hasClass("fade")?this.$backdrop.one("bsTransitionEnd",f).emulateTransitionEnd(150):f()}else b&&b()},c.prototype.checkScrollbar=function(){document.body.clientWidth>=window.innerWidth||(this.scrollbarWidth=this.scrollbarWidth||this.measureScrollbar())},c.prototype.setScrollbar=function(){var a=parseInt(this.$body.css("padding-right")||0,10);this.scrollbarWidth&&this.$body.css("padding-right",a+this.scrollbarWidth)},c.prototype.resetScrollbar=function(){this.$body.css("padding-right","")},c.prototype.measureScrollbar=function(){var a=document.createElement("div");a.className="modal-scrollbar-measure",this.$body.append(a);var b=a.offsetWidth-a.clientWidth;return this.$body[0].removeChild(a),b};var d=a.fn.modal;a.fn.modal=b,a.fn.modal.Constructor=c,a.fn.modal.noConflict=function(){return a.fn.modal=d,this},a(document).on("click.bs.modal.data-api",'[data-toggle="modal"]',function(c){var d=a(this),e=d.attr("href"),f=a(d.attr("data-target")||e&&e.replace(/.*(?=#[^\s]+$)/,"")),g=f.data("bs.modal")?"toggle":a.extend({remote:!/#/.test(e)&&e},f.data(),d.data());d.is("a")&&c.preventDefault(),f.one("show.bs.modal",function(a){a.isDefaultPrevented()||f.one("hidden.bs.modal",function(){d.is(":visible")&&d.trigger("focus")})}),b.call(f,g,this)})}(jQuery),+function(a){"use strict";function b(b){return this.each(function(){var d=a(this),e=d.data("bs.tooltip"),f="object"==typeof b&&b;(e||"destroy"!=b)&&(e||d.data("bs.tooltip",e=new c(this,f)),"string"==typeof b&&e[b]())})}var c=function(a,b){this.type=this.options=this.enabled=this.timeout=this.hoverState=this.$element=null,this.init("tooltip",a,b)};c.VERSION="3.2.0",c.DEFAULTS={animation:!0,placement:"top",selector:!1,template:'<div class="tooltip" role="tooltip"><div class="tooltip-arrow"></div><div class="tooltip-inner"></div></div>',trigger:"hover focus",title:"",delay:0,html:!1,container:!1,viewport:{selector:"body",padding:0}},c.prototype.init=function(b,c,d){this.enabled=!0,this.type=b,this.$element=a(c),this.options=this.getOptions(d),this.$viewport=this.options.viewport&&a(this.options.viewport.selector||this.options.viewport);for(var e=this.options.trigger.split(" "),f=e.length;f--;){var g=e[f];if("click"==g)this.$element.on("click."+this.type,this.options.selector,a.proxy(this.toggle,this));else if("manual"!=g){var h="hover"==g?"mouseenter":"focusin",i="hover"==g?"mouseleave":"focusout";this.$element.on(h+"."+this.type,this.options.selector,a.proxy(this.enter,this)),this.$element.on(i+"."+this.type,this.options.selector,a.proxy(this.leave,this))}}this.options.selector?this._options=a.extend({},this.options,{trigger:"manual",selector:""}):this.fixTitle()},c.prototype.getDefaults=function(){return c.DEFAULTS},c.prototype.getOptions=function(b){return b=a.extend({},this.getDefaults(),this.$element.data(),b),b.delay&&"number"==typeof b.delay&&(b.delay={show:b.delay,hide:b.delay}),b},c.prototype.getDelegateOptions=function(){var b={},c=this.getDefaults();return this._options&&a.each(this._options,function(a,d){c[a]!=d&&(b[a]=d)}),b},c.prototype.enter=function(b){var c=b instanceof this.constructor?b:a(b.currentTarget).data("bs."+this.type);return c||(c=new this.constructor(b.currentTarget,this.getDelegateOptions()),a(b.currentTarget).data("bs."+this.type,c)),clearTimeout(c.timeout),c.hoverState="in",c.options.delay&&c.options.delay.show?void(c.timeout=setTimeout(function(){"in"==c.hoverState&&c.show()},c.options.delay.show)):c.show()},c.prototype.leave=function(b){var c=b instanceof this.constructor?b:a(b.currentTarget).data("bs."+this.type);return c||(c=new this.constructor(b.currentTarget,this.getDelegateOptions()),a(b.currentTarget).data("bs."+this.type,c)),clearTimeout(c.timeout),c.hoverState="out",c.options.delay&&c.options.delay.hide?void(c.timeout=setTimeout(function(){"out"==c.hoverState&&c.hide()},c.options.delay.hide)):c.hide()},c.prototype.show=function(){var b=a.Event("show.bs."+this.type);if(this.hasContent()&&this.enabled){this.$element.trigger(b);var c=a.contains(document.documentElement,this.$element[0]);if(b.isDefaultPrevented()||!c)return;var d=this,e=this.tip(),f=this.getUID(this.type);this.setContent(),e.attr("id",f),this.$element.attr("aria-describedby",f),this.options.animation&&e.addClass("fade");var g="function"==typeof this.options.placement?this.options.placement.call(this,e[0],this.$element[0]):this.options.placement,h=/\s?auto?\s?/i,i=h.test(g);i&&(g=g.replace(h,"")||"top"),e.detach().css({top:0,left:0,display:"block"}).addClass(g).data("bs."+this.type,this),this.options.container?e.appendTo(this.options.container):e.insertAfter(this.$element);var j=this.getPosition(),k=e[0].offsetWidth,l=e[0].offsetHeight;if(i){var m=g,n=this.$element.parent(),o=this.getPosition(n);g="bottom"==g&&j.top+j.height+l-o.scroll>o.height?"top":"top"==g&&j.top-o.scroll-l<0?"bottom":"right"==g&&j.right+k>o.width?"left":"left"==g&&j.left-k<o.left?"right":g,e.removeClass(m).addClass(g)}var p=this.getCalculatedOffset(g,j,k,l);this.applyPlacement(p,g);var q=function(){d.$element.trigger("shown.bs."+d.type),d.hoverState=null};a.support.transition&&this.$tip.hasClass("fade")?e.one("bsTransitionEnd",q).emulateTransitionEnd(150):q()}},c.prototype.applyPlacement=function(b,c){var d=this.tip(),e=d[0].offsetWidth,f=d[0].offsetHeight,g=parseInt(d.css("margin-top"),10),h=parseInt(d.css("margin-left"),10);isNaN(g)&&(g=0),isNaN(h)&&(h=0),b.top=b.top+g,b.left=b.left+h,a.offset.setOffset(d[0],a.extend({using:function(a){d.css({top:Math.round(a.top),left:Math.round(a.left)})}},b),0),d.addClass("in");var i=d[0].offsetWidth,j=d[0].offsetHeight;"top"==c&&j!=f&&(b.top=b.top+f-j);var k=this.getViewportAdjustedDelta(c,b,i,j);k.left?b.left+=k.left:b.top+=k.top;var l=k.left?2*k.left-e+i:2*k.top-f+j,m=k.left?"left":"top",n=k.left?"offsetWidth":"offsetHeight";d.offset(b),this.replaceArrow(l,d[0][n],m)},c.prototype.replaceArrow=function(a,b,c){this.arrow().css(c,a?50*(1-a/b)+"%":"")},c.prototype.setContent=function(){var a=this.tip(),b=this.getTitle();a.find(".tooltip-inner")[this.options.html?"html":"text"](b),a.removeClass("fade in top bottom left right")},c.prototype.hide=function(){function b(){"in"!=c.hoverState&&d.detach(),c.$element.trigger("hidden.bs."+c.type)}var c=this,d=this.tip(),e=a.Event("hide.bs."+this.type);return this.$element.removeAttr("aria-describedby"),this.$element.trigger(e),e.isDefaultPrevented()?void 0:(d.removeClass("in"),a.support.transition&&this.$tip.hasClass("fade")?d.one("bsTransitionEnd",b).emulateTransitionEnd(150):b(),this.hoverState=null,this)},c.prototype.fixTitle=function(){var a=this.$element;(a.attr("title")||"string"!=typeof a.attr("data-original-title"))&&a.attr("data-original-title",a.attr("title")||"").attr("title","")},c.prototype.hasContent=function(){return this.getTitle()},c.prototype.getPosition=function(b){b=b||this.$element;var c=b[0],d="BODY"==c.tagName;return a.extend({},"function"==typeof c.getBoundingClientRect?c.getBoundingClientRect():null,{scroll:d?document.documentElement.scrollTop||document.body.scrollTop:b.scrollTop(),width:d?a(window).width():b.outerWidth(),height:d?a(window).height():b.outerHeight()},d?{top:0,left:0}:b.offset())},c.prototype.getCalculatedOffset=function(a,b,c,d){return"bottom"==a?{top:b.top+b.height,left:b.left+b.width/2-c/2}:"top"==a?{top:b.top-d,left:b.left+b.width/2-c/2}:"left"==a?{top:b.top+b.height/2-d/2,left:b.left-c}:{top:b.top+b.height/2-d/2,left:b.left+b.width}},c.prototype.getViewportAdjustedDelta=function(a,b,c,d){var e={top:0,left:0};if(!this.$viewport)return e;var f=this.options.viewport&&this.options.viewport.padding||0,g=this.getPosition(this.$viewport);if(/right|left/.test(a)){var h=b.top-f-g.scroll,i=b.top+f-g.scroll+d;h<g.top?e.top=g.top-h:i>g.top+g.height&&(e.top=g.top+g.height-i)}else{var j=b.left-f,k=b.left+f+c;j<g.left?e.left=g.left-j:k>g.width&&(e.left=g.left+g.width-k)}return e},c.prototype.getTitle=function(){var a,b=this.$element,c=this.options;return a=b.attr("data-original-title")||("function"==typeof c.title?c.title.call(b[0]):c.title)},c.prototype.getUID=function(a){do a+=~~(1e6*Math.random());while(document.getElementById(a));return a},c.prototype.tip=function(){return this.$tip=this.$tip||a(this.options.template)},c.prototype.arrow=function(){return this.$arrow=this.$arrow||this.tip().find(".tooltip-arrow")},c.prototype.validate=function(){this.$element[0].parentNode||(this.hide(),this.$element=null,this.options=null)},c.prototype.enable=function(){this.enabled=!0},c.prototype.disable=function(){this.enabled=!1},c.prototype.toggleEnabled=function(){this.enabled=!this.enabled},c.prototype.toggle=function(b){var c=this;b&&(c=a(b.currentTarget).data("bs."+this.type),c||(c=new this.constructor(b.currentTarget,this.getDelegateOptions()),a(b.currentTarget).data("bs."+this.type,c))),c.tip().hasClass("in")?c.leave(c):c.enter(c)},c.prototype.destroy=function(){clearTimeout(this.timeout),this.hide().$element.off("."+this.type).removeData("bs."+this.type)};var d=a.fn.tooltip;a.fn.tooltip=b,a.fn.tooltip.Constructor=c,a.fn.tooltip.noConflict=function(){return a.fn.tooltip=d,this}}(jQuery),+function(a){"use strict";function b(b){return this.each(function(){var d=a(this),e=d.data("bs.popover"),f="object"==typeof b&&b;(e||"destroy"!=b)&&(e||d.data("bs.popover",e=new c(this,f)),"string"==typeof b&&e[b]())})}var c=function(a,b){this.init("popover",a,b)};if(!a.fn.tooltip)throw new Error("Popover requires tooltip.js");c.VERSION="3.2.0",c.DEFAULTS=a.extend({},a.fn.tooltip.Constructor.DEFAULTS,{placement:"right",trigger:"click",content:"",template:'<div class="popover" role="tooltip"><div class="arrow"></div><h3 class="popover-title"></h3><div class="popover-content"></div></div>'}),c.prototype=a.extend({},a.fn.tooltip.Constructor.prototype),c.prototype.constructor=c,c.prototype.getDefaults=function(){return c.DEFAULTS},c.prototype.setContent=function(){var a=this.tip(),b=this.getTitle(),c=this.getContent();a.find(".popover-title")[this.options.html?"html":"text"](b),a.find(".popover-content").empty()[this.options.html?"string"==typeof c?"html":"append":"text"](c),a.removeClass("fade top bottom left right in"),a.find(".popover-title").html()||a.find(".popover-title").hide()},c.prototype.hasContent=function(){return this.getTitle()||this.getContent()},c.prototype.getContent=function(){var a=this.$element,b=this.options;return a.attr("data-content")||("function"==typeof b.content?b.content.call(a[0]):b.content)},c.prototype.arrow=function(){return this.$arrow=this.$arrow||this.tip().find(".arrow")},c.prototype.tip=function(){return this.$tip||(this.$tip=a(this.options.template)),this.$tip};var d=a.fn.popover;a.fn.popover=b,a.fn.popover.Constructor=c,a.fn.popover.noConflict=function(){return a.fn.popover=d,this}}(jQuery),+function(a){"use strict";function b(c,d){var e=a.proxy(this.process,this);this.$body=a("body"),this.$scrollElement=a(a(c).is("body")?window:c),this.options=a.extend({},b.DEFAULTS,d),this.selector=(this.options.target||"")+" .nav li > a",this.offsets=[],this.targets=[],this.activeTarget=null,this.scrollHeight=0,this.$scrollElement.on("scroll.bs.scrollspy",e),this.refresh(),this.process()}function c(c){return this.each(function(){var d=a(this),e=d.data("bs.scrollspy"),f="object"==typeof c&&c;e||d.data("bs.scrollspy",e=new b(this,f)),"string"==typeof c&&e[c]()})}b.VERSION="3.2.0",b.DEFAULTS={offset:10},b.prototype.getScrollHeight=function(){return this.$scrollElement[0].scrollHeight||Math.max(this.$body[0].scrollHeight,document.documentElement.scrollHeight)},b.prototype.refresh=function(){var b="offset",c=0;a.isWindow(this.$scrollElement[0])||(b="position",c=this.$scrollElement.scrollTop()),this.offsets=[],this.targets=[],this.scrollHeight=this.getScrollHeight();var d=this;this.$body.find(this.selector).map(function(){var d=a(this),e=d.data("target")||d.attr("href"),f=/^#./.test(e)&&a(e);return f&&f.length&&f.is(":visible")&&[[f[b]().top+c,e]]||null}).sort(function(a,b){return a[0]-b[0]}).each(function(){d.offsets.push(this[0]),d.targets.push(this[1])})},b.prototype.process=function(){var a,b=this.$scrollElement.scrollTop()+this.options.offset,c=this.getScrollHeight(),d=this.options.offset+c-this.$scrollElement.height(),e=this.offsets,f=this.targets,g=this.activeTarget;if(this.scrollHeight!=c&&this.refresh(),b>=d)return g!=(a=f[f.length-1])&&this.activate(a);if(g&&b<=e[0])return g!=(a=f[0])&&this.activate(a);for(a=e.length;a--;)g!=f[a]&&b>=e[a]&&(!e[a+1]||b<=e[a+1])&&this.activate(f[a])},b.prototype.activate=function(b){this.activeTarget=b,a(this.selector).parentsUntil(this.options.target,".active").removeClass("active");var c=this.selector+'[data-target="'+b+'"],'+this.selector+'[href="'+b+'"]',d=a(c).parents("li").addClass("active");d.parent(".dropdown-menu").length&&(d=d.closest("li.dropdown").addClass("active")),d.trigger("activate.bs.scrollspy")};var d=a.fn.scrollspy;a.fn.scrollspy=c,a.fn.scrollspy.Constructor=b,a.fn.scrollspy.noConflict=function(){return a.fn.scrollspy=d,this},a(window).on("load.bs.scrollspy.data-api",function(){a('[data-spy="scroll"]').each(function(){var b=a(this);c.call(b,b.data())})})}(jQuery),+function(a){"use strict";function b(b){return this.each(function(){var d=a(this),e=d.data("bs.tab");e||d.data("bs.tab",e=new c(this)),"string"==typeof b&&e[b]()})}var c=function(b){this.element=a(b)};c.VERSION="3.2.0",c.prototype.show=function(){var b=this.element,c=b.closest("ul:not(.dropdown-menu)"),d=b.data("target");if(d||(d=b.attr("href"),d=d&&d.replace(/.*(?=#[^\s]*$)/,"")),!b.parent("li").hasClass("active")){var e=c.find(".active:last a")[0],f=a.Event("show.bs.tab",{relatedTarget:e});if(b.trigger(f),!f.isDefaultPrevented()){var g=a(d);this.activate(b.closest("li"),c),this.activate(g,g.parent(),function(){b.trigger({type:"shown.bs.tab",relatedTarget:e})})}}},c.prototype.activate=function(b,c,d){function e(){f.removeClass("active").find("> .dropdown-menu > .active").removeClass("active"),b.addClass("active"),g?(b[0].offsetWidth,b.addClass("in")):b.removeClass("fade"),b.parent(".dropdown-menu")&&b.closest("li.dropdown").addClass("active"),d&&d()}var f=c.find("> .active"),g=d&&a.support.transition&&f.hasClass("fade");g?f.one("bsTransitionEnd",e).emulateTransitionEnd(150):e(),f.removeClass("in")};var d=a.fn.tab;a.fn.tab=b,a.fn.tab.Constructor=c,a.fn.tab.noConflict=function(){return a.fn.tab=d,this},a(document).on("click.bs.tab.data-api",'[data-toggle="tab"], [data-toggle="pill"]',function(c){c.preventDefault(),b.call(a(this),"show")})}(jQuery),+function(a){"use strict";function b(b){return this.each(function(){var d=a(this),e=d.data("bs.affix"),f="object"==typeof b&&b;e||d.data("bs.affix",e=new c(this,f)),"string"==typeof b&&e[b]()})}var c=function(b,d){this.options=a.extend({},c.DEFAULTS,d),this.$target=a(this.options.target).on("scroll.bs.affix.data-api",a.proxy(this.checkPosition,this)).on("click.bs.affix.data-api",a.proxy(this.checkPositionWithEventLoop,this)),this.$element=a(b),this.affixed=this.unpin=this.pinnedOffset=null,this.checkPosition()};c.VERSION="3.2.0",c.RESET="affix affix-top affix-bottom",c.DEFAULTS={offset:0,target:window},c.prototype.getPinnedOffset=function(){if(this.pinnedOffset)return this.pinnedOffset;this.$element.removeClass(c.RESET).addClass("affix");var a=this.$target.scrollTop(),b=this.$element.offset();return this.pinnedOffset=b.top-a},c.prototype.checkPositionWithEventLoop=function(){setTimeout(a.proxy(this.checkPosition,this),1)},c.prototype.checkPosition=function(){if(this.$element.is(":visible")){var b=a(document).height(),d=this.$target.scrollTop(),e=this.$element.offset(),f=this.options.offset,g=f.top,h=f.bottom;"object"!=typeof f&&(h=g=f),"function"==typeof g&&(g=f.top(this.$element)),"function"==typeof h&&(h=f.bottom(this.$element));var i=null!=this.unpin&&d+this.unpin<=e.top?!1:null!=h&&e.top+this.$element.height()>=b-h?"bottom":null!=g&&g>=d?"top":!1;if(this.affixed!==i){null!=this.unpin&&this.$element.css("top","");var j="affix"+(i?"-"+i:""),k=a.Event(j+".bs.affix");this.$element.trigger(k),k.isDefaultPrevented()||(this.affixed=i,this.unpin="bottom"==i?this.getPinnedOffset():null,this.$element.removeClass(c.RESET).addClass(j).trigger(a.Event(j.replace("affix","affixed"))),"bottom"==i&&this.$element.offset({top:b-this.$element.height()-h}))}}};var d=a.fn.affix;a.fn.affix=b,a.fn.affix.Constructor=c,a.fn.affix.noConflict=function(){return a.fn.affix=d,this},a(window).on("load",function(){a('[data-spy="affix"]').each(function(){var c=a(this),d=c.data();d.offset=d.offset||{},d.offsetBottom&&(d.offset.bottom=d.offsetBottom),d.offsetTop&&(d.offset.top=d.offsetTop),b.call(c,d)})})}(jQuery);
```
