## Supplementary File 2 for "Single individual structural variant detection uncovers widespread hemizygosity in molluscs": jquery-migrate-1.js

```
/*! jQuery Migrate v1.2.1 | (c) 2005, 2013 jQuery Foundation, Inc. and other contributors | jquery.org/license */
jQuery.migrateMute===void 0&&(jQuery.migrateMute=!0),function(e,t,n){function r(n){var r=t.console;i[n]||(i[n]=!0,e.migrateWarnings.push(n),r&&r.warn&&!e.migrateMute&&(r.warn("JQMIGRATE: "+n),e.migrateTrace&&r.trace&&r.trace()))}function a(t,a,i,o){if(Object.defineProperty)try{return Object.defineProperty(t,a,{configurable:!0,enumerable:!0,get:function(){return r(o),i},set:function(e){r(o),i=e}}),n}catch(s){}e._definePropertyBroken=!0,t[a]=i}var i={};e.migrateWarnings=[],!e.migrateMute&&t.console&&t.console.log&&t.console.log("JQMIGRATE: Logging is active"),e.migrateTrace===n&&(e.migrateTrace=!0),e.migrateReset=function(){i={},e.migrateWarnings.length=0},"BackCompat"===document.compatMode&&r("jQuery is not compatible with Quirks Mode");var o=e("<input/>",{size:1}).attr("size")&&e.attrFn,s=e.attr,u=e.attrHooks.value&&e.attrHooks.value.get||function(){return null},c=e.attrHooks.value&&e.attrHooks.value.set||function(){return n},l=/^(?:input|button)$/i,d=/^[238]$/,p=/^(?:autofocus|autoplay|async|checked|controls|defer|disabled|hidden|loop|multiple|open|readonly|required|scoped|selected)$/i,f=/^(?:checked|selected)$/i;a(e,"attrFn",o||{},"jQuery.attrFn is deprecated"),e.attr=function(t,a,i,u){var c=a.toLowerCase(),g=t&&t.nodeType;return u&&(4>s.length&&r("jQuery.fn.attr( props, pass ) is deprecated"),t&&!d.test(g)&&(o?a in o:e.isFunction(e.fn[a])))?e(t)[a](i):("type"===a&&i!==n&&l.test(t.nodeName)&&t.parentNode&&r("Can't change the 'type' of an input or button in IE 6/7/8"),!e.attrHooks[c]&&p.test(c)&&(e.attrHooks[c]={get:function(t,r){var a,i=e.prop(t,r);return i===!0||"boolean"!=typeof i&&(a=t.getAttributeNode(r))&&a.nodeValue!==!1?r.toLowerCase():n},set:function(t,n,r){var a;return n===!1?e.removeAttr(t,r):(a=e.propFix[r]||r,a in t&&(t[a]=!0),t.setAttribute(r,r.toLowerCase())),r}},f.test(c)&&r("jQuery.fn.attr('"+c+"') may use property instead of attribute")),s.call(e,t,a,i))},e.attrHooks.value={get:function(e,t){var n=(e.nodeName||"").toLowerCase();return"button"===n?u.apply(this,arguments):("input"!==n&&"option"!==n&&r("jQuery.fn.attr('value') no longer gets properties"),t in e?e.value:null)},set:function(e,t){var a=(e.nodeName||"").toLowerCase();return"button"===a?c.apply(this,arguments):("input"!==a&&"option"!==a&&r("jQuery.fn.attr('value', val) no longer sets properties"),e.value=t,n)}};var g,h,v=e.fn.init,m=e.parseJSON,y=/^([^<]*)(<[\w\W]+>)([^>]*)$/;e.fn.init=function(t,n,a){var i;return t&&"string"==typeof t&&!e.isPlainObject(n)&&(i=y.exec(e.trim(t)))&&i[0]&&("<"!==t.charAt(0)&&r("$(html) HTML strings must start with '<' character"),i[3]&&r("$(html) HTML text after last tag is ignored"),"#"===i[0].charAt(0)&&(r("HTML string cannot start with a '#' character"),e.error("JQMIGRATE: Invalid selector string (XSS)")),n&&n.context&&(n=n.context),e.parseHTML)?v.call(this,e.parseHTML(i[2],n,!0),n,a):v.apply(this,arguments)},e.fn.init.prototype=e.fn,e.parseJSON=function(e){return e||null===e?m.apply(this,arguments):(r("jQuery.parseJSON requires a valid JSON string"),null)},e.uaMatch=function(e){e=e.toLowerCase();var t=/(chrome)[ \/]([\w.]+)/.exec(e)||/(webkit)[ \/]([\w.]+)/.exec(e)||/(opera)(?:.*version|)[ \/]([\w.]+)/.exec(e)||/(msie) ([\w.]+)/.exec(e)||0>e.indexOf("compatible")&&/(mozilla)(?:.*? rv:([\w.]+)|)/.exec(e)||[];return{browser:t[1]||"",version:t[2]||"0"}},e.browser||(g=e.uaMatch(navigator.userAgent),h={},g.browser&&(h[g.browser]=!0,h.version=g.version),h.chrome?h.webkit=!0:h.webkit&&(h.safari=!0),e.browser=h),a(e,"browser",e.browser,"jQuery.browser is deprecated"),e.sub=function(){function t(e,n){return new t.fn.init(e,n)}e.extend(!0,t,this),t.superclass=this,t.fn=t.prototype=this(),t.fn.constructor=t,t.sub=this.sub,t.fn.init=function(r,a){return a&&a instanceof e&&!(a instanceof t)&&(a=t(a)),e.fn.init.call(this,r,a,n)},t.fn.init.prototype=t.fn;var n=t(document);return r("jQuery.sub() is deprecated"),t},e.ajaxSetup({converters:{"text json":e.parseJSON}});var b=e.fn.data;e.fn.data=function(t){var a,i,o=this[0];return!o||"events"!==t||1!==arguments.length||(a=e.data(o,t),i=e._data(o,t),a!==n&&a!==i||i===n)?b.apply(this,arguments):(r("Use of jQuery.fn.data('events') is deprecated"),i)};var j=/\/(java|ecma)script/i,w=e.fn.andSelf||e.fn.addBack;e.fn.andSelf=function(){return r("jQuery.fn.andSelf() replaced by jQuery.fn.addBack()"),w.apply(this,arguments)},e.clean||(e.clean=function(t,a,i,o){a=a||document,a=!a.nodeType&&a[0]||a,a=a.ownerDocument||a,r("jQuery.clean() is deprecated");var s,u,c,l,d=[];if(e.merge(d,e.buildFragment(t,a).childNodes),i)for(c=function(e){return!e.type||j.test(e.type)?o?o.push(e.parentNode?e.parentNode.removeChild(e):e):i.appendChild(e):n},s=0;null!=(u=d[s]);s++)e.nodeName(u,"script")&&c(u)||(i.appendChild(u),u.getElementsByTagName!==n&&(l=e.grep(e.merge([],u.getElementsByTagName("script")),c),d.splice.apply(d,[s+1,0].concat(l)),s+=l.length));return d});var Q=e.event.add,x=e.event.remove,k=e.event.trigger,N=e.fn.toggle,T=e.fn.live,M=e.fn.die,S="ajaxStart|ajaxStop|ajaxSend|ajaxComplete|ajaxError|ajaxSuccess",C=RegExp("\\b(?:"+S+")\\b"),H=/(?:^|\s)hover(\.\S+|)\b/,A=function(t){return"string"!=typeof t||e.event.special.hover?t:(H.test(t)&&r("'hover' pseudo-event is deprecated, use 'mouseenter mouseleave'"),t&&t.replace(H,"mouseenter$1 mouseleave$1"))};e.event.props&&"attrChange"!==e.event.props[0]&&e.event.props.unshift("attrChange","attrName","relatedNode","srcElement"),e.event.dispatch&&a(e.event,"handle",e.event.dispatch,"jQuery.event.handle is undocumented and deprecated"),e.event.add=function(e,t,n,a,i){e!==document&&C.test(t)&&r("AJAX events should be attached to document: "+t),Q.call(this,e,A(t||""),n,a,i)},e.event.remove=function(e,t,n,r,a){x.call(this,e,A(t)||"",n,r,a)},e.fn.error=function(){var e=Array.prototype.slice.call(arguments,0);return r("jQuery.fn.error() is deprecated"),e.splice(0,0,"error"),arguments.length?this.bind.apply(this,e):(this.triggerHandler.apply(this,e),this)},e.fn.toggle=function(t,n){if(!e.isFunction(t)||!e.isFunction(n))return N.apply(this,arguments);r("jQuery.fn.toggle(handler, handler...) is deprecated");var a=arguments,i=t.guid||e.guid++,o=0,s=function(n){var r=(e._data(this,"lastToggle"+t.guid)||0)%o;return e._data(this,"lastToggle"+t.guid,r+1),n.preventDefault(),a[r].apply(this,arguments)||!1};for(s.guid=i;a.length>o;)a[o++].guid=i;return this.click(s)},e.fn.live=function(t,n,a){return r("jQuery.fn.live() is deprecated"),T?T.apply(this,arguments):(e(this.context).on(t,this.selector,n,a),this)},e.fn.die=function(t,n){return r("jQuery.fn.die() is deprecated"),M?M.apply(this,arguments):(e(this.context).off(t,this.selector||"**",n),this)},e.event.trigger=function(e,t,n,a){return n||C.test(e)||r("Global events are undocumented and deprecated"),k.call(this,e,t,n||document,a)},e.each(S.split("|"),function(t,n){e.event.special[n]={setup:function(){var t=this;return t!==document&&(e.event.add(document,n+"."+e.guid,function(){e.event.trigger(n,null,t,!0)}),e._data(this,n,e.guid++)),!1},teardown:function(){return this!==document&&e.event.remove(document,n+"."+e._data(this,n)),!1}}})}(jQuery,window);
```
