## Supplementary File 2 for "Single individual structural variant detection uncovers widespread hemizygosity in molluscs": jquery.js

```
RegExp.escape=function(s){return s.replace(/[-\/\\^$*+?.()|[\]{}]/g,'\\$&');};(function($){'use strict'
$.csv={defaults:{separator:',',delimiter:'"',headers:true},hooks:{castToScalar:function(value,state){var hasDot=/\./;if(isNaN(value)){return value;}else{if(hasDot.test(value)){return parseFloat(value);}else{var integer=parseInt(value);if(isNaN(integer)){return null;}else{return integer;}}}}},parsers:{parse:function(csv,options){var separator=options.separator;var delimiter=options.delimiter;if(!options.state.rowNum){options.state.rowNum=1;}
if(!options.state.colNum){options.state.colNum=1;}
var data=[];var entry=[];var state=0;var value=''
var exit=false;function endOfEntry(){state=0;value='';if(options.start&&options.state.rowNum<options.start){entry=[];options.state.rowNum++;options.state.colNum=1;return;}
if(options.onParseEntry===undefined){data.push(entry);}else{var hookVal=options.onParseEntry(entry,options.state);if(hookVal!==false){data.push(hookVal);}}
entry=[];if(options.end&&options.state.rowNum>=options.end){exit=true;}
options.state.rowNum++;options.state.colNum=1;}
function endOfValue(){if(options.onParseValue===undefined){entry.push(value);}else{var hook=options.onParseValue(value,options.state);if(hook!==false){entry.push(hook);}}
value='';state=0;options.state.colNum++;}
var escSeparator=RegExp.escape(separator);var escDelimiter=RegExp.escape(delimiter);var match=/(D|S|\n|\r|[^DS\r\n]+)/;var matchSrc=match.source;matchSrc=matchSrc.replace(/S/g,escSeparator);matchSrc=matchSrc.replace(/D/g,escDelimiter);match=RegExp(matchSrc,'gm');csv.replace(match,function(m0){if(exit){return;}
switch(state){case 0:if(m0===separator){value+='';endOfValue();break;}
if(m0===delimiter){state=1;break;}
if(m0==='\n'){endOfValue();endOfEntry();break;}
if(/^\r$/.test(m0)){break;}
value+=m0;state=3;break;case 1:if(m0===delimiter){state=2;break;}
value+=m0;state=1;break;case 2:if(m0===delimiter){value+=m0;state=1;break;}
if(m0===separator){endOfValue();break;}
if(m0==='\n'){endOfValue();endOfEntry();break;}
if(/^\r$/.test(m0)){break;}
throw new Error('CSVDataError: Illegal State [Row:'+options.state.rowNum+'][Col:'+options.state.colNum+']');case 3:if(m0===separator){endOfValue();break;}
if(m0==='\n'){endOfValue();endOfEntry();break;}
if(/^\r$/.test(m0)){break;}
if(m0===delimiter){throw new Error('CSVDataError: Illegal Quote [Row:'+options.state.rowNum+'][Col:'+options.state.colNum+']');}
throw new Error('CSVDataError: Illegal Data [Row:'+options.state.rowNum+'][Col:'+options.state.colNum+']');default:throw new Error('CSVDataError: Unknown State [Row:'+options.state.rowNum+'][Col:'+options.state.colNum+']');}});if(entry.length!==0){endOfValue();endOfEntry();}
return data;},splitLines:function(csv,options){var separator=options.separator;var delimiter=options.delimiter;if(!options.state.rowNum){options.state.rowNum=1;}
var entries=[];var state=0;var entry='';var exit=false;function endOfLine(){state=0;if(options.start&&options.state.rowNum<options.start){entry='';options.state.rowNum++;return;}
if(options.onParseEntry===undefined){entries.push(entry);}else{var hookVal=options.onParseEntry(entry,options.state);if(hookVal!==false){entries.push(hookVal);}}
entry='';if(options.end&&options.state.rowNum>=options.end){exit=true;}
options.state.rowNum++;}
var escSeparator=RegExp.escape(separator);var escDelimiter=RegExp.escape(delimiter);var match=/(D|S|\n|\r|[^DS\r\n]+)/;var matchSrc=match.source;matchSrc=matchSrc.replace(/S/g,escSeparator);matchSrc=matchSrc.replace(/D/g,escDelimiter);match=RegExp(matchSrc,'gm');csv.replace(match,function(m0){if(exit){return;}
switch(state){case 0:if(m0===separator){entry+=m0;state=0;break;}
if(m0===delimiter){entry+=m0;state=1;break;}
if(m0==='\n'){endOfLine();break;}
if(/^\r$/.test(m0)){break;}
entry+=m0;state=3;break;case 1:if(m0===delimiter){entry+=m0;state=2;break;}
entry+=m0;state=1;break;case 2:var prevChar=entry.substr(entry.length-1);if(m0===delimiter&&prevChar===delimiter){entry+=m0;state=1;break;}
if(m0===separator){entry+=m0;state=0;break;}
if(m0==='\n'){endOfLine();break;}
if(m0==='\r'){break;}
throw new Error('CSVDataError: Illegal state [Row:'+options.state.rowNum+']');case 3:if(m0===separator){entry+=m0;state=0;break;}
if(m0==='\n'){endOfLine();break;}
if(m0==='\r'){break;}
if(m0===delimiter){throw new Error('CSVDataError: Illegal quote [Row:'+options.state.rowNum+']');}
throw new Error('CSVDataError: Illegal state [Row:'+options.state.rowNum+']');default:throw new Error('CSVDataError: Unknown state [Row:'+options.state.rowNum+']');}});if(entry!==''){endOfLine();}
return entries;},parseEntry:function(csv,options){var separator=options.separator;var delimiter=options.delimiter;if(!options.state.rowNum){options.state.rowNum=1;}
if(!options.state.colNum){options.state.colNum=1;}
var entry=[];var state=0;var value='';function endOfValue(){if(options.onParseValue===undefined){entry.push(value);}else{var hook=options.onParseValue(value,options.state);if(hook!==false){entry.push(hook);}}
value='';state=0;options.state.colNum++;}
if(!options.match){var escSeparator=RegExp.escape(separator);var escDelimiter=RegExp.escape(delimiter);var match=/(D|S|\n|\r|[^DS\r\n]+)/;var matchSrc=match.source;matchSrc=matchSrc.replace(/S/g,escSeparator);matchSrc=matchSrc.replace(/D/g,escDelimiter);options.match=RegExp(matchSrc,'gm');}
csv.replace(options.match,function(m0){switch(state){case 0:if(m0===separator){value+='';endOfValue();break;}
if(m0===delimiter){state=1;break;}
if(m0==='\n'||m0==='\r'){break;}
value+=m0;state=3;break;case 1:if(m0===delimiter){state=2;break;}
value+=m0;state=1;break;case 2:if(m0===delimiter){value+=m0;state=1;break;}
if(m0===separator){endOfValue();break;}
if(m0==='\n'||m0==='\r'){break;}
throw new Error('CSVDataError: Illegal State [Row:'+options.state.rowNum+'][Col:'+options.state.colNum+']');case 3:if(m0===separator){endOfValue();break;}
if(m0==='\n'||m0==='\r'){break;}
if(m0===delimiter){throw new Error('CSVDataError: Illegal Quote [Row:'+options.state.rowNum+'][Col:'+options.state.colNum+']');}
throw new Error('CSVDataError: Illegal Data [Row:'+options.state.rowNum+'][Col:'+options.state.colNum+']');default:throw new Error('CSVDataError: Unknown State [Row:'+options.state.rowNum+'][Col:'+options.state.colNum+']');}});endOfValue();return entry;}},toArray:function(csv,options,callback){var options=(options!==undefined?options:{});var config={};config.callback=((callback!==undefined&&typeof(callback)==='function')?callback:false);config.separator='separator'in options?options.separator:$.csv.defaults.separator;config.delimiter='delimiter'in options?options.delimiter:$.csv.defaults.delimiter;var state=(options.state!==undefined?options.state:{});var options={delimiter:config.delimiter,separator:config.separator,onParseEntry:options.onParseEntry,onParseValue:options.onParseValue,state:state}
var entry=$.csv.parsers.parseEntry(csv,options);if(!config.callback){return entry;}else{config.callback('',entry);}},toArrays:function(csv,options,callback){var options=(options!==undefined?options:{});var config={};config.callback=((callback!==undefined&&typeof(callback)==='function')?callback:false);config.separator='separator'in options?options.separator:$.csv.defaults.separator;config.delimiter='delimiter'in options?options.delimiter:$.csv.defaults.delimiter;var data=[];var options={delimiter:config.delimiter,separator:config.separator,onParseEntry:options.onParseEntry,onParseValue:options.onParseValue,start:options.start,end:options.end,state:{rowNum:1,colNum:1}};data=$.csv.parsers.parse(csv,options);if(!config.callback){return data;}else{config.callback('',data);}},toObjects:function(csv,options,callback){var options=(options!==undefined?options:{});var config={};config.callback=((callback!==undefined&&typeof(callback)==='function')?callback:false);config.separator='separator'in options?options.separator:$.csv.defaults.separator;config.delimiter='delimiter'in options?options.delimiter:$.csv.defaults.delimiter;config.headers='headers'in options?options.headers:$.csv.defaults.headers;options.start='start'in options?options.start:1;if(config.headers){options.start++;}
if(options.end&&config.headers){options.end++;}
var lines=[];var data=[];var options={delimiter:config.delimiter,separator:config.separator,onParseEntry:options.onParseEntry,onParseValue:options.onParseValue,start:options.start,end:options.end,state:{rowNum:1,colNum:1},match:false};var headerOptions={delimiter:config.delimiter,separator:config.separator,start:1,end:1,state:{rowNum:1,colNum:1}}
var headerLine=$.csv.parsers.splitLines(csv,headerOptions);var headers=$.csv.toArray(headerLine[0],options);var lines=$.csv.parsers.splitLines(csv,options);options.state.colNum=1;if(headers){options.state.rowNum=2;}else{options.state.rowNum=1;}
for(var i=0,len=lines.length;i<len;i++){var entry=$.csv.toArray(lines[i],options);var object={};for(var j in headers){object[headers[j]]=entry[j];}
data.push(object);options.state.rowNum++;}
if(!config.callback){return data;}else{config.callback('',data);}},fromArrays:function(arrays,options,callback){var options=(options!==undefined?options:{});var config={};config.callback=((callback!==undefined&&typeof(callback)==='function')?callback:false);config.separator='separator'in options?options.separator:$.csv.defaults.separator;config.delimiter='delimiter'in options?options.delimiter:$.csv.defaults.delimiter;config.escaper='escaper'in options?options.escaper:$.csv.defaults.escaper;config.experimental='experimental'in options?options.experimental:false;if(!config.experimental){throw new Error('not implemented');}
var output=[];for(i in arrays){output.push(arrays[i]);}
if(!config.callback){return output;}else{config.callback('',output);}},fromObjects2CSV:function(objects,options,callback){var options=(options!==undefined?options:{});var config={};config.callback=((callback!==undefined&&typeof(callback)==='function')?callback:false);config.separator='separator'in options?options.separator:$.csv.defaults.separator;config.delimiter='delimiter'in options?options.delimiter:$.csv.defaults.delimiter;config.experimental='experimental'in options?options.experimental:false;if(!config.experimental){throw new Error('not implemented');}
var output=[];for(i in objects){output.push(arrays[i]);}
if(!config.callback){return output;}else{config.callback('',output);}}};$.csvEntry2Array=$.csv.toArray;$.csv2Array=$.csv.toArrays;$.csv2Dictionary=$.csv.toObjects;})(jQuery);
```
