## Supplementary File 2 for "Single individual structural variant detection uncovers widespread hemizygosity in molluscs": jquery_003.js

```
(function(e){var t,n,r,i,a,o,s,l,c,u,d,p,h,m=0,g={},f=[],b=0,v={},y=[],_=null,k=new Image,E=/\.(jpg|gif|png|bmp|jpeg)(.*)?$/i,w=/[^\.]\.(swf)\s*$/i,C=1,x=0,T="",S=!1,A=e.extend(e("<div/>")[0],{prop:0}),R=e.browser.msie&&7>e.browser.version&&!window.XMLHttpRequest,I=function(){n.hide(),k.onerror=k.onload=null,_&&_.abort(),t.empty()},D=function(){!1===g.onError(f,m,g)?(n.hide(),S=!1):(g.titleShow=!1,g.width="auto",g.height="auto",t.html('<p id="fancybox-error">The requested content cannot be loaded.<br />Please try again later.</p>'),L())},O=function(){var r,i,a,s,l,c,u=f[m];if(I(),g=e.extend({},e.fn.fancybox.defaults,e(u).data("fancybox")===void 0?g:e(u).data("fancybox")),c=g.onStart(f,m,g),c===!1)S=!1;else if("object"==typeof c&&(g=e.extend(g,c)),a=g.title||(u.nodeName?e(u).attr("title"):u.title)||"",u.nodeName&&!g.orig&&(g.orig=e(u).children("img:first").length?e(u).children("img:first"):e(u)),""===a&&g.orig&&g.titleFromAlt&&(a=g.orig.attr("alt")),r=g.href||(u.nodeName?e(u).attr("href"):u.href)||null,(/^(?:javascript)/i.test(r)||"#"==r)&&(r=null),g.type?(i=g.type,r||(r=g.content)):g.content?i="html":r&&(i=r.match(E)?"image":r.match(w)?"swf":e(u).hasClass("iframe")?"iframe":0===r.indexOf("#")?"inline":"ajax"),i)switch("inline"==i&&(u=r.substr(r.indexOf("#")),i=e(u).length>0?"inline":"ajax"),g.type=i,g.href=r,g.title=a,g.autoDimensions&&("html"==g.type||"inline"==g.type||"ajax"==g.type?(g.width="auto",g.height="auto"):g.autoDimensions=!1),g.modal&&(g.overlayShow=!0,g.hideOnOverlayClick=!1,g.hideOnContentClick=!1,g.enableEscapeButton=!1,g.showCloseButton=!1),g.padding=parseInt(g.padding,10),g.margin=parseInt(g.margin,10),t.css("padding",g.padding+g.margin),e(".fancybox-inline-tmp").unbind("fancybox-cancel").bind("fancybox-change",function(){e(this).replaceWith(o.children())}),i){case"html":t.html(g.content),L();break;case"inline":if(e(u).parent().is("#fancybox-content")===!0){S=!1;break}e('<div class="fancybox-inline-tmp" />').hide().insertBefore(e(u)).bind("fancybox-cleanup",function(){e(this).replaceWith(o.children())}).bind("fancybox-cancel",function(){e(this).replaceWith(t.children())}),e(u).appendTo(t),L();break;case"image":S=!1,e.fancybox.showActivity(),k=new Image,k.onerror=function(){D()},k.onload=function(){S=!0,k.onerror=k.onload=null,g.width=k.width,g.height=k.height,e("<img />").attr({id:"fancybox-img",src:k.src,alt:g.title}).appendTo(t),N()},k.src=r;break;case"swf":g.scrolling="no",s='<object classid="clsid:D27CDB6E-AE6D-11cf-96B8-444553540000" width="'+g.width+'" height="'+g.height+'"><param name="movie" value="'+r+'"></param>',l="",e.each(g.swf,function(e,t){s+='<param name="'+e+'" value="'+t+'"></param>',l+=" "+e+'="'+t+'"'}),s+='<embed src="'+r+'" type="application/x-shockwave-flash" width="'+g.width+'" height="'+g.height+'"'+l+"></embed></object>",t.html(s),L();break;case"ajax":S=!1,e.fancybox.showActivity(),g.ajax.win=g.ajax.success,_=e.ajax(e.extend({},g.ajax,{url:r,data:g.ajax.data||{},error:function(e){e.status>0&&D()},success:function(e,i,a){if(200==("object"==typeof a?a:_).status){if("function"==typeof g.ajax.win){if(c=g.ajax.win(r,e,i,a),c===!1)return n.hide(),void 0;("string"==typeof c||"object"==typeof c)&&(e=c)}t.html(e),L()}}}));break;case"iframe":N()}else D()},L=function(){var n=g.width,r=g.height;n=(""+n).indexOf("%")>-1?parseInt((e(window).width()-2*g.margin)*parseFloat(n)/100,10)+"px":"auto"==n?"auto":n+"px",r=(""+r).indexOf("%")>-1?parseInt((e(window).height()-2*g.margin)*parseFloat(r)/100,10)+"px":"auto"==r?"auto":r+"px",t.wrapInner('<div style="width:'+n+";height:"+r+";overflow: "+("auto"==g.scrolling?"auto":"yes"==g.scrolling?"scroll":"hidden")+';position:relative;"></div>'),g.width=t.width(),g.height=t.height(),N()},N=function(){var d,_;if(n.hide(),i.is(":visible")&&!1===v.onCleanup(y,b,v))e.event.trigger("fancybox-cancel"),S=!1;else{if(S=!0,e(o.add(r)).unbind(),e(window).unbind("resize.fb scroll.fb"),e(document).unbind("keydown.fb"),i.is(":visible")&&"outside"!==v.titlePosition&&i.css("height",i.height()),y=f,b=m,v=g,v.overlayShow?(r.css({"background-color":v.overlayColor,opacity:v.overlayOpacity,cursor:v.hideOnOverlayClick?"pointer":"auto",height:e(document).height()}),r.is(":visible")||(R&&e("select:not(#fancybox-tmp select)").filter(function(){return"hidden"!==this.style.visibility}).css({visibility:"hidden"}).one("fancybox-cleanup",function(){this.style.visibility="inherit"}),r.show())):r.hide(),h=j(),T=v.title||"",x=0,l.empty().removeAttr("style").removeClass(),v.titleShow!==!1&&(d=e.isFunction(v.titleFormat)?v.titleFormat(T,y,b,v):T&&T.length?"float"==v.titlePosition?'<table id="fancybox-title-float-wrap" cellpadding="0" cellspacing="0"><tr><td id="fancybox-title-float-left"></td><td id="fancybox-title-float-main">'+T+'</td><td id="fancybox-title-float-right"></td></tr></table>':'<div id="fancybox-title-'+v.titlePosition+'">'+T+"</div>":!1,T=d,T&&""!==T))switch(l.addClass("fancybox-title-"+v.titlePosition).html(T).appendTo("body").show(),v.titlePosition){case"inside":l.css({width:h.width-2*v.padding,marginLeft:v.padding,marginRight:v.padding}),x=l.outerHeight(!0),l.appendTo(a),h.height+=x;break;case"over":l.css({marginLeft:v.padding,width:h.width-2*v.padding,bottom:v.padding}).appendTo(a);break;case"float":l.css("left",-1*parseInt((l.width()-h.width-40)/2,10)).appendTo(i);break;default:l.css({width:h.width-2*v.padding,paddingLeft:v.padding,paddingRight:v.padding}).appendTo(i)}l.hide(),i.is(":visible")?(e(s.add(c).add(u)).hide(),d=i.position(),p={top:d.top,left:d.left,width:i.width(),height:i.height()},_=p.width==h.width&&p.height==h.height,o.fadeTo(v.changeFade,.3,function(){var n=function(){o.html(t.contents()).fadeTo(v.changeFade,1,P)};e.event.trigger("fancybox-change"),o.empty().removeAttr("filter").css({"border-width":v.padding,width:h.width-2*v.padding,height:g.autoDimensions?"auto":h.height-x-2*v.padding}),_?n():(A.prop=0,e(A).animate({prop:1},{duration:v.changeSpeed,easing:v.easingChange,step:B,complete:n}))})):(i.removeAttr("style"),o.css("border-width",v.padding),"elastic"==v.transitionIn?(p=z(),o.html(t.contents()),i.show(),v.opacity&&(h.opacity=0),A.prop=0,e(A).animate({prop:1},{duration:v.speedIn,easing:v.easingIn,step:B,complete:P})):("inside"==v.titlePosition&&x>0&&l.show(),o.css({width:h.width-2*v.padding,height:g.autoDimensions?"auto":h.height-x-2*v.padding}).html(t.contents()),i.css(h).fadeIn("none"==v.transitionIn?0:v.speedIn,P)))}},F=function(){(v.enableEscapeButton||v.enableKeyboardNav)&&e(document).bind("keydown.fb",function(t){27==t.keyCode&&v.enableEscapeButton?(t.preventDefault(),e.fancybox.close()):37!=t.keyCode&&39!=t.keyCode||!v.enableKeyboardNav||"INPUT"===t.target.tagName||"TEXTAREA"===t.target.tagName||"SELECT"===t.target.tagName||(t.preventDefault(),e.fancybox[37==t.keyCode?"prev":"next"]())}),v.showNavArrows?((v.cyclic&&y.length>1||0!==b)&&c.show(),(v.cyclic&&y.length>1||b!=y.length-1)&&u.show()):(c.hide(),u.hide())},P=function(){e.support.opacity||(o.get(0).style.removeAttribute("filter"),i.get(0).style.removeAttribute("filter")),g.autoDimensions&&o.css("height","auto"),i.css("height","auto"),T&&T.length&&l.show(),v.showCloseButton&&s.show(),F(),v.hideOnContentClick&&o.bind("click",e.fancybox.close),v.hideOnOverlayClick&&r.bind("click",e.fancybox.close),e(window).bind("resize.fb",e.fancybox.resize),v.centerOnScroll&&e(window).bind("scroll.fb",e.fancybox.center),"iframe"==v.type&&e('<iframe id="fancybox-frame" name="fancybox-frame'+(new Date).getTime()+'" frameborder="0" hspace="0" '+(e.browser.msie?'allowtransparency="true""':"")+' scrolling="'+g.scrolling+'" src="'+v.href+'"></iframe>').appendTo(o),i.show(),S=!1,e.fancybox.center(),v.onComplete(y,b,v);var t,n;y.length-1>b&&(t=y[b+1].href,t!==void 0&&t.match(E)&&(n=new Image,n.src=t)),b>0&&(t=y[b-1].href,t!==void 0&&t.match(E)&&(n=new Image,n.src=t))},B=function(e){var t={width:parseInt(p.width+(h.width-p.width)*e,10),height:parseInt(p.height+(h.height-p.height)*e,10),top:parseInt(p.top+(h.top-p.top)*e,10),left:parseInt(p.left+(h.left-p.left)*e,10)};h.opacity!==void 0&&(t.opacity=.5>e?.5:e),i.css(t),o.css({width:t.width-2*v.padding,height:t.height-x*e-2*v.padding})},M=function(){return[e(window).width()-2*v.margin,e(window).height()-2*v.margin,e(document).scrollLeft()+v.margin,e(document).scrollTop()+v.margin]},j=function(){var e=M(),t={},n=v.autoScale,r=2*v.padding;return t.width=(""+v.width).indexOf("%")>-1?parseInt(e[0]*parseFloat(v.width)/100,10):v.width+r,t.height=(""+v.height).indexOf("%")>-1?parseInt(e[1]*parseFloat(v.height)/100,10):v.height+r,n&&(t.width>e[0]||t.height>e[1])&&("image"==g.type||"swf"==g.type?(n=v.width/v.height,t.width>e[0]&&(t.width=e[0],t.height=parseInt((t.width-r)/n+r,10)),t.height>e[1]&&(t.height=e[1],t.width=parseInt((t.height-r)*n+r,10))):(t.width=Math.min(t.width,e[0]),t.height=Math.min(t.height,e[1]))),t.top=parseInt(Math.max(e[3]-20,e[3]+.5*(e[1]-t.height-40)),10),t.left=parseInt(Math.max(e[2]-20,e[2]+.5*(e[0]-t.width-40)),10),t},z=function(){var t=g.orig?e(g.orig):!1,n={};return t&&t.length?(n=t.offset(),n.top+=parseInt(t.css("paddingTop"),10)||0,n.left+=parseInt(t.css("paddingLeft"),10)||0,n.top+=parseInt(t.css("border-top-width"),10)||0,n.left+=parseInt(t.css("border-left-width"),10)||0,n.width=t.width(),n.height=t.height(),n={width:n.width+2*v.padding,height:n.height+2*v.padding,top:n.top-v.padding-20,left:n.left-v.padding-20}):(t=M(),n={width:2*v.padding,height:2*v.padding,top:parseInt(t[3]+.5*t[1],10),left:parseInt(t[2]+.5*t[0],10)}),n},K=function(){n.is(":visible")?(e("div",n).css("top",-40*C+"px"),C=(C+1)%12):clearInterval(d)};e.fn.fancybox=function(t){return e(this).length?(e(this).data("fancybox",e.extend({},t,e.metadata?e(this).metadata():{})).unbind("click.fb").bind("click.fb",function(t){t.preventDefault(),S||(S=!0,e(this).blur(),f=[],m=0,t=e(this).attr("rel")||"",t&&""!=t&&"nofollow"!==t?(f=e("a[rel="+t+"], area[rel="+t+"]"),m=f.index(this)):f.push(this),O())}),this):this},e.fancybox=function(t,n){var r;if(!S){if(S=!0,r=n!==void 0?n:{},f=[],m=parseInt(r.index,10)||0,e.isArray(t)){for(var i=0,a=t.length;a>i;i++)"object"==typeof t[i]?e(t[i]).data("fancybox",e.extend({},r,t[i])):t[i]=e({}).data("fancybox",e.extend({content:t[i]},r));f=jQuery.merge(f,t)}else"object"==typeof t?e(t).data("fancybox",e.extend({},r,t)):t=e({}).data("fancybox",e.extend({content:t},r)),f.push(t);(m>f.length||0>m)&&(m=0),O()}},e.fancybox.showActivity=function(){clearInterval(d),n.show(),d=setInterval(K,66)},e.fancybox.hideActivity=function(){n.hide()},e.fancybox.next=function(){return e.fancybox.pos(b+1)},e.fancybox.prev=function(){return e.fancybox.pos(b-1)},e.fancybox.pos=function(e){S||(e=parseInt(e),f=y,e>-1&&y.length>e?(m=e,O()):v.cyclic&&y.length>1&&(m=e>=y.length?0:y.length-1,O()))},e.fancybox.cancel=function(){S||(S=!0,e.event.trigger("fancybox-cancel"),I(),g.onCancel(f,m,g),S=!1)},e.fancybox.close=function(){function t(){r.fadeOut("fast"),l.empty().hide(),i.hide(),e.event.trigger("fancybox-cleanup"),o.empty(),v.onClosed(y,b,v),y=g=[],b=m=0,v=g={},S=!1}if(!S&&!i.is(":hidden"))if(S=!0,v&&!1===v.onCleanup(y,b,v))S=!1;else if(I(),e(s.add(c).add(u)).hide(),e(o.add(r)).unbind(),e(window).unbind("resize.fb scroll.fb"),e(document).unbind("keydown.fb"),o.find("iframe").attr("src",R&&/^https/i.test(window.location.href||"")?"javascript:void(false)":"about:blank"),"inside"!==v.titlePosition&&l.empty(),i.stop(),"elastic"==v.transitionOut){p=z();var n=i.position();h={top:n.top,left:n.left,width:i.width(),height:i.height()},v.opacity&&(h.opacity=1),l.empty().hide(),A.prop=1,e(A).animate({prop:0},{duration:v.speedOut,easing:v.easingOut,step:B,complete:t})}else i.fadeOut("none"==v.transitionOut?0:v.speedOut,t)},e.fancybox.resize=function(){r.is(":visible")&&r.css("height",e(document).height()),e.fancybox.center(!0)},e.fancybox.center=function(e){var t,n;S||(n=e===!0?1:0,t=M(),!n&&(i.width()>t[0]||i.height()>t[1])||i.stop().animate({top:parseInt(Math.max(t[3]-20,t[3]+.5*(t[1]-o.height()-40)-v.padding)),left:parseInt(Math.max(t[2]-20,t[2]+.5*(t[0]-o.width()-40)-v.padding))},"number"==typeof e?e:200))},e.fancybox.init=function(){e("#fancybox-wrap").length||(e("body").append(t=e('<div id="fancybox-tmp"></div>'),n=e('<div id="fancybox-loading"><div></div></div>'),r=e('<div id="fancybox-overlay"></div>'),i=e('<div id="fancybox-wrap"></div>')),a=e('<div id="fancybox-outer"></div>').append('<div class="fancybox-bg" id="fancybox-bg-n"></div><div class="fancybox-bg" id="fancybox-bg-ne"></div><div class="fancybox-bg" id="fancybox-bg-e"></div><div class="fancybox-bg" id="fancybox-bg-se"></div><div class="fancybox-bg" id="fancybox-bg-s"></div><div class="fancybox-bg" id="fancybox-bg-sw"></div><div class="fancybox-bg" id="fancybox-bg-w"></div><div class="fancybox-bg" id="fancybox-bg-nw"></div>').appendTo(i),a.append(o=e('<div id="fancybox-content"></div>'),s=e('<a id="fancybox-close"></a>'),l=e('<div id="fancybox-title"></div>'),c=e('<a href="javascript:;" id="fancybox-left"><span class="fancy-ico" id="fancybox-left-ico"></span></a>'),u=e('<a href="javascript:;" id="fancybox-right"><span class="fancy-ico" id="fancybox-right-ico"></span></a>')),s.click(e.fancybox.close),n.click(e.fancybox.cancel),c.click(function(t){t.preventDefault(),e.fancybox.prev()}),u.click(function(t){t.preventDefault(),e.fancybox.next()}),e.fn.mousewheel&&i.bind("mousewheel.fb",function(t,n){S?t.preventDefault():(0==e(t.target).get(0).clientHeight||e(t.target).get(0).scrollHeight===e(t.target).get(0).clientHeight)&&(t.preventDefault(),e.fancybox[n>0?"prev":"next"]())}),e.support.opacity||i.addClass("fancybox-ie"),R&&(n.addClass("fancybox-ie6"),i.addClass("fancybox-ie6"),e('<iframe id="fancybox-hide-sel-frame" src="'+(/^https/i.test(window.location.href||"")?"javascript:void(false)":"about:blank")+'" scrolling="no" border="0" frameborder="0" tabindex="-1"></iframe>').prependTo(a)))},e.fn.fancybox.defaults={padding:10,margin:40,opacity:!1,modal:!1,cyclic:!1,scrolling:"auto",width:560,height:340,autoScale:!0,autoDimensions:!0,centerOnScroll:!1,ajax:{},swf:{wmode:"transparent"},hideOnOverlayClick:!0,hideOnContentClick:!1,overlayShow:!0,overlayOpacity:.7,overlayColor:"#777",titleShow:!0,titlePosition:"float",titleFormat:null,titleFromAlt:!1,transitionIn:"fade",transitionOut:"fade",speedIn:300,speedOut:300,changeSpeed:300,changeFade:"fast",easingIn:"swing",easingOut:"swing",showCloseButton:!0,showNavArrows:!0,enableEscapeButton:!0,enableKeyboardNav:!0,onStart:function(){},onCancel:function(){},onComplete:function(){},onCleanup:function(){},onClosed:function(){},onError:function(){}},e(document).ready(function(){e.fancybox.init()})})(jQuery);
```
