## Supplementary figures and images for "Single individual structural variant detection uncovers widespread hemizygosity in molluscs"

### C1q_complete_phylogeny.pdf

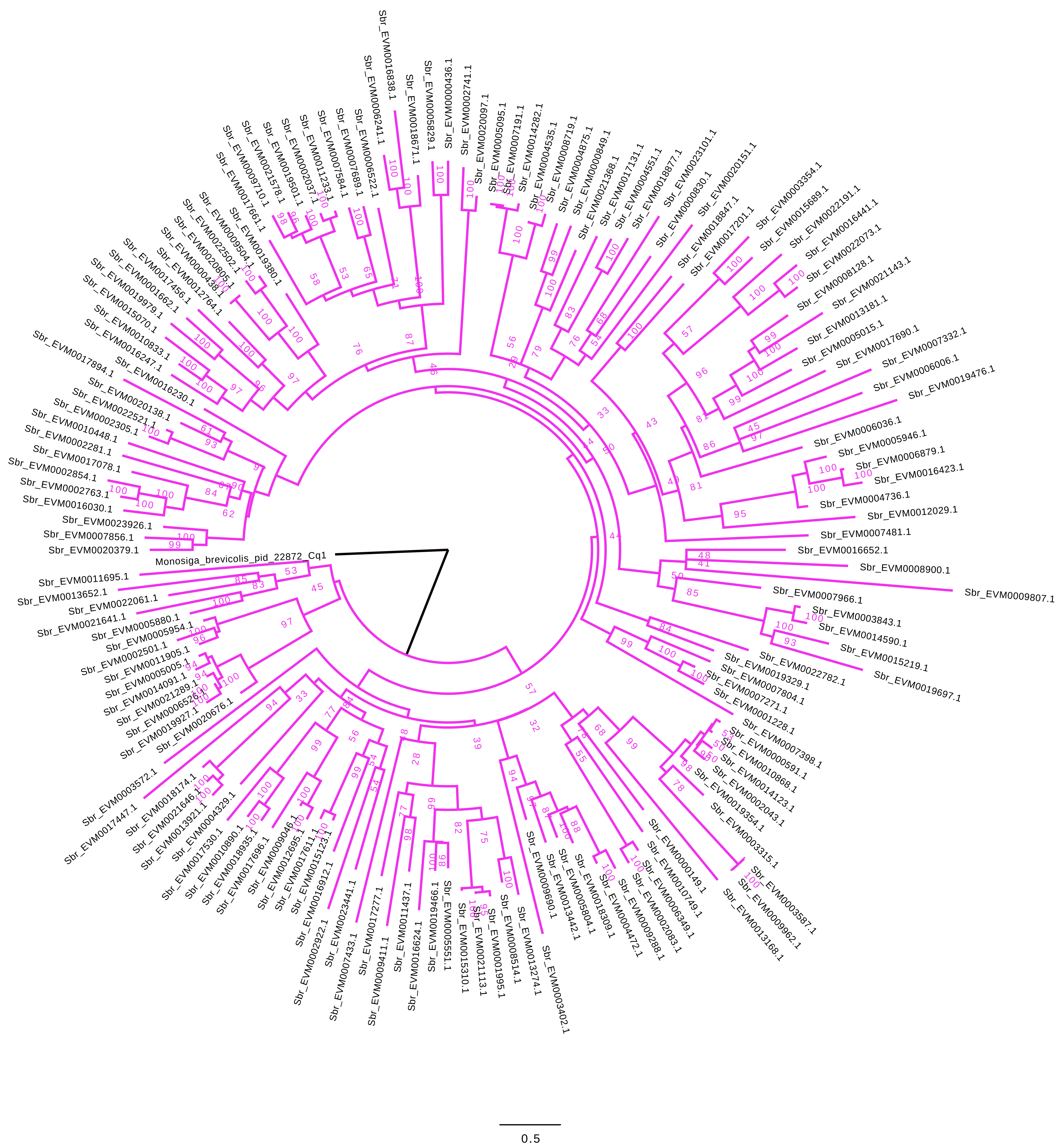

### HSP70_12_complete_phylogeny.pdf

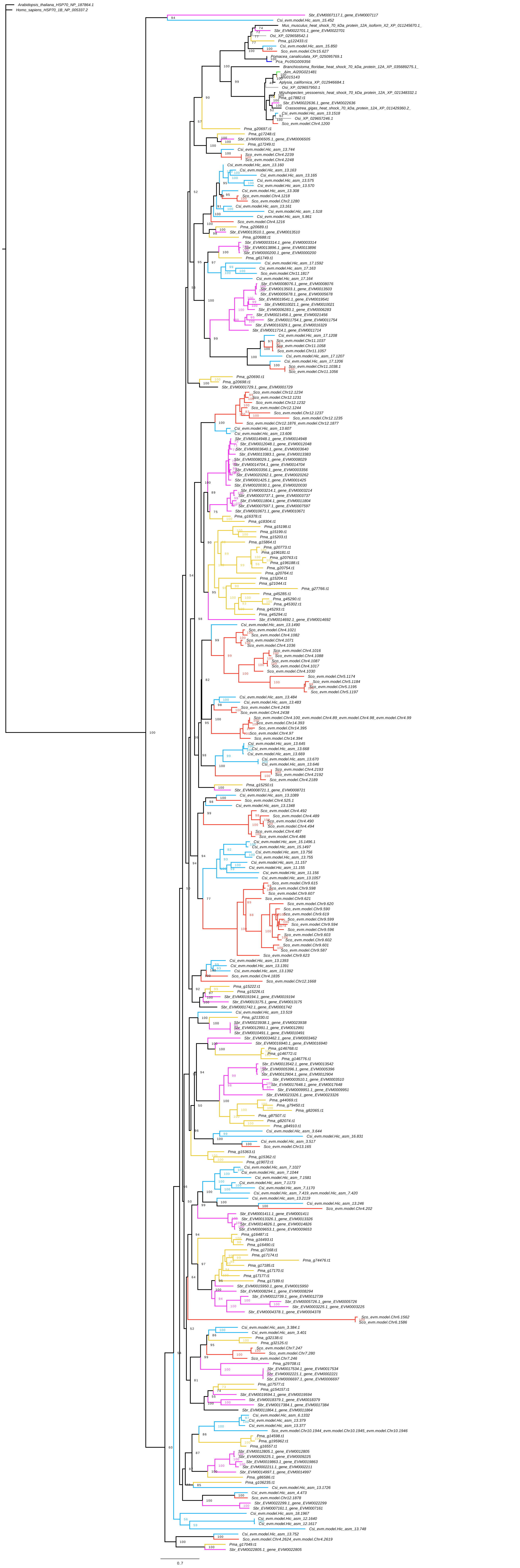

### HSP70_complete_phylogeny.pdf

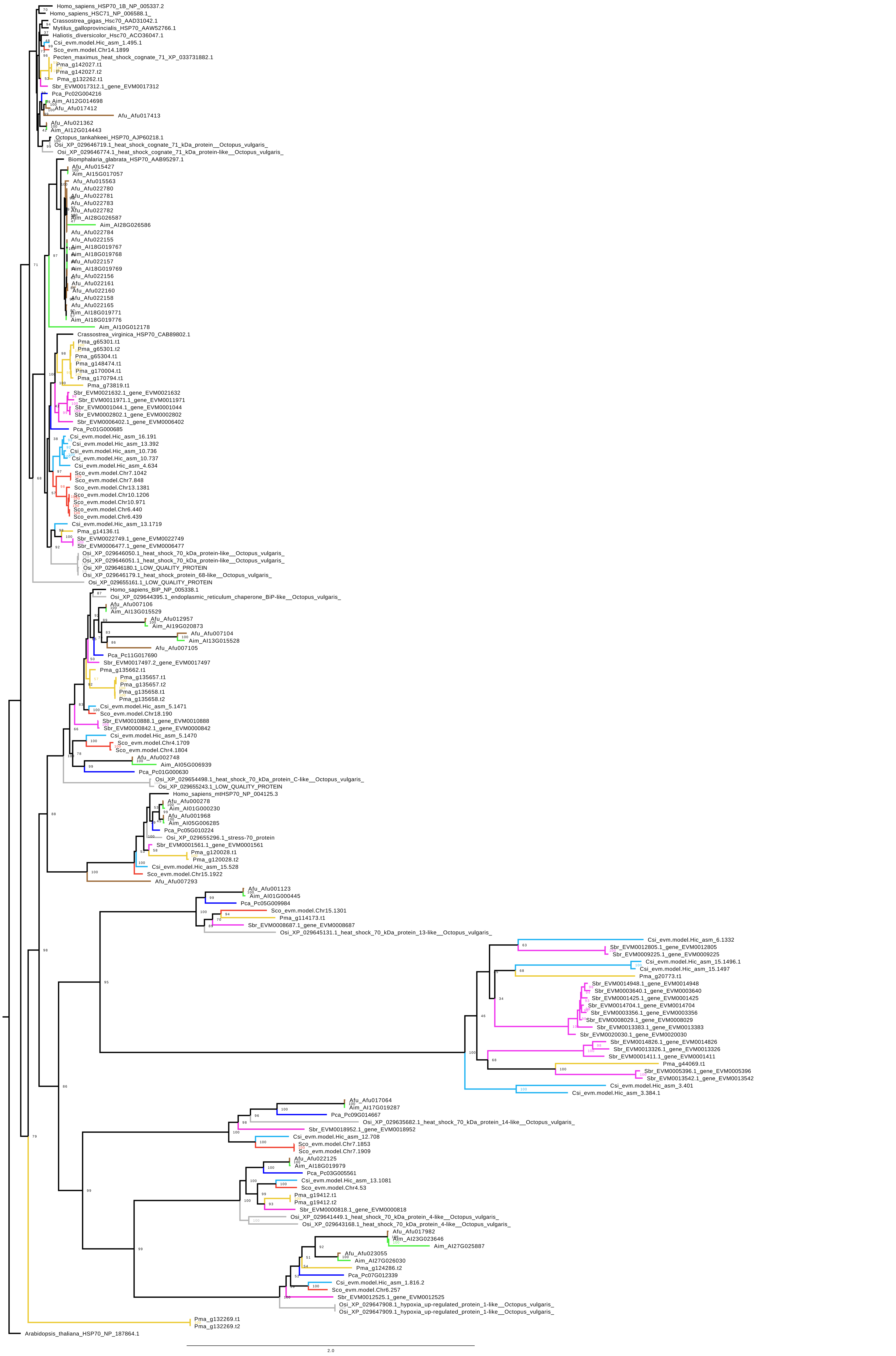

### plot.png

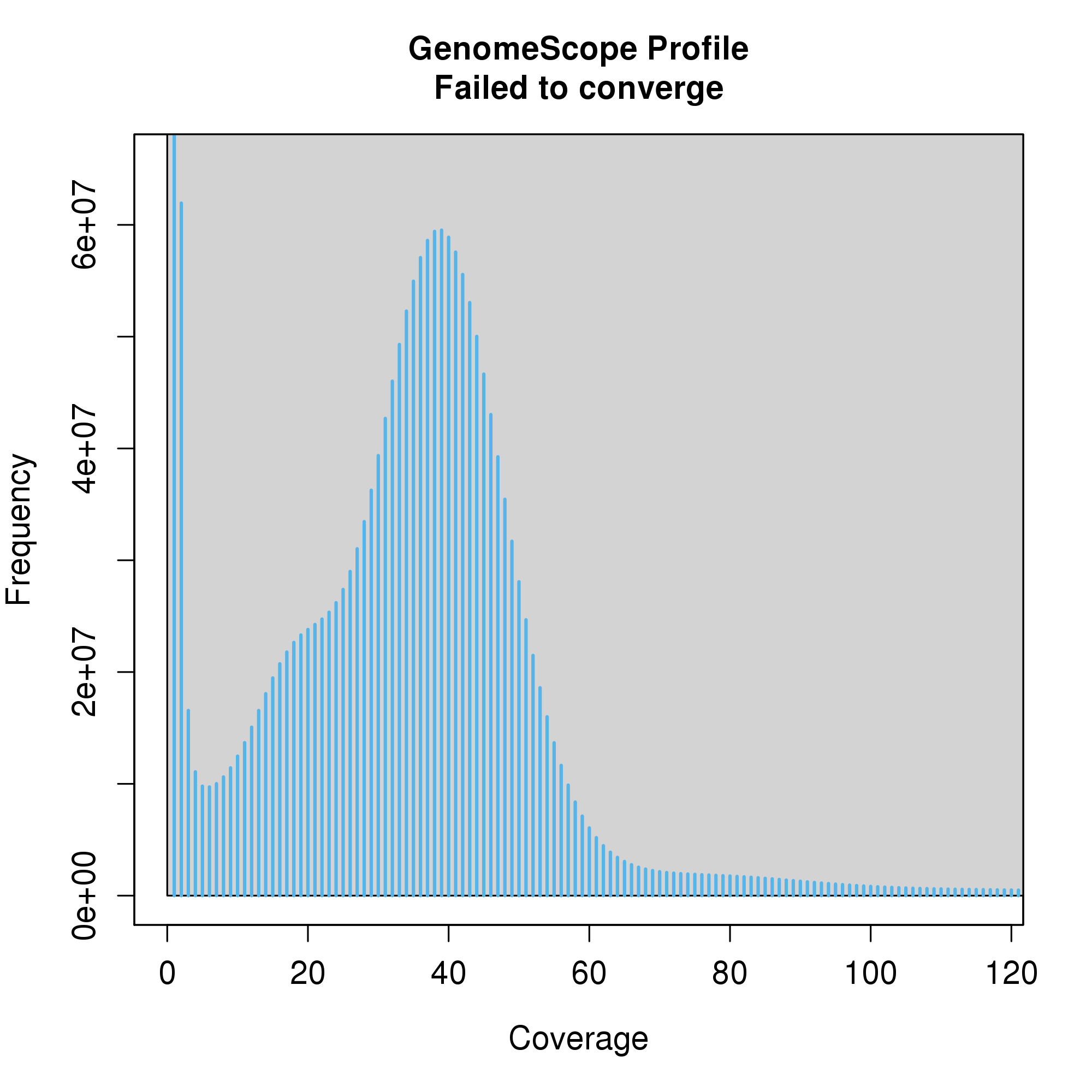

### plot_002.png

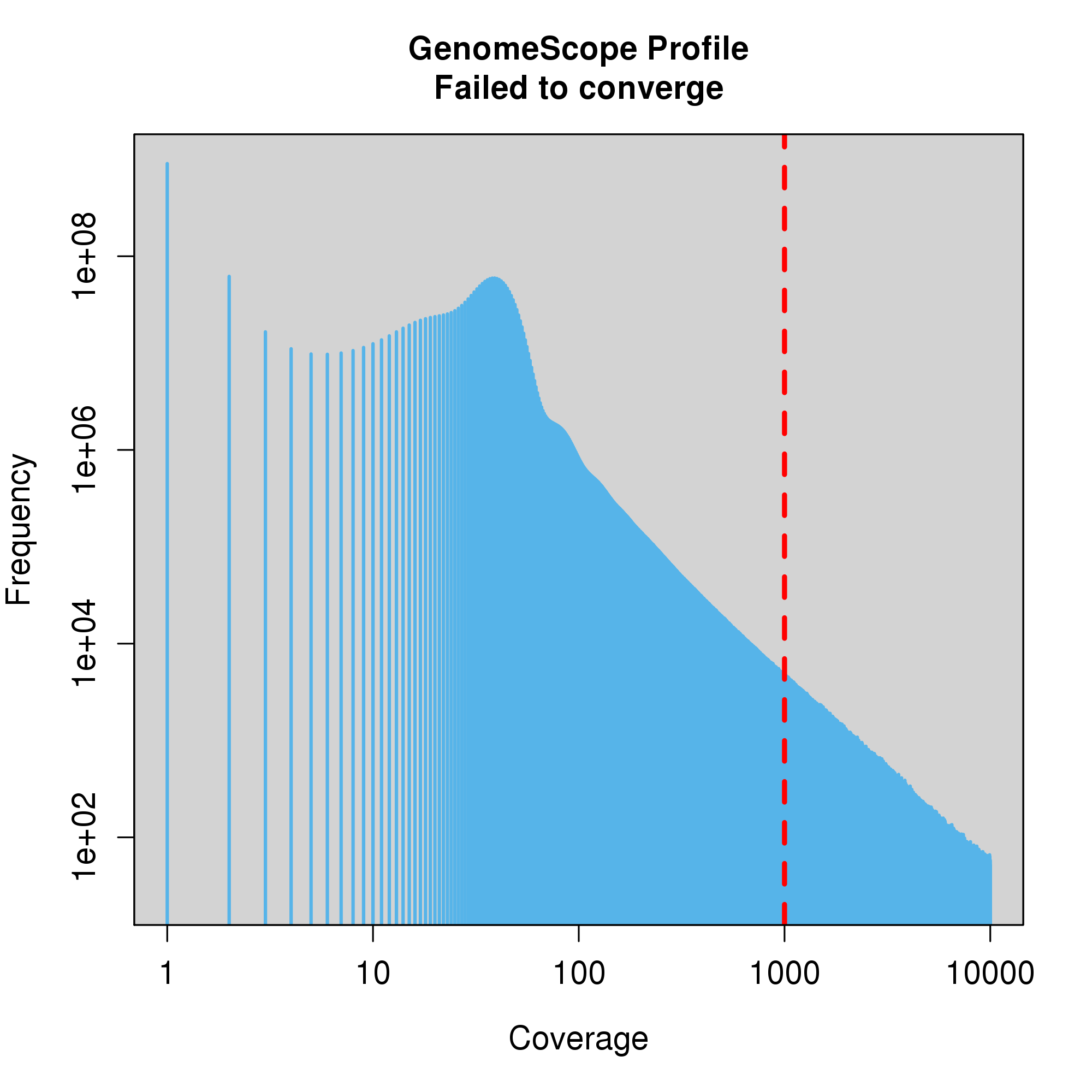
