## Supplementary Figures for "Single individual structural variant detection uncovers widespread hemizygosity in molluscs"

### Figure S1

Hypothetical transposition (birth) and gene loss (death) scenario

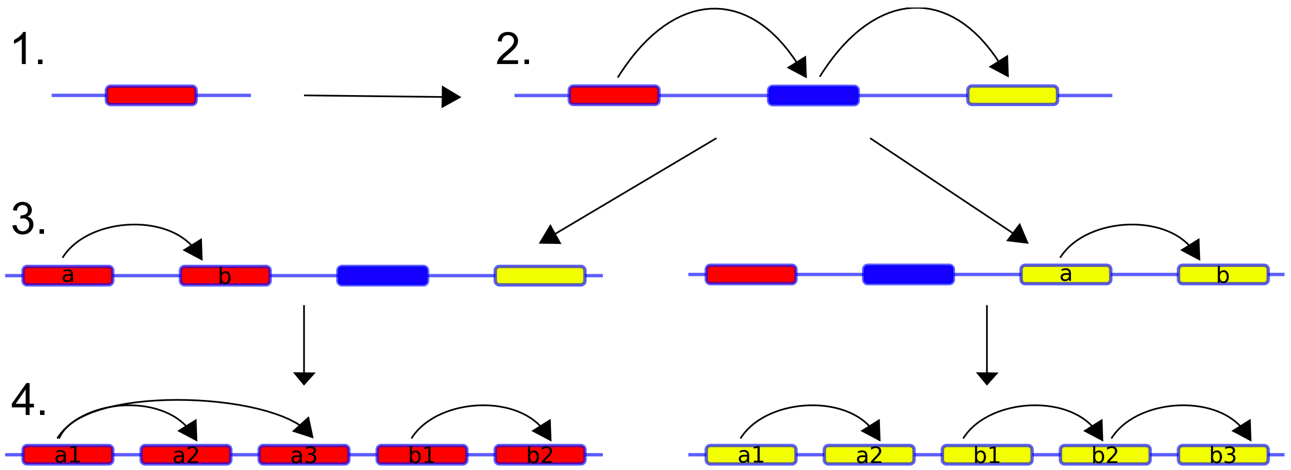


Resulting phylogenetic tree

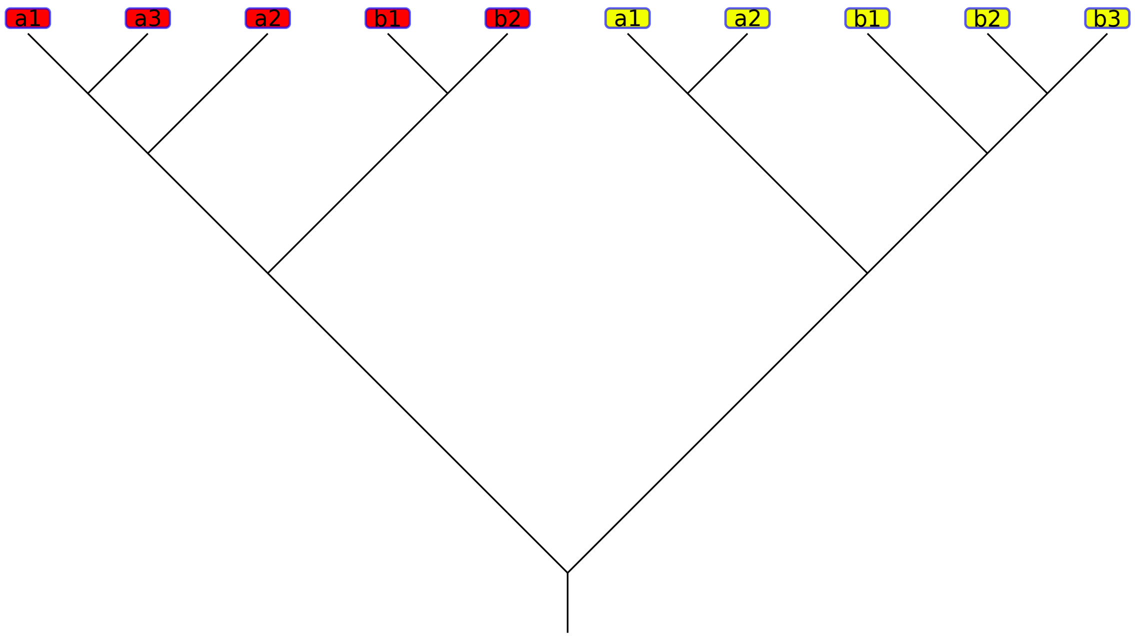

**Figure S2**

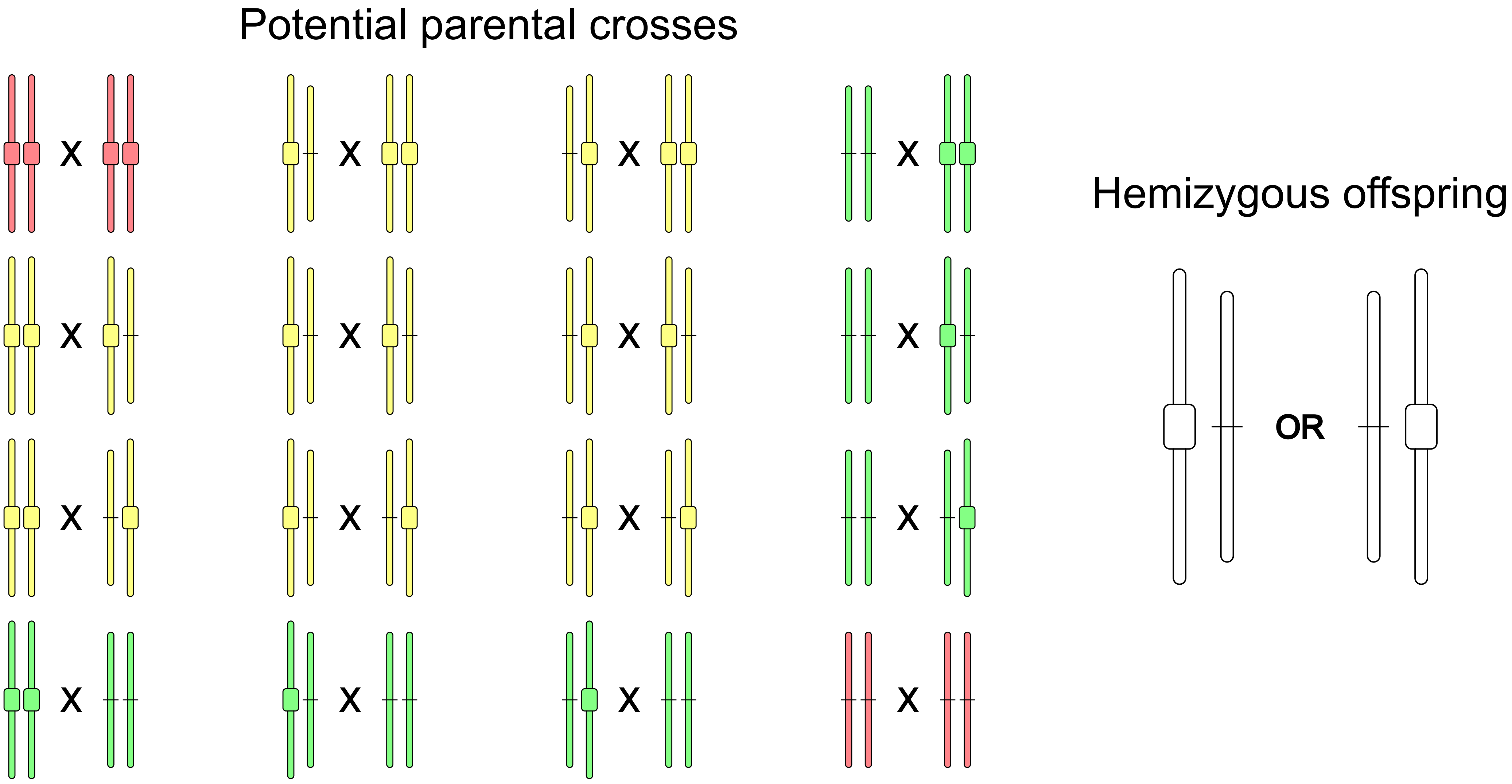

### Figure S3

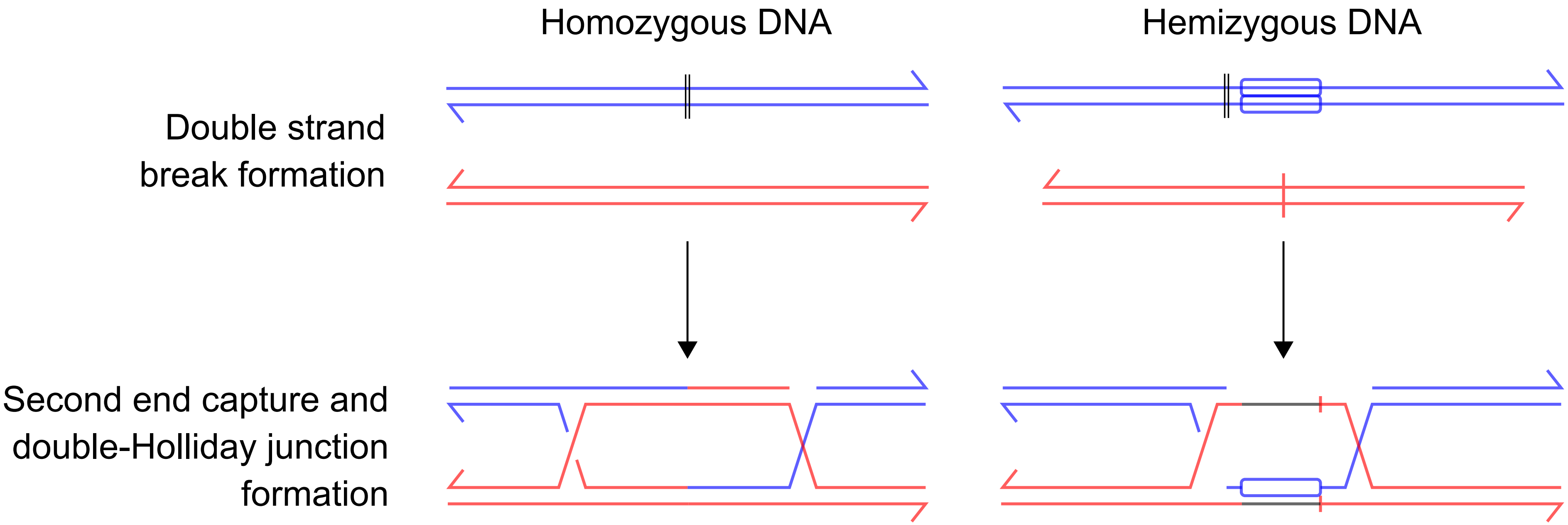
Homozygous DNA
Hemizygous DNA
Double strand
break formation
Second end capture and
double-Holliday junction
formation
